# Agomelatine drives sex-specific neuroprotection and reduced pathology in rat and human Alzheimer’s models

**DOI:** 10.64898/2026.01.17.700104

**Authors:** Grace A. Terry, Siaresh Aziz, Nishat Raihana, Lei Xie, Patricia Rockwell, Peter Serrano, Maria E. Figueiredo-Pereira

## Abstract

Alzheimer’s disease (AD) remains without effective disease-modifying therapies, underscoring the need for interventions that target interconnected molecular and cellular processes driving cognitive decline. Leveraging a cross-species translational framework integrating a progressive rat model of Alzheimer’s disease with human iPSC-derived neurons carrying familial AD mutations, we identify agomelatine (AGO) as a disease-modifying candidate. A clinically used melatonergic agonist and 5-HT_2C_ serotonergic antagonist, we found that AGO acts as a sex-selective modulator of AD-related neuronal and microglial dysfunction with therapeutic relevance across species.

TgF344-AD rats and their wildtype littermates received chronic dietary AGO (∼10 mg/kg/day) from 5 to 11 months of age and underwent hippocampal-dependent spatial learning assessment, quantitative hippocampal histopathology, and bulk RNA sequencing to evaluate the therapeutic effect on cognition, pathology, and molecular mechanisms. Human isogenic iPSC-derived cortical neurons carrying PSEN2^N141I^ or APPV^717I^ mutations were treated with 20 µM AGO followed by bulk RNA sequencing, to define AGO-driven transcriptional pathway modulation in AD neurons

In TgF344-AD rats, AGO produced robust female-specific benefits. AGO selectively restored hippocampal-dependant cognitive performance in female but not male transgenic rats. These improvements were independent of amyloid burden and instead aligned with reductions in microgliosis and pathogenic AT8-positive tau phosphorylation. Additionally, AGO normalized reactive and amoeboid microglial states exclusively in females and enhanced doublecortin-defined neurogenesis without altering mature NeuN⁺ neuronal density. This coordinated hippocampal stabilization highlights AGO’s capacity to restore plasticity rather than simply suppress pathology.

Transcriptomic analyses revealed sex-divergent mechanisms underlying these effects. In females, AGO activated metabolic, oxygen-handling, lipid-processing, neuroimmune, and CREB/IGF-1 signaling pathways while suppressing ER-stress, epigenetic, and ion-channel transcripts, changes consistent with resilience-promoting cellular reprogramming. In males, AGO preferentially modulated mitochondrial redox biology, transcriptional regulators, and extracellular matrix components. Despite these differences, both sexes showed AGO-induced engagement of conserved AD-relevant pathways, including shared induction of synaptic plasticity and hemoglobin/oxygen-transport related genes, suggesting a convergent neuroprotective molecular signature.

To translate these findings to a human system, we examined AGO’s effects in PSEN2^N141I^ and APP^V717I^ iPSC-derived cortical neurons. Both mutations produced convergent deficits in synaptic integrity, neuronal maturity, trophic signaling, proteostasis, metabolism, and excitability, alongside dysregulated developmental and ECM-remodeling programs. AGO partially reversed these pathogenic transcriptional changes, up-regulating synaptic, metabolic, vesicle-trafficking, and redox-stress resilience genes while suppressing pathological developmental and inflammatory pathways, demonstrating conserved engagement of neuronal recovery programs.

Together, these results identify AGO as a promising non-amyloid therapeutic candidate capable of modulating AD-relevant pathways in rodents and human models. The sex-selective efficacy observed *in vivo*, combined with conserved transcriptional responses across species, underscores the translational relevance of AGO-driven molecular reprogramming in AD.

## Introduction

Alzheimer’s disease (AD) is a progressive neurodegenerative disorder defined by extracellular amyloid-β (Aβ) plaques, intracellular neurofibrillary tangles, neuroinflammation, synaptic and neuronal loss leading to cognitive decline.^1^ These pathological processes contribute to widespread brain atrophy and are frequently accompanied by sleep disruption and depression, which further exacerbate cognitive impairment.^2,3^ Age remains the strongest risk factor for AD, yet mood disorders and circadian dysregulation have also been identified as significant contributors to disease vulnerability.^4^ Monoaminergic disruption, particularly within serotonergic pathways, is observed in both depression and AD, suggesting potentially convergent therapeutic targets.^5^

Despite decades of research, the precise mechanisms initiating and sustaining AD pathology remain unclear. Familial AD, caused by autosomal-dominant mutations in APP, PSEN1, or PSEN2, shares its core neuropathological features with the vastly more common sporadic form, though upstream molecular drivers likely differ.^6^ This limited mechanistic understanding has hindered therapeutic development. AD is currently the sixth leading cause of death in the United States, and unlike other major diseases, mortality from AD has risen sharply in recent decades, underscoring a persistent gap in treatment efficacy.^4^

Sex is an essential biological variable in AD. Approximately two-thirds of individuals with AD are women, and disease onset, symptom burden, and trajectory differ between sexes.^7^ While the mechanisms underlying this disparity remain incompletely understood, hormonal transitions, particularly during the perimenopausal period, are implicated.^8^ Estrogen supports synaptic plasticity, reduces tau phosphorylation, enhances neurotrophic signaling, and regulates neuronal metabolism. Abrupt declines in estrogen and changes in gonadal hormone balance during midlife may increase female vulnerability to neurodegeneration.^9,10^ These differences suggest that therapeutic responsiveness may vary by sex.

Available therapies provide only modest symptomatic benefit. Recently approved anti-amyloid monoclonal antibodies reduce Aβ burden and slow functional decline; however, they require intensive clinical monitoring, carry substantial risk of amyloid-related imaging abnormalities, and have not been shown to restore cognition or halt disease progression.^10^ Importantly, amyloid reduction does not consistently correlate with clinical improvement, reflecting the multifactorial nature of AD and the need for strategies that target additional pathological pathways.^11^

Drug repurposing, the strategy used in our current study, offers a powerful and efficient path to disease-modifying therapies for AD by capitalizing on established safety and pharmacokinetic profiles. Using a high-throughput in silico bioinformatics pipeline described in our previous publication^12^ and in^13^, we identified agomelatine (AGO) as a top candidate with predicted therapeutic potential in AD. AGO is a clinically approved, well-tolerated MT_1_/MT_2_ melatonergic agonist and 5-HT_2C_ serotonergic antagonist that readily penetrates the blood-brain barrier, providing a translationally advantageous therapeutic profile for targeting AD-relevant pathways.^14,15^ Originally developed as an antidepressant, AGO has demonstrated efficacy in mood and anxiety disorders and possesses antioxidant and neurotrophic properties relevant to AD.^16^ Its dual mechanism enables coordinated modulation of circadian, melatonergic, and serotonergic pathways, each implicated in AD pathogenesis. Importantly, 5-HT_2C_ antagonism increases cortical dopamine and norepinephrine availability while MT_1_/MT_2_ activation enhances synaptic plasticity, brain-derived neurotrophic factor (BDNF) signaling, and leads to the inhibition of GSK3β through activation of Wnt signaling.^17^ These pleiotropic actions suggest that AGO may correct multiple signaling disruptions underlying AD.

Melatonin signaling represents a biologically relevant pathway in AD.^18^ Melatonin levels decline with aging and more severely in AD, contributing to circadian disruption, increased oxidative stress, reduced BDNF expression, and impaired synaptic function.^19–21^ Activation of MT_1_ and MT_2_ receptors engages signaling cascades that enhance neuronal survival, inhibit GSK3β activity, promote BDNF-dependent TrkB signaling, and support memory consolidation.^22^ Thus, restoring melatonergic signaling may offer multifaceted neuroprotective effects.

Moreover, depression, a major risk factor and common comorbidity in AD, shares molecular features with AD, including impaired neurogenesis, reduced BDNF, chronic neuroinflammation, and monoaminergic dysfunction.^3^ These convergent mechanisms suggest that antidepressants with neurotrophic or anti-inflammatory actions may have benefit for AD-related pathology.

AD pathogenesis involves widespread dysregulation of interconnected signaling networks. Imbalanced amyloidogenic processing of APP increases Aβ production leading to downstream toxicity.^23^ GSK3β-mediated hyperphosphorylation of tau promotes neurofibrillary tangle formation and accelerates synaptic loss.^24^ Disruption of MAPK/ERK and PI3K/Akt/mTOR pathways undermines synaptic plasticity and memory formation, while reductions in BDNF further impair neuronal survival and neurogenesis.^1,25,26^ Mitochondrial dysfunction and oxidative stress emerge early in disease, contributing to bioenergetic failure and activation of apoptotic cascades.^27–29^ Chronic microglial activation becomes maladaptive, sustaining inflammation, and amplifying synaptic injury.^30,31^ Together, these alterations create a self-propagating cycle of neurodegeneration.

To investigate AGO’s therapeutic potential, we employed both the TgF344-AD rat model^32^, which recapitulates progressive Aβ deposition, tau pathology, neuroinflammation, synaptic and neuronal loss, and cognitive decline, as well as human isogenic iPSC-derived cortical neurons expressing familial AD mutations (PSEN2^N141I^ and APP^V717I^).^33^ This dual-system approach enables evaluation of AGO’s behavioural and neuropathological effects *in vivo*, while providing mechanistic insight into its transcriptional impact in a human based model. Across rodent and human models, agomelatine modulated synaptic, tau-related, and neuroimmune pathways with strongest effects in females, positioning it as a non-amyloid therapeutic candidate and underscoring sex as a critical variable for AD mechanism and translation.

## Materials and methods

### Animals

Male and female Fisher transgenic 344-AD (TgF344-AD) rats expressing human Swedish amyloid precursor protein (APPsw, KM670/671NL) and Δ exon 9 presenelin-1 (PS1ΔE9), and wild-type (WT) controls were obtained from the Rat Resource and Research Center, University of Missouri (RRID: RRRC_00699). Animals arrived at 4 weeks of age and were pair-housed under a 12-h light/dark cycle with ad libitum access to food and water. All animal procedures were performed in compliance with the relevant guidelines and regulations of the Institutional Animal Care and Use Committee (IACUC) at Hunter College. All experimental procedures were approved by the IACUC and agreed with all relevant standards and regulations outlined in the ARRIVE Essential 10 guidelines and the AVMA Guidelines for Euthanasia of Animals.^34^

### Agomelatine treatment

Agomelatine (AGO) was administered ad libitum in Purina 5001 rodent chow (Research Diets, Inc.) beginning at 5 months of age and maintained until 11 months of age. Treatment started early in the course of pathology when therapeutic intervention is predicted to be most effective. Chow was formulated to deliver approximately 10 mg/kg/day, based on previous studies.^35,36^ Untreated TgF344-AD and WT controls received non-drug chow. Weekly body weight and food consumption measurements were used to calculate AGO dosing (Supplementary Fig. 1). Rats were assigned to four experimental groups: WT untreated (WTNT, 7 male, 7 female), WT AGO-treated (WTTR, 4 male, 4 female), TgF344-AD untreated (TGNT, 5 male, 6 female), and TgF344-AD AGO-treated (TGTR, 6 male, 5 female), (Supplementary Table 1). The study timeline is provided in Fig. 1A.

**Figure 1.**
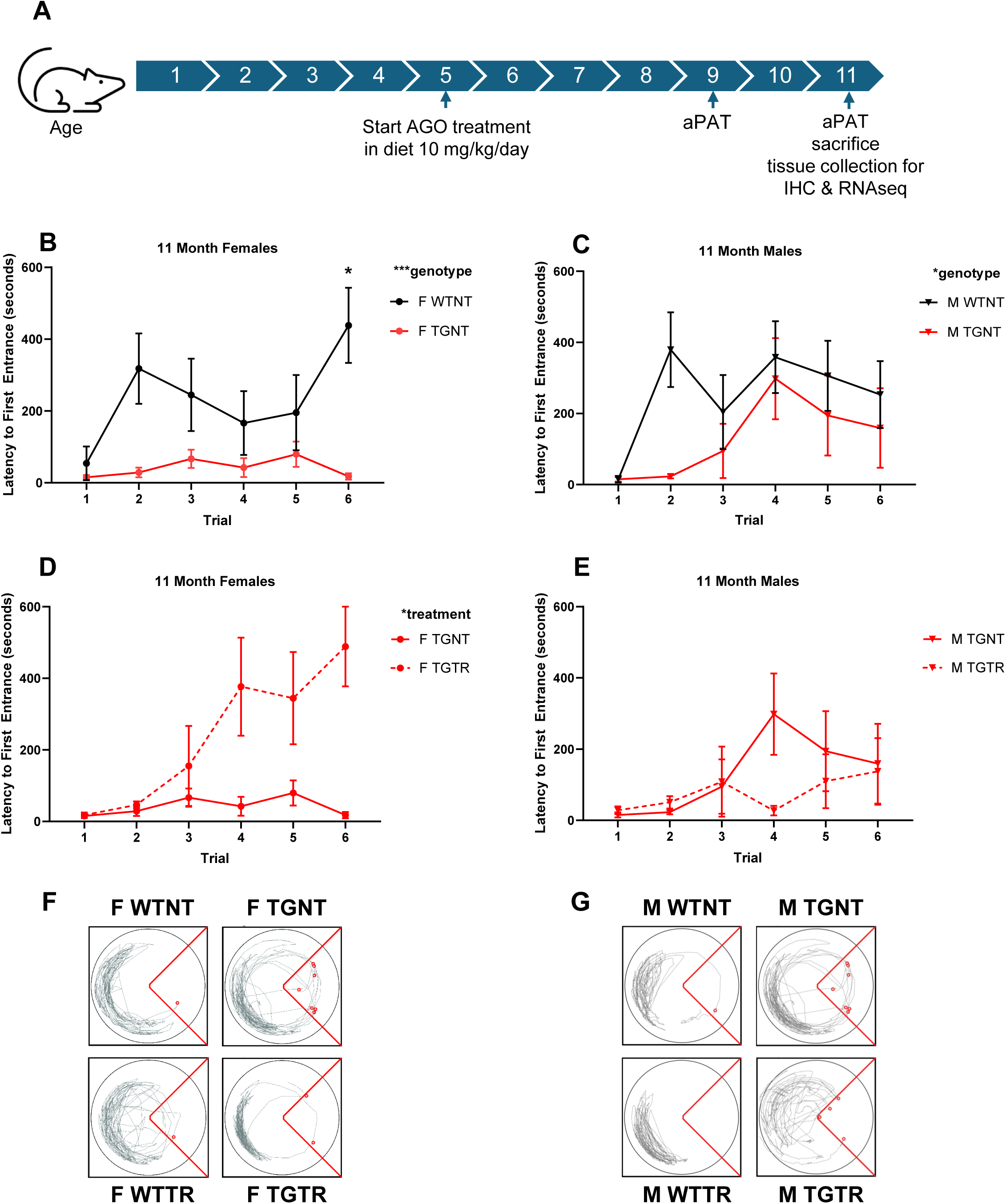
AGO mitigates spatial learning deficits in 11-month-old female but not male Tg-F344-AD rats. (**A**) Experimental design. All rats were exposed to the aPAT at 9 and 11 months of age. Rats were tasked with avoiding the shock zone over a course of six trials. (**B**) At 11 months of age, both female [*P* = 0.0371] (**C**) and male [*P* = 0.0009] TgF344-AD rats show impaired hippocampal dependent spatial learning in the active place avoidance task compared to WT littermates. (**D**) TGTR female rats perform significantly better than TGNT females [*P* = 0.0341] but (**E**) TGTR males do not perform significantly different from TGNT males [*P* = 0.7814]. (**F**) Traces from trial 3 showing female aPAT performance across groups. (**G**) Traces from trial 3 showing male aPAT performance across groups. All comparisons were run using three-way ANOVAs with Bonferroni post-hoc analyses. Error bars show the sample error of the mean and are only shown in one direction for visual clarity. aPAT: active place avoidance task, WTNT: wild-type not treated, WTTR: wild-type treated, TGNT: transgenic not treated, TGTR: transgenic treated, F: female, M: male. * <0.05, ** <0.01, ***<0.001, ****<0.0001.

### Active place avoidance task

Spatial learning and hippocampal-dependent memory were assessed at 9 and 11 months of age using the active place avoidance task (aPAT).^37^ Rats were tested in a rotating circular arena containing an unmarked, fixed shock zone. Entry into the zone triggered a 0.2 amp foot shock every 1.5 s until exit. Extra-maze visual cues enabled rats to maintain spatial orientation as the arena rotated at 1 rpm. Behavior was recorded using an overhead camera and analyzed with Trackpad software. Among several parameters in aPAT, we chose to compare latency to first entrance into the shock zone, a commonly used hippocampal-dependent index of spatial learning and initial recall of the shock-associated location. Rats were singly housed for 30 min before testing. A 10 min habituation trial (shock off) was followed by six 10 min training trials separated by 10 min intervals. A 10 min probe trial without shock was performed 24 h later. Shock zone location was changed between the two testing ages.

### Tissue collection

At 11 months, rats were anesthetized with ketamine/xylazine (100/10 mg/kg, i.p.) and perfused with ice-cold RNase-free PBS. Brains were removed and hemisected. The right hemisphere was microdissected (prefrontal, cingulate, entorhinal cortices; hippocampus) and snap-frozen for RNA sequencing. The left hemisphere was post-fixed in 4% paraformaldehyde for 48 h at 4°C, cryoprotected in 30% sucrose, flash-frozen in 2-methylbutane, and stored at −80°C. Coronal sections (30 µm) were cut using a Leica CM3050S cryostat and stored in cryoprotectant at −20°C.

### Immunohistochemistry

IHC was performed on dorsal hippocampal (left hemisphere) sections (−3.36 to −4.40 mm bregma), as described in^38^. Sections were mounted on gelatin-coated slides, washed in 0.3% Triton-X/PBS (T-PBS), and incubated with 0.05 M glycine to reduce autofluorescence. Blocking was performed with 15% normal goat serum (NGS), followed by overnight incubation at 4°C with primary antibodies [Aβ 4G8, Iba1, AT8, NeuN, doublecortin (DCX); Supplementary Table 2]. Alexa Fluor secondary antibodies (1:250, Supplementary Table 2) were applied for 1 h. Sections were washed and coverslipped with DAPI-containing medium.

Images were acquired at 10× using a Zeiss AxioImager M2 microscope equipped with AxioCam MRm Rev. 3 and a motorized stage. Exposure settings were held constant. Whole-hippocampus mosaics were generated and saved as ZVI files.

### IHC quantification

ZVI images were converted to TIFF formats in FIJI.^39^ Batch-processing scripts developed in our lab^12^ were used to quantify positive signal based on pixel-intensity thresholds. Background subtraction was performed using the rolling-ball method.^40^ Positive signals from each image were extracted using the average pixel intensity + (modifier[1(Iba-1) or 2(Abeta, AT8, NeuN, DCX)] x standard deviation). Two hippocampal sections per rat were analyzed.

Quantification included the whole left dorsal hippocampus and its subregions cornu ammonis 1 (CA1), cornu ammonis 3 (CA3), dentate gyrus (DG), and subiculum (SB). DCX analyses focused on the subgranular zone. Microglial morphology was classified using form factor (FF = 4π*area/perimeter²): ramified (0–0.499), reactive (0.5–0.699), and amoeboid (≥0.70).^41,42^ Particle size was set to 50-infinity. Analyses were performed using GraphPad Prism 10.5.

### Hippocampal RNA sequencing

Bulk RNA sequencing was performed at the UCLA Technology Center for Genomics and Bioinformatics (Illumina NovaSeq6000; 2×50 bp). Hippocampal (right hemisphere) samples (*n* = 5 per group: TGNT male, TGTR male, TGNT female, TGTR female) were analyzed. RNA was extracted and libraries were prepared using the KAPA mRNA-Seq Hyper Prep Kit. Reads were quality-checked using Illumina SAV, demultiplexed with bcl2fastq v2.19.1.403, aligned to the rat Rn6 genome with STAR v2.7.9a, and quantified using STAR’s built-in gene counting.^43^

### DESeq2 analysis

Differentially expressed genes (DEGs) were identified using DESeq2 in RStudio.^44,45^ Significance criteria were *P* < 0.05 and |log₂ fold-change| > 1. Gene ontology analyses (biological processes, molecular functions, cellular components) were conducted using Bioconductor and clusterProfiler.^46,47^ Male and female analyses were carried out separately.

### iPSC culture

Human male iPSC lines (WT KOLF2.1J, cat # JIPSC001000; PSEN2^N141I^ SNV/SNV, cat # JIPSC001050; APP^V717I^ SNV/SNV, cat # JIPSC001202; from Jackson Laboratory^48^) were thawed onto Synthemax-coated plates at 5×10⁵ cells/well in StemFlex medium with RevitaCell. After 24 h, RevitaCell was removed and media were changed every other day. Cells were passaged at ∼80% confluence with ReLeSR. After two passages, cells were transitioned to Essential 8 medium and maintained with daily media changes at 37°C, 5% CO₂.

### NGN2 transfection

Neuronal induction was initiated using the PiggyBac transposon system. Cells were dissociated with Accutase and replated in CEPT-supplemented medium at 1.5×10⁶ cells/well.^49^ One hour later, a 1 μg to 3 μg ratio of Super piggyBac Transposase expression vector (System Biosciences cat#PB210PA-1) to PB-hNGN2 plasmid (Addgene cat#198397, developed by Dr. Micheal Ward) in lipofectamine solution was added dropwise and incubated overnight.^50^ Puromycin selection was applied (0.5 mg/mL for 2 days, then 1.0 mg/mL for 2 days). Transfected cultures were expanded and cryopreserved.

### NGN2-directed cortical neuron differentiation

Differentiation proceeded in two phases: induction and maturation.^51,52^ For induction, NGN2-transfected cells were plated at 1×10⁵ cells/well on Matrigel and treated with doxycycline (2 µg/mL) for 3 days in KnockOut DMEM/F12 with N2, NEAA, GlutaMAX, and RevitaCell (first 24 h). Precursors were replated on day 3 onto dPGA/laminin-coated 96-well plates (1.5×10⁴ cells/well) and matured in a 50% DMEM/F12 + 50% BrainPhys medium supplemented with N21 Max, laminin, BDNF, GDNF, and NT-3. Doxycycline, 5-fluoro-2′-deoxyuridine, and uridine were included until day 10. Media were changed twice weekly. Cells were harvested at day 35.

### AGO treatment of iPSC-derived neurons

AGO was dissolved in DMSO and applied at 20 µM beginning on day 14 (early) or day 21 (late). Final DMSO concentration was 0.1% in all wells. Controls received 0.1% DMSO. Treatments were maintained until day 35 when neurons are considered mature.^51^

### RNA sequencing of iPSC-derived neurons

RNA sequencing was performed by Alithea Genomics (Epalinges, Switzerland) using DRUG-seq (BRB-seq). Samples were sequenced at 5 million reads per sample (≥ 3 biological replicates/group). Differentially expressed genes (DEGs) were identified using DESeq2^44,45^ (*P* < 0.05; |log₂ fold-change| > 1), and gene ontology (GO) enrichment was performed using Bioconductor^46^ and clusterProfiler.^53^ Volcano plots were generated using Qiagen IPA program.

### Statistical analyses

For every IHC data set we ran a PRISM ROUT outlier test set at Q =1%. There were no outliers detected. Behavioral analyses used two- or three-way repeated-measures ANOVA with Bonferroni post hoc tests. IHC analyses used two-way ANOVA (Aβ) or three-way ANOVA (Iba1 morphology, AT8, DCX, NeuN) followed by Sidak post hoc tests. The alpha level was set at *P* < 0.05 with a 95% confidence interval for each effect. GraphPad’s Prism version 8 (La Jolla, CA, USA) was used.

### Data availability

All data supporting this manuscript are available upon reasonable request. RNA-seq datasets will be deposited in the Gene Expression Omnibus (GEO) upon acceptance, with accession numbers provided in the final manuscript. Behavioral, imaging, and analysis scripts are available from the corresponding author.

## Results

### AGO reduced spatial learning deficits selectively in female TgF344-AD rats

Cognitive impairment is a defining feature of AD and contributes substantially to diminished quality of life. Thus, improvement in cognitive performance is essential for therapeutic success. To assess AGO’s effect on spatial learning we used the aPAT at both 9 and 11 months of age. As we previously reported^54^, at 9 months of age, there were no differences in spatial learning (latency to first entrance) between WT and TgF344-AD rats in males or females [male genotype effect: *F*(1,15) = 1.375, *P* = 0.2592; female genotype effect: *F*(1,20) = 3.025, *P* = 0.0973] nor were any significant treatment effects observed [male treatment effect: *F*(1,15) = 0.7502, *P* = 0.4001; female treatment effect: *F*(1,20) = 0.02369, *P* = 0.8792; Supplementary Fig. 2, Supplementary Table 3]. As shown in Fig. 1B-C, at 11 months of age, TgF344-AD rats exhibited significantly impaired spatial learning compared to WT rats in both males and females [male genotype effect: *F*(1,18) = 15.85, *P* = 0.0009; female genotype effect: *F*(1,18) = 5.069, *P* = 0.0371], supporting this model’s development of age-dependant cognitive decline. In female rats, AGO treatment improved spatial learning at 11 months of age [treatment effect: *F*(1,18) = 5.258, *P* = 0.0341; Fig. 1D, Supplementary Table 4]. In contrast, AGO did not produce comparable benefits in males with no treatment effect observed [treatment effect: *F*(1,18) = 0.07933, *P* = 0.7814; Fig. 1E, Supplementary Table 4].

Three-way ANOVA revealed no effects of sex or treatment on cognition in across all wildtype groups (male WTNT, male WTTR, female WTNT, female WTTR) at 11-months [treatment: F(1,18) = 0.2745, P = 0.6067; sex: F(1,18) = 0.1139, P = 0.7396; Supplementary Table 4]. In contrast, analysis across transgenic rats (male TGNT, male TGTR, female TGNT, female TGTR) exhibited a significant treatment effect and a strong treatment × sex interaction, without a main effect of sex [treatment: F(1,18) = 4.438, P = 0.0494; sex: F(1,18) = 1.139, P = 0.3000; treatment × sex: F(1,18) = 13.62, P = 0.0017], indicating comparable baseline cognition between sexes but a preferential cognitive benefit of AGO in females. Figures 1F and 1G show the traces for individual rats during Trial 3 across the four treatment conditions.

These findings show that spatial learning deficits are not yet present at 9 months of age but clearly emerge in TgF344-AD rats by 11 months of age, and that AGO selectively reduces these deficits in females, with no comparable effect in males. Full statistical results are provided in Supplementary Tables 3-4.

### AGO did not alter hippocampal Aβ plaque burden in TgF344-AD rats

We quantified Aβ plaque burden within the whole left dorsal hippocampus and its four subregions (CA1, CA3, DG and SB), to determine whether AGO treatment modulated histopathology (Fig. 2A-D). Across both sexes, AGO administration did not alter Aβ plaque accumulation in the left hippocampal region [treatment effect: *F*(1,17) = 0.1601, *P* = 0.6941; Fig. 2E] or within the DG [treatment effect: *F*(1,17) = 0.9629, *P* = 0.3402; Fig. 2F]. Furthermore, no sex-dependent differences in Aβ burden were observed between male and female untreated TgF344-AD rats, indicating comparable baseline amyloid pathology across sexes [hippocampus: *F*(1,17) = 2.024, *P* = 0.1729; DG: *F*(1,17) = 1.687, *P* = 0.2113; Fig. 2E-F, Supplementary Table 5].

**Figure 2.**
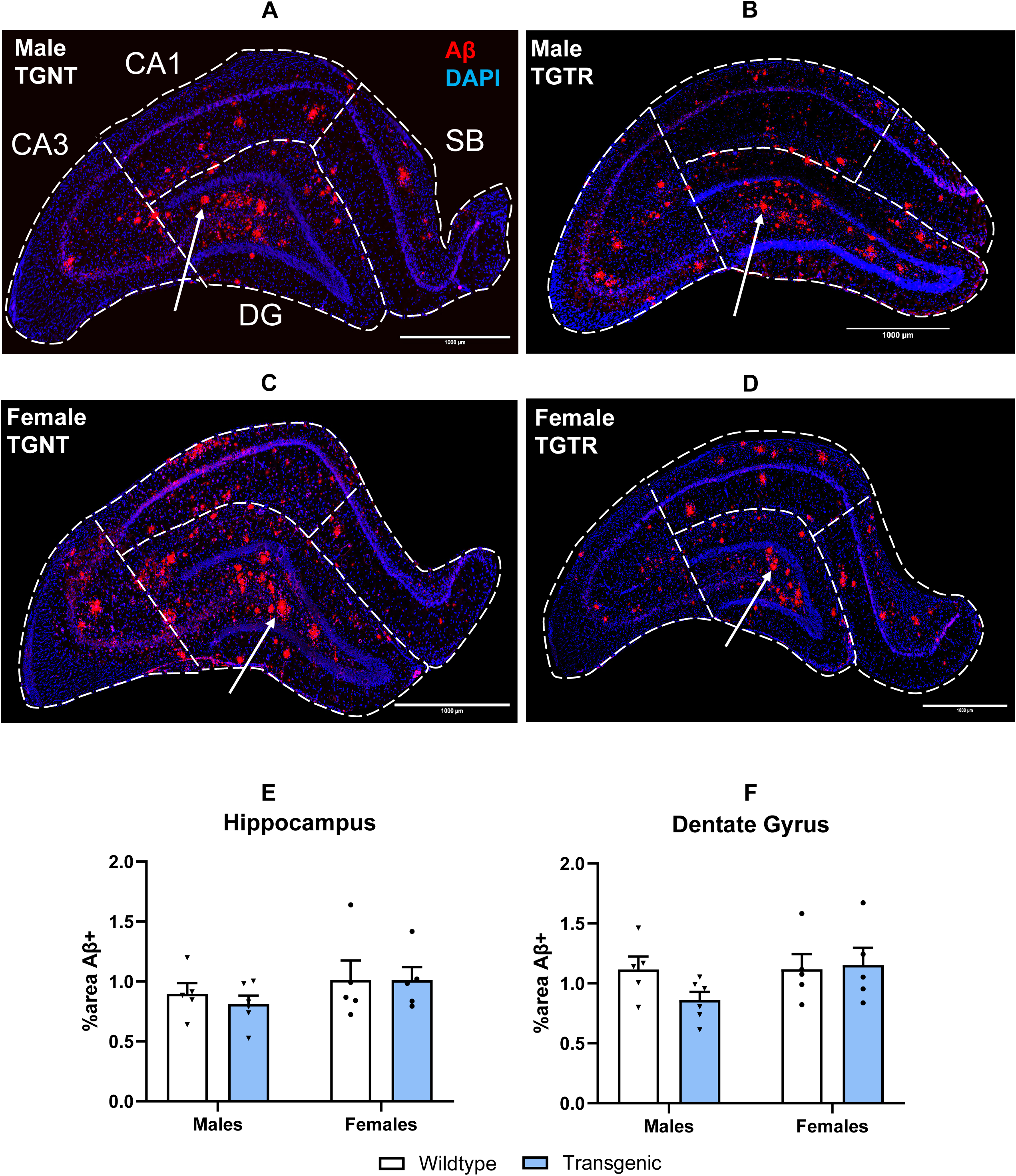
AGO does not alter hippocampal Aβ plaque burden in TgF344-AD rats. (**A–D**) Representative Aβ immunohistochemistry in the dorsal hippocampus, with subregions indicated. (**E–F**) Quantitative analysis revealed no effect of AGO on Aβ plaque burden across the dorsal hippocampus (*P* = 0.6941) or within the dentate gyrus (DG; *P* = 0.3402). Aβ immunoreactivity (red) was used to visualize plaque deposition, and nuclei were counterstained with DAPI (blue). Scale bar, 1000 µm. Data were analyzed by two-way ANOVA. N = 5–6 per group. CA1, Cornu Ammonis 1; CA3, Cornu Ammonis 3; DG, dentate gyrus; SB, subiculum; IHC, immunohistochemistry; TGNT, transgenic not treated; TGTR, transgenic treated.

These findings indicate that AGO does not modify Aβ plaque burden in the hippocampus or its subregions, with comparable baseline amyloid pathology observed across sexes. Full statistical results are provided in Supplementary Table 5.

### AGO attenuated microgliosis in female but not male TgF344-AD rats

To assess microglial activation, we quantified Iba1 immunoreactivity (Fig. 3A-H), a pan-microglial marker that labels most microglia subpopulations and macrophages, and is upregulated when microglia/macrophages become activated. We classified microglia as ramified, reactive, or amoeboid based on form-factor circularity.^38^ The percentage area of Iba1⁺ signal was measured in the left dorsal hippocampus and its four subregions (CA1, CA3, DG, SB) in all groups.

**Figure 3.**
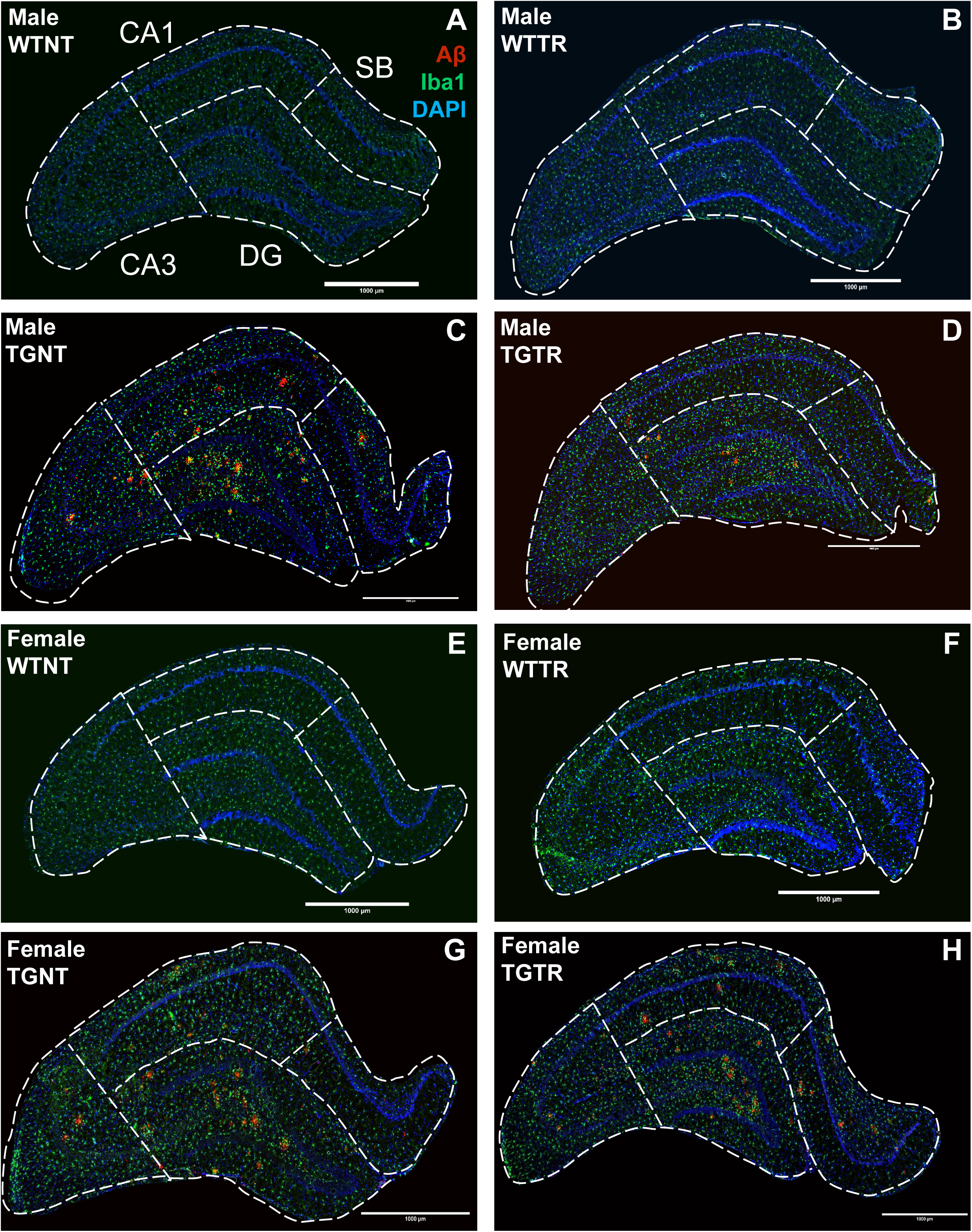
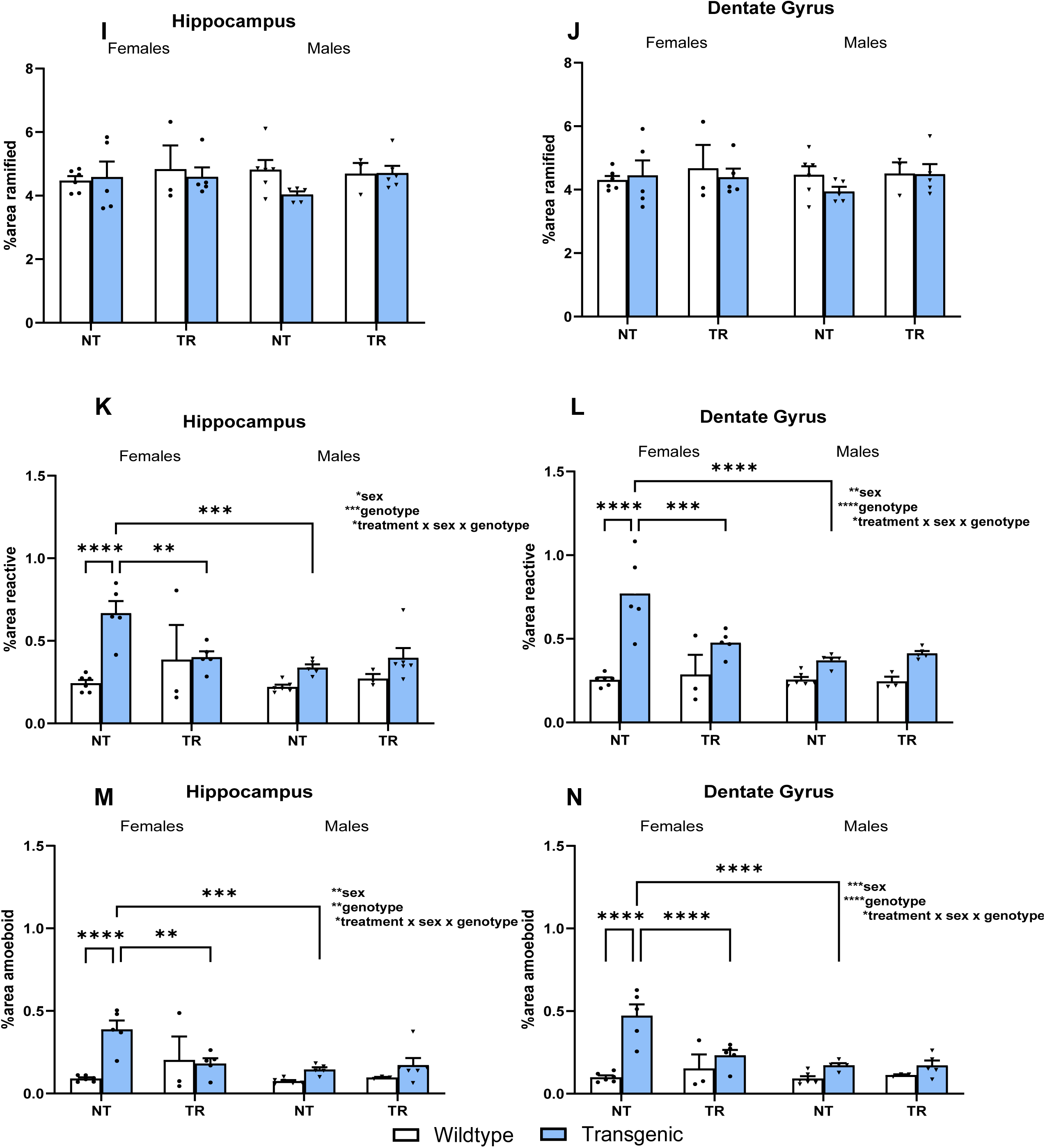
Sex-specific normalization of reactive and amoeboid microglia by AGO in TgF344-AD rats. (**A–H**) Representative images of the dorsal hippocampus stained for Iba1 (green) to label microglia, Aβ (4G8; red) to visualize plaques, and DAPI (blue) to label nuclei, with hippocampal subregions indicated. (**I–J**) Quantification of ramified microglia revealed no significant effects of genotype or AGO treatment in either the hippocampus (genotype: *P* = 0.3469, treatment: *P* = 0.3351) or DG (genotype: *P* = 0.4832, treatment: *P* = 0.3514). (**K–L**) Reactive microglia were significantly increased in TgF344-AD rats relative to WT in both the hippocampus (genotype: *P* = 0.0005) and DG (genotype: *P* < 0.0001); notably, AGO selectively reversed this increase in females, but not males, in both regions: hippocampus (female TGTR vs TGNT post-hoc: *P* = 0.038], and DG (female TGTR vs TGNT post-hoc: *P* = 0.0025). (**M–N**) Amoeboid microglia similarly accumulated in TgF344-AD rats in the hippocampus (genotype: *P* = 0.0017) and DG (genotype: *P* < 0.0001). This pathological shift was selectively normalized by AGO in females only, in the hippocampus (female TGTR vs TGNT post-hoc: *P* = 0.0154) and DG (female TGTR vs TGNT post-hoc: *P* = 0.0006). Scale bar, 1000 µm. Data were analyzed using three-way ANOVA with Sidak’s post hoc tests. AGO, agomelatine; WTNT, wild-type not treated; WTTR, wild-type treated; TGNT, transgenic not treated; TGTR, transgenic treated; CA1, Cornu Ammonis 1; CA3, Cornu Ammonis 3; DG, dentate gyrus; SB, subiculum; F, female; M, male, G*: genotype effect, t*: treatment effect. *P < 0.05, **P < 0.01, ***P < 0.001, ****P < 0.0001.

At baseline, TGNT rats showed robust microgliosis, with significantly elevated reactive and amoeboid microglia relative to WTNT controls [Fig. 3K–N, hippocampus reactive: *F*(1,31) = 15.09, *P* = 0.0005; hippocampus amoeboid: *F*(1,31) = 11.81, *P* = 0.0017; DG reactive: *F*(1,30) = 44.88, *P* < 0.0001; DG amoeboid: *F*(1,30) = 30.45, *P* < 0.0001]. In contrast, ramified microglia did not differ for either sex [Fig. 3I-J, hippocampus: *F*(1,31) = 0.9123, *P* = 0.3469; DG: *F*(1,30) = 0.5040, *P* = 0.4832].

AGO produced sex-specific effects. In females, TGTR rats showed significant reductions in reactive and amoeboid microglia compared with TGNT [Fig. 3K–N, post-hoc analysis between female TGNT and TGTR groups: hippocampus reactive: *P* = 0.038; hippocampus amoeboid: *P* = 0.0154; DG reactive: *P* = 0.0025; DG amoeboid: *P* = 0.0006], preserving levels to WTNT values. In males, AGO had no effect [Fig. 3K–N, post-hoc analysis between male TGNT and TGTR groups hippocampus reactive: *P* = 0.9995; hippocampus amoeboid: *P* > 0.9999; DG reactive: *P* > 0.9999; DG amoeboid: *P* > 0.9999].

Male TGNT rats exhibited significantly lower reactive and amoeboid microglial burden than TGNT females [hippocampus reactive: *P* = 0.0052; hippocampus amoeboid: *P* = 0.0029; DG reactive: *P* < 0.0001; DG amoeboid: *P* < 0.0001], indicating a baseline sex difference.

Overall, these findings show that AGO attenuates microgliosis in a sex-dependent manner, normalizing reactive and amoeboid microglia in females but not males, and that at baseline female TgF344-AD rats exhibit higher levels of reactive and amoeboid microglia than males. Full statistical results are provided in Supplementary Table 6.

### AGO decreased pathogenic AT8-positive tau hyperphosphorylation in TgF344-AD rats

Hippocampal tau pathology was quantified using AT8, a well-established marker of pathological, triple-phosphorylated PHF-tau (pS202/pT205/pS208) associated with AD (Fig. 4A–H). AGO produced a significant overall reduction in hippocampal AT8 immunoreactivity [main treatment effect: *F*(1,31) = 5.815, *P* = 0.0249; Fig. 4I], indicating attenuation of pathogenic tau phosphorylation. Interestingly, this effect was sex- and subregion-specific: in the DG, as AGO selectively reduced AT8 density in females but not males [three-way ANOVA Treatment × Sex interaction: *F*(1,31) = 4.822, *P* = 0.0357; two-way ANOVA males: *F*(1,16) = 0.1750, *P* = 0.6813; two-way ANOVA females: *F*(1,15) = 7.524, *P* = 0.0151; Fig. 4J]. Baseline AT8 levels did not differ between untreated males and females in either the hippocampus or DG, indicating comparable tau pathology across sexes prior to treatment [hippocampus: *F*(1,31) = 0.2097, *P* = 0.6502; dentate gyrus: *F*(1,31) = 0.1665, *P* = 0.6860]. Collectively, these data demonstrate that AGO suppresses hippocampal tau hyperphosphorylation, with a female-specific reduction in the DG and no detectable effects in the remaining hippocampal subregions. Full statistical analyses are provided in Supplementary Table 7.

**Figure 4.**
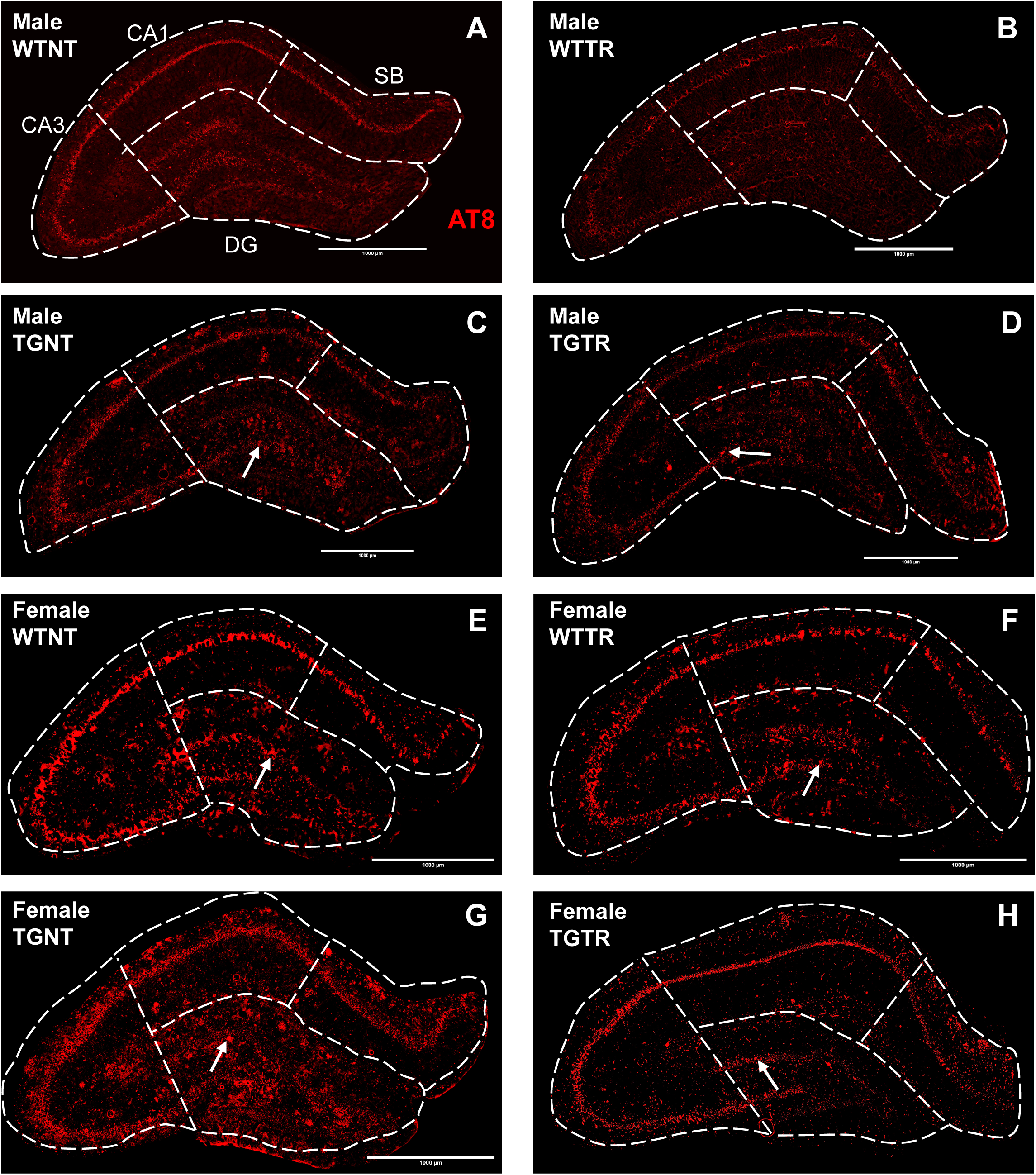
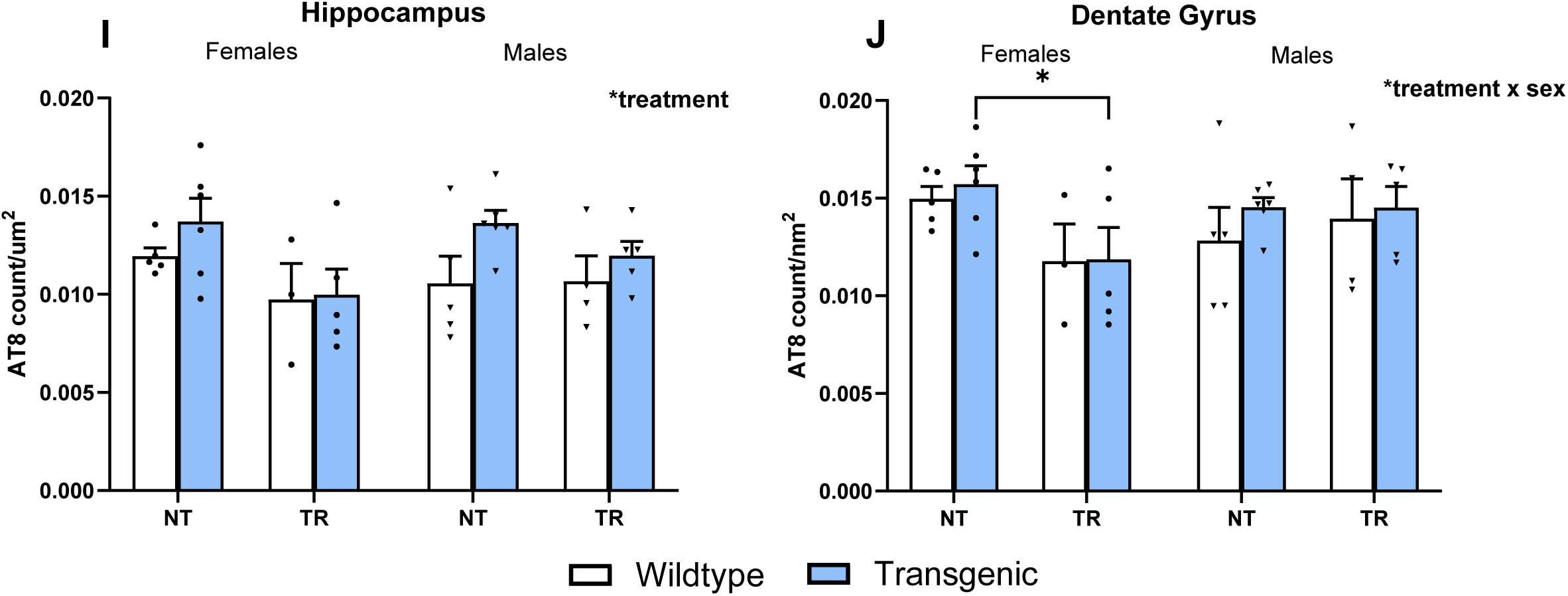
AGO decreases hyperphosphorylation of tau in a sex-dependant manner. (**A-H**) Representative images of dorsal hippocampi stained for AT8 (red). (**I-J**) Quantification of AT8 density shows no significant differences between WT and TgF344-AD rats in the hippocampus (genotype: *P* = 0.0538) or DG (genotype: *P* = 0.4061). Treatment with AGO decrease AT8 levels across the hippocampus (treatment: *P* = 0.0249) and induced a female specific decrease in AT8 levels in the DG (treatment*sex: *P* = 0.0357). AT8 was used to stain for phosphorylated tau (pS202/pT205/pS208) measured by count/µm2. Scale bar: 1000 µm. AGO: agomelatine; WTNT: wild-type not treated; WTTR: wild-type treated; TGNT: transgenic not treated; TGTR: transgenic treated, CA3: Cornu Ammonis 3, CA1: Cornu Ammonis 1, SB: subiculum, DG: dentate gyrus, F: female, M: male, G*: genotype effect, t*: treatment effect. Analysis was run using three-way ANOVA with Sidak’s post hoc tests; * p < 0.05, ** p < 0.01, *** p < 0.001, **** p < 0.0001.

### AGO elevated hippocampal neurogenesis (DCX^+^) without affecting mature (NeuN^+^) neurons in TgF344-AD rats

We evaluated hippocampal neurogenesis by quantifying doublecortin levels (DCX+) within the subgranular zone (SGZ), a neurogenic niche located along the inner border of the DG granule cell layer, noting that DCX labels immature neurons whereas NeuN marks mature, post-mitotic neurons. Representative SGZ labeling and the DCX expression mask are shown in Fig. 5A-H. DCX levels were significantly reduced in TgF344-AD rats compared to WT controls [genotype effect: *F*(1,31) = 4.977, *P* = 0.0331], indicating impaired neurogenic capacity in the transgenic rats (Fig. 5I). AGO treatment produced a significant overall increase in DCX expression across groups [treatment effect: *F*(1,31) = 4.461, *P* = 0.0428]. Full statistical analysis results are provided in Supplementary Table 8.

**Figure 5.**
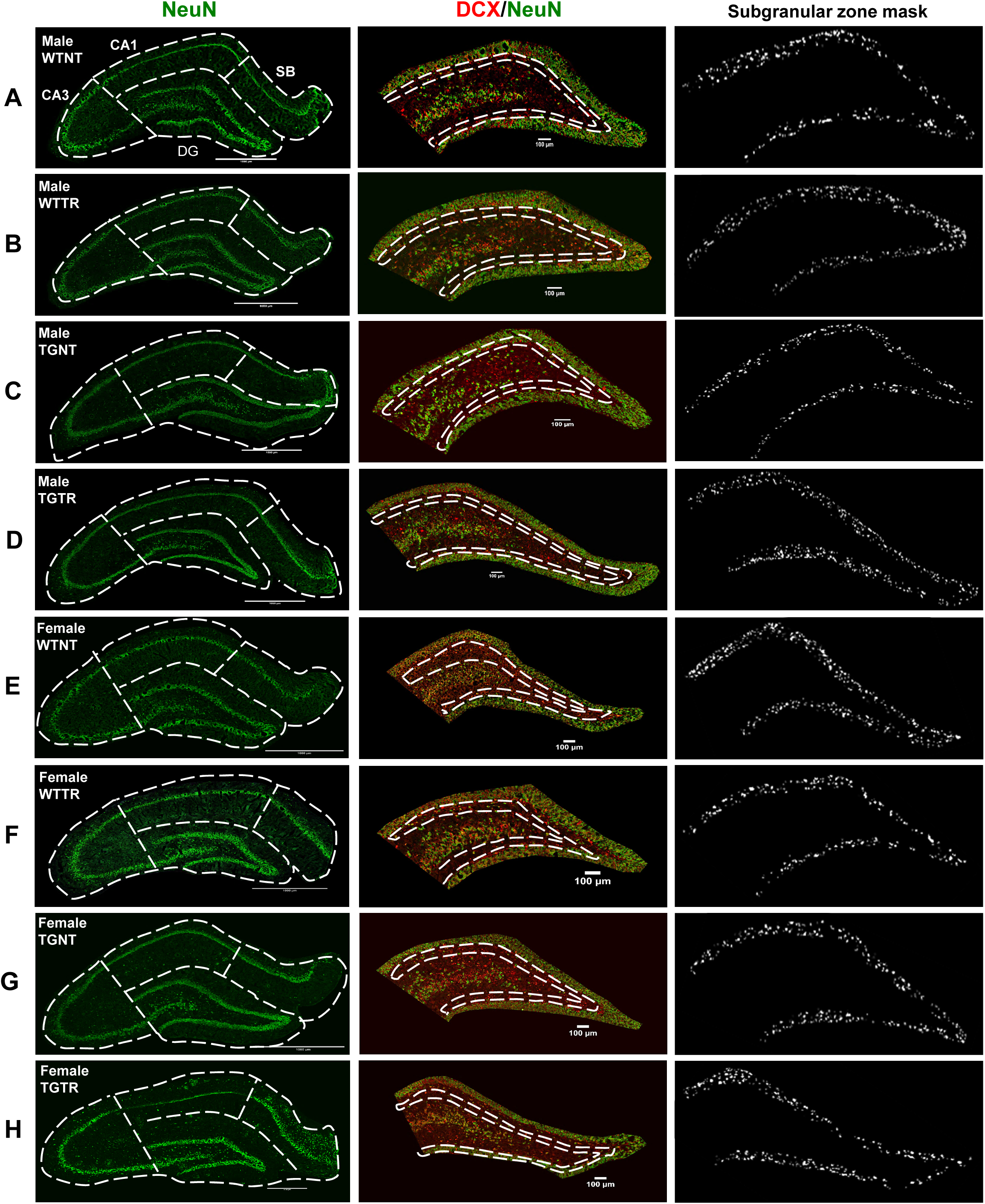
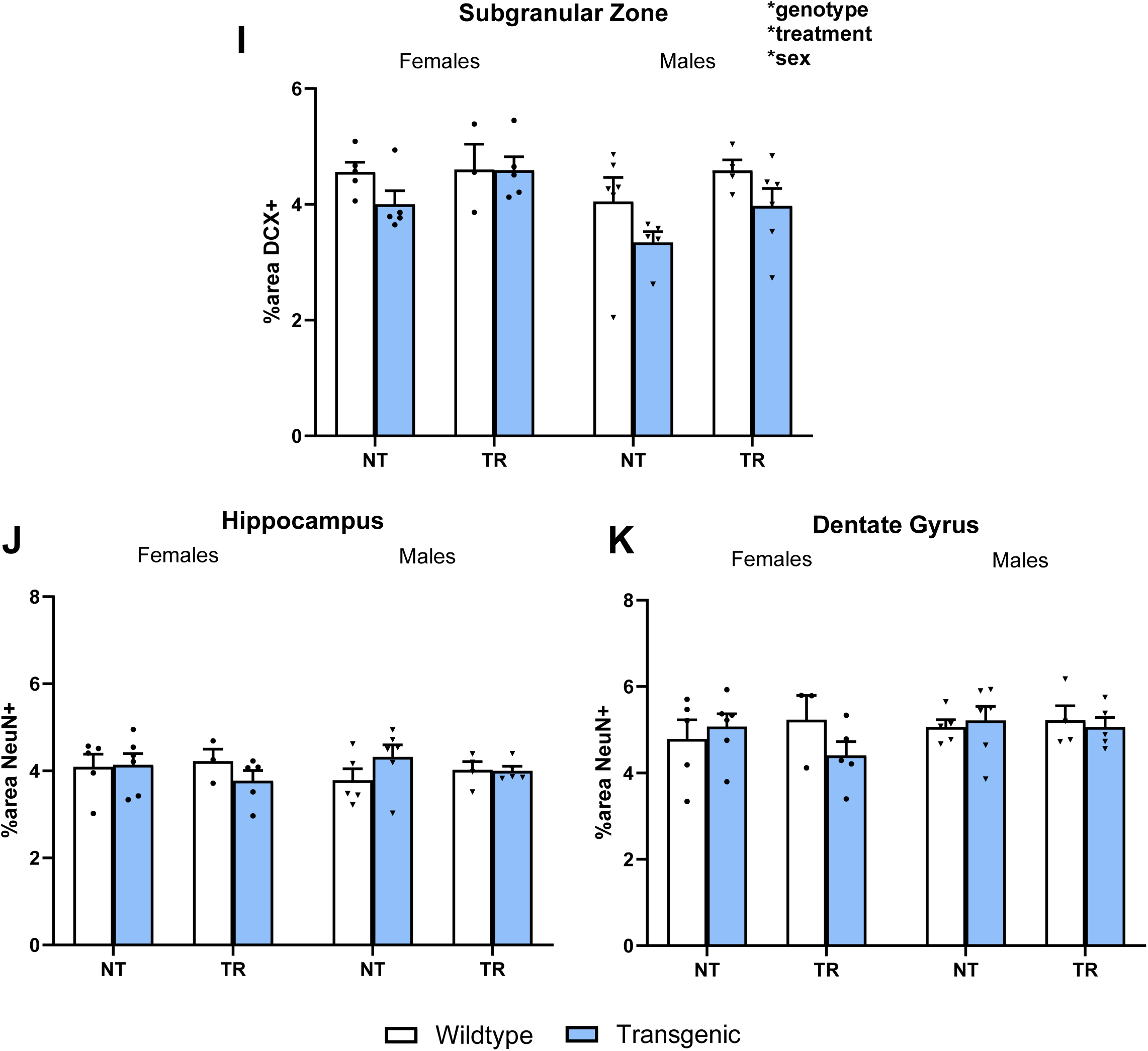
AGO selectively enhances subgranular zone neurogenesis without altering hippocampal neuronal density. (**A-H**) Representative images of dorsal hippocampi stained for NeuN (green) and subgranular zone stained for DCX (red) alongside mask (inverted) of positive DCX staining in the subgranular zone. (**I**) Quantification of DCX % area shows decreased DCX levels in TgF344-AD rats compared to WT (genotype: *P* = 0.0331). AGO increases DCX levels (treatment: *P* = 0.0428). Sex differences were also observed in DCX levels with females displaying higher levels of DCX staining (sex: *P* = 0.0423). (**J-K**) No differences among any groups were observed in NeuN levels across the hippocampus (genotype: *P* = 0.8803, treatment: *P* = 0.6636) or within the DG (genotype: *P* = 0.5727, treatment: *P* = 0.8314). Scale bar: 1000 µm. AGO: agomelatine; WTNT: wild-type not treated; WTTR: wild-type treated; TGNT: transgenic not treated; TGTR: transgenic treated, CA3: Cornu Ammonis 3, CA1: Cornu Ammonis 1, SB: subiculum, DG: dentate gyrus, F: female, M: male, G*: genotype effect, t*: treatment effect. Analysis was run using three-way ANOVA with Sidak’s post hoc tests; * p < 0.05, ** p < 0.01, *** p < 0.001, **** p < 0.0001.

To assess neuronal density, we quantified NeuN immunoreactivity (% area) across the hippocampus. NeuN levels did not differ by genotype or treatment [hippocampus genotype: *F*(1,31) = 0.02306, *P* = 0.8803; hippocampus treatment: *F*(1,31) = 0.1929, *P* = 0.6636; DG genotype: *F*(1,31) = 0.3250, *P*=0.5727; DG treatment: *F*(1,31) = 0.04609, *P* = 0.8314; Fig. 5J-K, Supplementary Table 9].

In sum, these findings indicate that AGO enhances DCX-defined neurogenesis despite reduced baseline DCX levels in TgF344-AD rats, while NeuN immunoreactivity, and thus overall neuronal density, remains unchanged across genotype and treatment groups. Full statistical results are provided in Supplementary Tables 8-9.

### AGO drives robust sex-dependent transcriptomic remodeling in the hippocampus of TgF344-AD rats

We evaluated transcriptional changes induced by AGO treatment in the dorsal hippocampus of male and female TgF344-AD rats. AGO treatment resulted in differential expression of 94 genes in females and 66 genes in males relative to untreated TgF344-AD controls (Fig. 6A-C).

**Figure 6.**
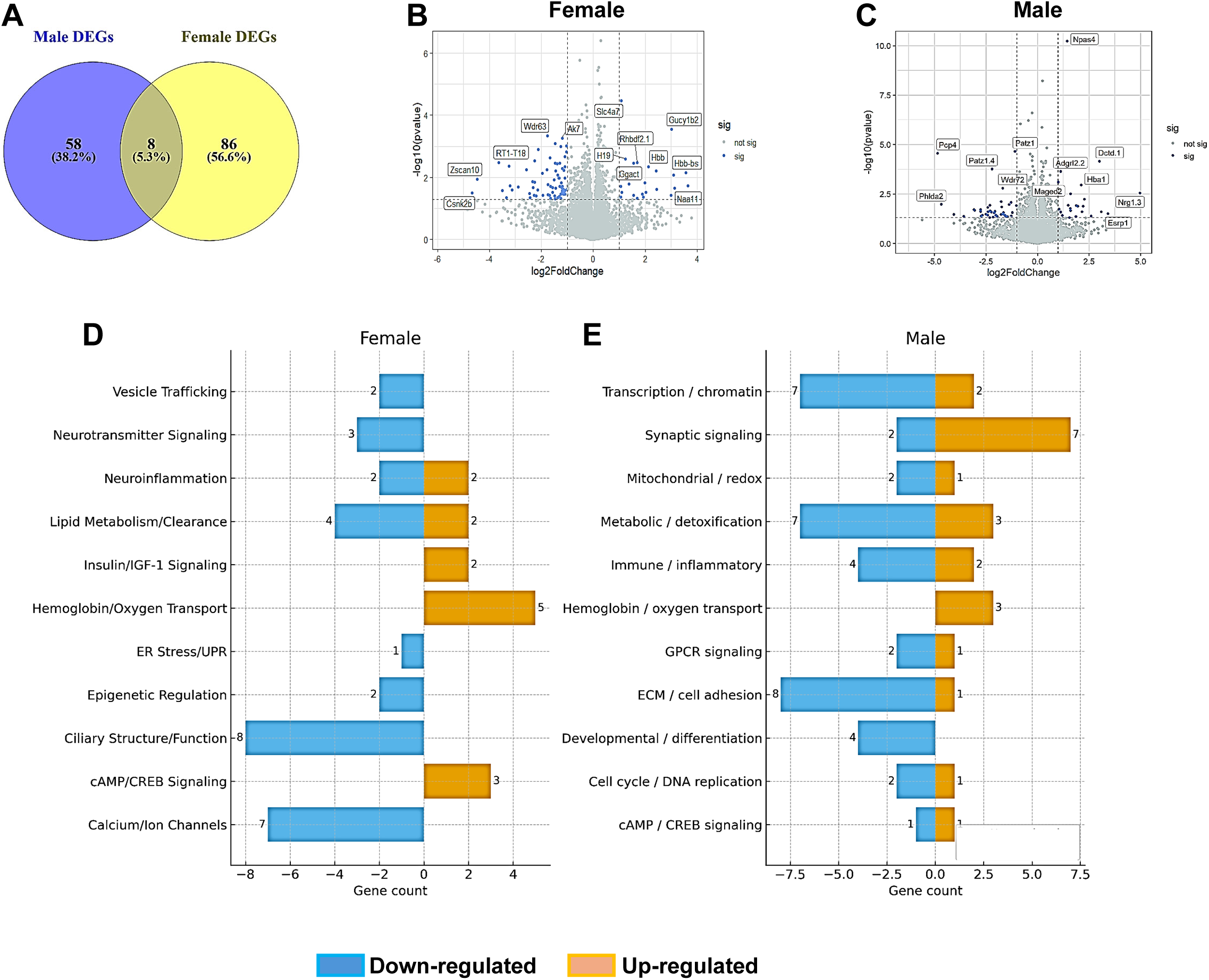
AGO elicits sex-dependent modulation of hippocampal gene expression in TgF344-AD rats. (**A**) AGO differentially regulated the expression of 94 genes in the hippocampus of Tg-AD female rats and 66 genes in the hippocampus of Tg-AD male rats. (**B-C**) Volcano plots show significant (blue) and non-significant genes (gray) in males (**B**) and females (**C**). Each point represents one gene. X-axis shows log2FoldChange and Y-axis shows –log10 (*P* value). Dashed lines signify the cutoffs for significance. Significance thresholds were set at *P* < 0.05 and |log2FoldChange| > 1. N = 5 per group. DESeq2 was used for DEG identification. (**D-E**) Modulated pathways between AGO treated and untreated TgF344-AD rats in females (**D**) and males (**E**). The gene count per pathway is shown on the X-axis. Gene sets are shown on the Y-axis. N = 5 per group. DEG: Differentially expressed gene.

Pathway-resolved analysis of the 94 female and 66 male DEGs revealed marked sex differences in the molecular programs engaged by AGO treatment (Fig. 6D-E, Supplementary Table 10A and B). DEGs were defined as those meeting *P* < 0.05 and an absolute log₂ fold-change ≥ 1.0, after filtering out low-count transcripts for reliable detection. Despite limited overlap at the individual gene level, both sexes exhibited convergence within several Alzheimer’s-relevant biological pathways.

In females, AGO up-regulated hemoglobin/oxygen-transport genes (*Hbb, Hba1, Hbb-b1, Hbb-bs*), a signature linked to altered oxygen trafficking and amyloid-β interactions in AD brains.^55^ Enhanced cAMP/CREB signaling (*Creb1*, *Pde4d*, *Pde4c*) accompanied increases in insulin/IGF-1 pathway components (*Igf1*, *H19*), reflecting AGO-driven activation of pro-synaptic and metabolic resilience programs.^56^ Up-regulation of *Cd36* and *Gpihbp1*, together with down-regulation of *Abcg5*, *Cav3*, *Crabp2*, and *Pon3*, indicated engagement of lipid-handling and clearance pathways implicated in Aβ processing and neuroinflammation. Neuroimmune modulation was also evident, with bidirectional changes in *Ggact*, *Ahr*, *Ptch2*, and *Adamtsl5*, consistent with glial activation states known to shift in AD.^57^ Females exhibited substantial down-regulation of ciliary-structure transcripts (e.g., *Ccdc40*, *Drc7*, *Cfap161*, *Dnah1*), aligning with reports of ciliary impairment in neurodegenerative contexts.^58^ Additional pathway-level effects included reductions in calcium/ion channel genes, epigenetic regulators (*Dnmt3a*, *Zmynd10*), ER-stress mediators (*Ddit3*), inhibitory neurotransmission, and vesicle-trafficking factors, indicating broad neuroprotective effects across multiple cellular compartments and pathways.

In males, AGO preferentially modulated pathways centered on neuronal excitability and transcriptional control. The synaptic-signaling module was strongly activated (*Nrg1*, *Chat*, *Synpo2l*, *Cartpt*, *P2rx3*, *Npas4*), consistent with restoration of plasticity-related networks vulnerable in AD.^59^ Hemoglobin/oxygen-transport transcripts were similarly increased, paralleling the female response.^55^ Modulation of immune/inflammatory genes (*Cd3e*, *Ceacam1*, *RT1-CE11*, *Phf11b*, *Lbp*, *Ceacam9*) reflected AGO-driven shifts in neuroimmune tone, a recognized driver of AD progression.^60^ AGO also altered cAMP/CREB signaling (*Pde4d* ↑; *Crem* ↓), extracellular-matrix and adhesion pathways (*Efemp2*, *Sgcg*, *Itgb6*), and metabolic/detoxification genes associated with lipid turnover and oxidative stress. Additional male-specific effects were observed in mitochondrial/redox pathways,^61^ transcription/chromatin regulators (*Esrp1*, *Dctd*, *Otx2*, *Patz1*), and cell-cycle/DNA-replication transcripts, consistent with dysregulated neuronal homeostasis in AD.^62^

Together, these findings indicate that AGO engages sex-specific yet AD-relevant biological pathways. Females displayed broader remodeling across metabolic, inflammatory, ciliary, and epigenetic domains, whereas males showed stronger modulation of synaptic, transcriptional, and mitochondrial programs. These differential pathway-level signatures provide a mechanistic basis for the sex-specific phenotypic outcomes observed following AGO treatment in TgF344-AD rats. Moreover, AGO drives a strong, sex-conserved induction of hemoglobin/oxygen-transport genes, with especially pronounced effects in females. This oxygen-handling signature linked to Aβ biology and metabolic stress in AD, suggests that AGO engages a shared protective pathway that may contribute to its therapeutic impact.

### AGO restores shared transcriptional deficits in PSEN2^N141I^ and APP^V717I^ iPSC-derived neurons

To validate AGO’s transcriptional effects in a human AD context, we used an isogenic iPSC-derived cortical neuron system carrying either the PSEN2N^141I^ or APP^V717I^ familial mutations. In untreated neurons, RNA-seq at day 35, when neurons are mature, revealed robust pathway-level alterations in both mutant lines relative to WT controls (Fig. 7A–D). Across all four groups (early and late DMSO conditions for each mutation), 102 genes were commonly up-regulated and 142 were commonly down-regulated, demonstrating a strong shared transcriptional signature (Fig.7 E-F). Full DEG lists are provided in Supplementary Table 11.

**Figure 7.**
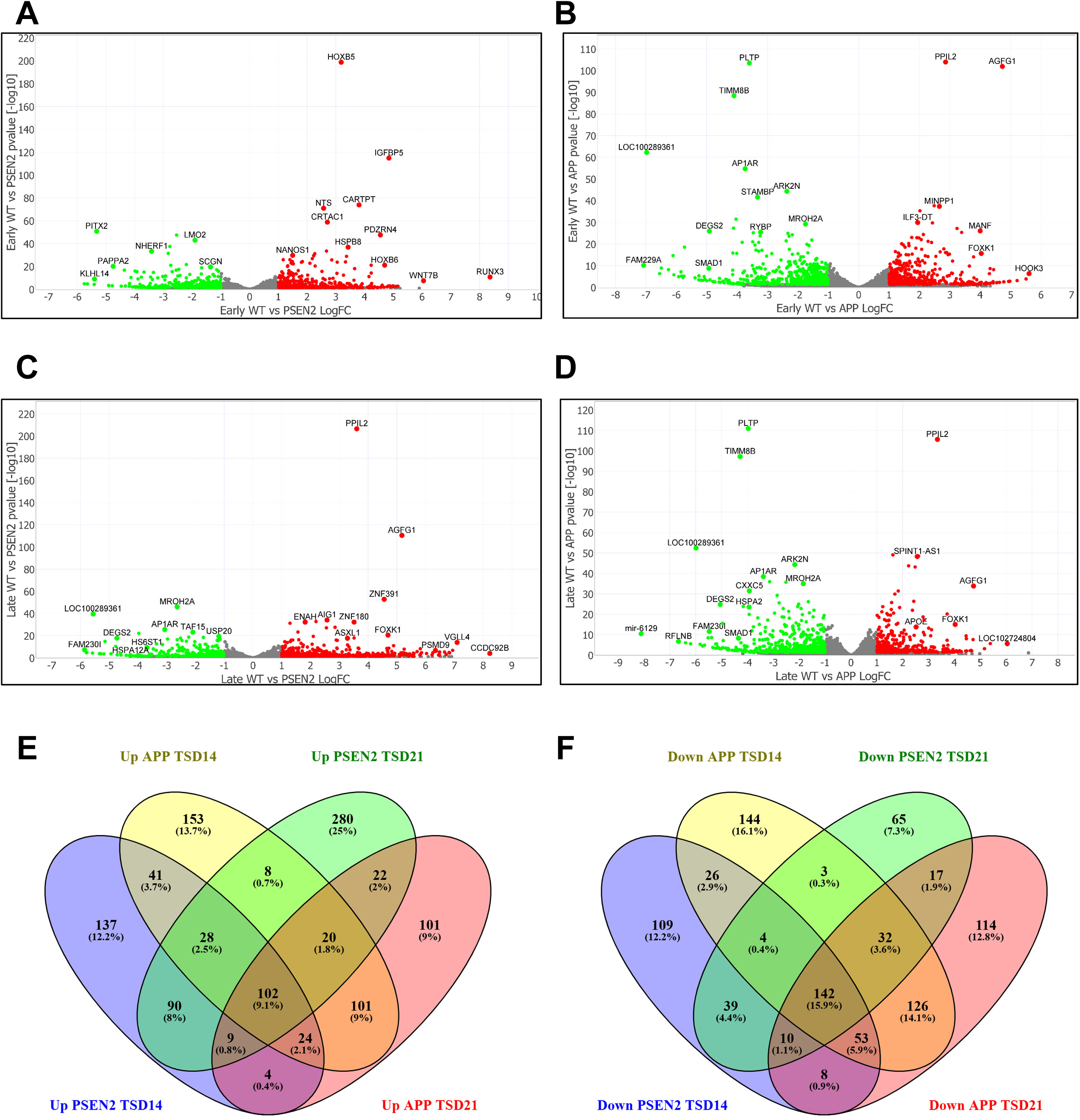
Distinct AD mutations converge on a shared transcriptomic signature of neuronal dysfunction. (**A-D**) Volcano plots show up-regulated (red), down-regulated (green), and non-significant (gray) genes across early DMSO treatment and late DMSO treatment for both PSEN2^N141I^ and APP^V717I^ mutant neurons. Each point represents one gene. X-axis shows log2FoldChange and Y-axis shows –log10 (*P* value). Dashed lines signify the cutoffs for significance. Significance thresholds were set at *P* < 0.05 and |log2FoldChange| > 1. (**E-F**) Among the DEGs identified, 102 are upregulated in all AD conditions (**E)** and 142 are down regulated in all AD conditions (**F)**. DEG: Differentially expressed gene.

Familial AD mutations in PSEN2^N141I^ and APP^V717I^ induced highly convergent transcriptional changes in iPSC-derived cortical neurons, with shared down- and up-regulated pathways across mutational drivers. Down-regulated DEGs enriched prominently for neurodevelopmental and axon-guidance regulators (*EPHA4, SLIT2/3, MET, ASTN2, SATB2*), synaptic-transmission genes (*ELFN1, NXPH1, GABRG3, CNTNAP2, SYNPO*), and ion-channel components governing neuronal excitability (*KCNJ3, KCNS1, NKAIN2/4*). Additional reductions in trophic-support pathways (*VEGFA, BMP2*), ECM and cytoskeletal regulators (*THBS2, SMOC1, ARHGAP24*), metabolic and redox-stress mediators (*PHGDH, SESN2, HMOX1*), ER-stress and proteostasis factors (*CHAC1, DDIT4*), and vesicle-trafficking genes (*TRIP10, RAB37*) indicated broad impairment of synaptic integrity, cellular metabolism, and protein-quality control.

In contrast, up-regulated DEGs reflected activation of aberrant developmental and patterning programs, including multiple HOX-family transcription factors (*HOXA1, HOXB2/3/5/6/7/8, HOXD-AS2*) and other neurodevelopmental regulators (*RUNX3, VAX2, TSHZ2, WNT7B*). Synaptic and neurotransmission-associated genes (*GABRA2/5, GRIK1, TRPC4, SLC17A6, RELN, CALB1, PVALB*) were also elevated, consistent with compensatory remodeling of excitatory–inhibitory signaling. Up-regulated ion-channel genes (*KCNJ2, KCNJ5, KCNH1, CACNG3*), trophic and chemokine mediators (*NTF3, CXCL12, VEGFC, TGFB2*), ECM and perineuronal-net components (*COL25A1, CRTAC1, CEMIP, TNR*), neuroimmune effectors (*IFITM2, HSPB8, WFDC2*), and metabolic/redox-associated transcripts (*ARRDC4, STEAP3, PLD6*) further pointed to widespread structural, inflammatory, and metabolic adaptation.

Together, these findings show that PSEN2^N141I^ and APP^V717I^ mutations drive a coordinated transcriptional state characterized by suppression of neuronal identity, synaptic function, excitability control, trophic signaling, and proteostasis, accompanied by aberrant activation of developmental, ECM-remodeling, neuroimmune, and stress-response programs in human cortical neurons.

AGO treatment produced broad transcriptomic effects in AD-mutant neurons (Fig. 8A–D). Notably, AGO up-regulated many genes that were suppressed in AD-mutant neurons and down-regulated genes that were elevated in AD-mutant neurons relative to WT. This bidirectional shift indicates a partial reversal of the AD-associated gene-expression profile (Fig. 8E–H), supporting AGO’s capacity to modulate pathological transcriptional signatures in human neurons. *PAPPA2*, a key gene studied in the context of stem cell research and a potential new target for AD,^63^ was found to be downregulated in both PSEN2^N141I^ and APP^V717I^ neurons. PAPPA2 (Pregnancy-associated plasma protein A2) is a protease involved in the growth hormone/IGF axis, crucial for development. Upon treatment, AGO restored *PAPPA2* expression confirming that AGO’s modulation of IGF signaling translates in human systems. Full DEG lists are provided in Supplementary Table 12-15.

**Figure 8.**
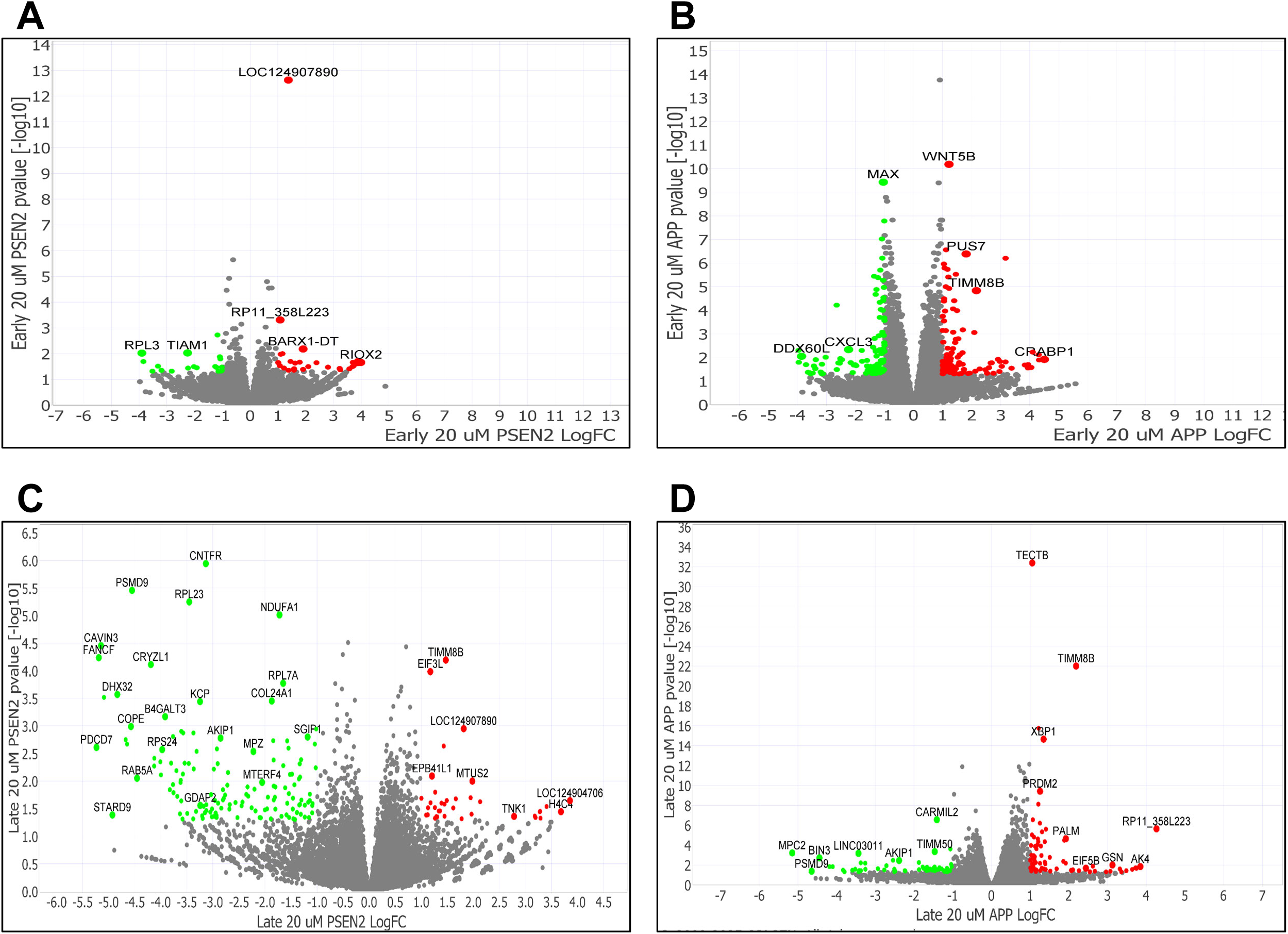
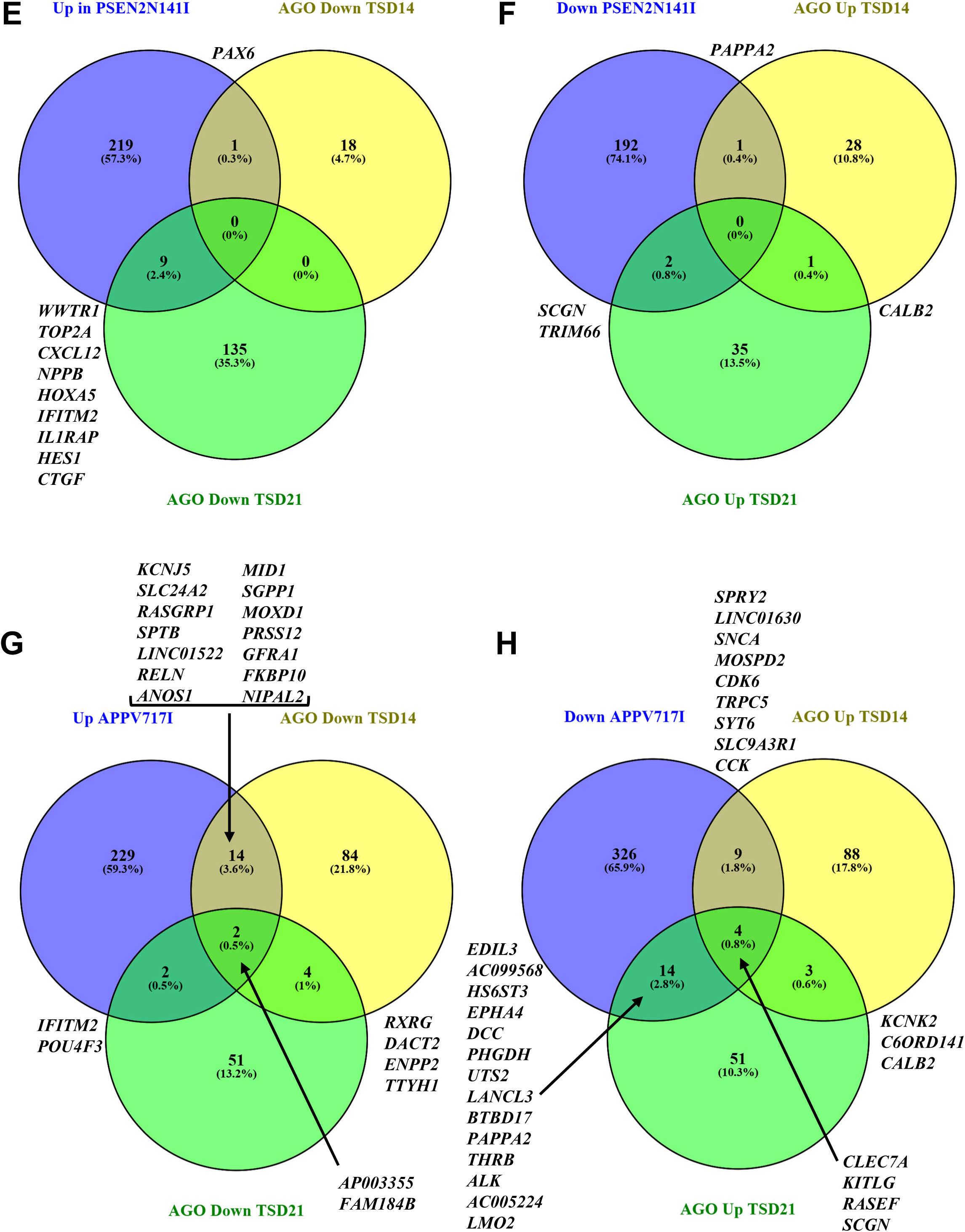
AGO reverses pathogenic transcriptomic signatures in human iPSC-derived AD-mutant cortical neurons. (**A-D**) Volcano plots show up-regulated (red), down-regulated (green), and non-significant genes (gray) genes across early AGO treatment and late AGO treatment for both PSEN2^N141I^ and APP^V717I^ mutant neurons. Each point represents one gene. X-axis shows log2FoldChange and Y-axis shows –log10 (*P* value). Dashed lines signify the cutoffs for significance. Significance thresholds were set at *P* < 0.05 and |log2FoldChange| > 1. (**E-H**) AGO reverses transcriptomic disruptions in PSEN2^N141I^ and APP^V717I^ mutant neurons. Gene names are provided. TSD: Treatment start date, DEG: Differentially expressed gene.

## Discussion

Our study identifies AGO as a sex-selective modulator of AD-related pathophysiology, restoring cognitive function, attenuating neuroinflammatory and tau-related pathology, and enhancing neurogenesis in female TgF344-AD rats, while engaging partially overlapping but distinct transcriptional programs in human iPSC-derived neurons carrying the PSEN2^N141I^ or APP^V717I^ mutations. Together, these findings uncover conserved and sex-divergent mechanisms by which AGO impacts AD-related molecular dysfunction and demonstrate the translational value of combining a rat model that recapitulates progressive, system-level pathology with human neurons that model early, cell specific effects of familial AD mutations.

### AGO improves cognition and neuronal plasticity through sex-specific mechanisms *in vivo*

AGO restored spatial learning selectively in female TgF344-AD rats at the age when cognitive decline first emerges, without altering performance in males. This behavioral rescue occurred despite no detectable effect on Aβ plaque burden, indicating that AGO acts downstream or independently of amyloid deposition. Instead, its beneficial effects aligned closely with reductions in microgliosis and tau hyperphosphorylation, two pathological processes increasingly recognized as critical drivers of cognitive impairment in AD.^64–72^ Importantly, microglial reactivity and tau pathology often exhibit sex-dependent trajectories in humans, consistent with the present findings. ^67–69,73^

AGO’s female-specific reduction of reactive and amoeboid microglia, together with restoration of DCX⁺ neurogenesis and DG-restricted reductions in AT8⁺ tau, suggests coordinated stabilization of hippocampal plasticity mechanisms.^57,74–76^ Neurogenesis deficits are a consistent early feature of AD,^77,78^ and microglia interactions within the DG strongly regulate progenitor survival and differentiation.^79^ These mechanistic links suggest that AGO’s enhancement of neurogenesis may arise secondary to normalization of neuroimmune tone and tau burden.^80^

### AGO responsiveness is driven by sex-specific transcriptomic remodeling

Bulk RNA-seq revealed that AGO engages widespread, sex-dependent transcriptional networks in the TgF344-AD hippocampus. Females showed remodeling across oxygen-handling, lipid metabolism, neuroimmune signalling, ciliary structure, calcium/ion-channel biology, and epigenetic pathways, which are disrupted early in AD and critical for synaptic plasticity, neurotrophic support, and metabolic resilience.^81^ Up-regulation of hemoglobin/oxygen-transport genes aligns with increasing recognition that neuronal hemoglobin participates in energy metabolism and amyloid–hemoglobin interactions relevant to AD pathology.^55,82^ These molecular changes may contribute to females’ enhanced therapeutic responsiveness.

In males, AGO preferentially modulated synaptic signaling, mitochondrial redox pathways, transcriptional control, and ion-channel biology. Although these neuronal pathways are also disrupted in AD,^83,84^ their restoration alone may be insufficient without the microglial and tau-modulatory effects seen in females. This asymmetry reinforces the view that sex-specific neuroimmune and metabolic effects shape treatment outcomes.^15,67–69,85^

### Shared AGO-responsive pathways in human iPSC-derived neurons and TgF344-AD rats

iPSC-derived cortical neurons harboring either the PSEN2^N141I^ or APP^V717I^ mutations exhibited a highly convergent set of disease-associated transcriptional changes, including down-regulation of synaptic genes, ion channels, trophic factors, proteostasis components, and metabolic regulators, which are hallmarks of early, cell-intrinsic AD vulnerability.^86–89^ Up-regulated pathways included HOX-driven developmental programs, ECM remodeling, neuroimmune mediators, and oxidative stress genes. This supports previous findings that identified common AD-endotypes in iPSC derived models of AD across mutational drivers.^33^

AGO partially reversed these transcriptional abnormalities, up-regulating genes suppressed in AD-mutant neurons and down-regulating those aberrantly elevated. Restoration of synaptic, mitochondrial, metabolic, and vesicle-trafficking pathways is consistent with AGO’s known neuromodulatory and antidepressant mechanisms,^90–92^ and aligns with the plasticity-based improvements and tau/microglial normalization observed *in vivo*.

Although this neuron-only system lacks microglia-neuron cross-talk, the overlap in AGO-responsive pathways suggests that AGO initiates conserved neuronal resilience programs that subsequently interact with sex-specific cellular contexts *in vivo*. This framework provides insight into AGO’s sex-specific behavioral and histopathological improvement in female rats but not males, despite shared upstream transcriptional targets.

The sex-dependent therapeutic profile of AGO likely reflects convergent effects of estrogen on CYP450-mediated drug metabolism and on melatonin receptor expression and downstream MT1/MT2 and ERK/CREB signaling, implying that AGO pharmacokinetics, receptor coupling, and neuroimmune interactions differ between males and females and amplify hippocampal responsiveness in females.^67,68^ Our results support microglial modulation as a key, sex-specific mechanism through which AGO confers neuroprotection, consistent with human studies demonstrating sex differences in neuroimmune activation, microglial aging, and inflammatory signaling.^93^ Further dose-response, pharmacokinetic, and hormone-depletion studies will be required to define the endocrine and molecular determinants of this sex bias.

### Integrating rat and human data

Across systems, AGO consistently modulated oxygen-handling, mitochondrial metabolism, synaptic signaling, neuroimmune-associated transcripts, and transcriptional regulators, which are fundamental pathways implicated in AD progression.^66,83,87^ Yet, some benefits observed *in vivo* (microglial normalization, DG-selective tau reduction, enhanced neurogenesis) require multicellular and systems-level interactions absent in culture. These results demonstrate the complementary value of combining a highly translational *in vivo* model with human neurons to dissect mechanisms governing therapeutic responsiveness.

## Conclusion

AGO exerts multifaceted, sex-dependent neuroprotective effects in TgF344-AD rats, improving cognition, reducing microgliosis and tau phosphorylation, and enhancing neurogenesis, supported by broad transcriptomic remodeling. Parallel studies in human AD-mutant cortical neurons revealed conserved AGO-mediated rebalancing of synaptic, metabolic, proteostatic, and developmental gene networks. These findings underscore AGO’s potential as a modulator of AD-relevant cellular programs and highlight the necessity of integrating human and animal models to reveal sex-specific therapeutic mechanisms in AD.

## Author contributions

G.T., P.S., P.R., and M.F.P., conceived the project and designed the experiments. L.X. performed the computational analysis supporting AGO treatment for AD. G.T. performed all experiments and analyzed the data. S.A. and N.R. provided essential technical assistance and support for the experimental work. G.T. and M.F.P wrote the manuscript. P.S. edited the manuscript. All authors approved the manuscript for submission.

## Supplementary material

Supplementary material is available.

