## Supplemental Tables and Figures for "Agomelatine drives sex-specific neuroprotection and reduced pathology in rat and human Alzheimer’s models"

**Supplemental Table 1 - Animals per group**

| Treatment | WT male | WT female | Tg-AD male | Tg-AD fema | Total |
|---|---|---|---|---|---|
| Agomelatine | 4 | 4 | 6 | 5 | 19 |
| Non-treated | 7 | 7 | 5 | 6 | 25 |
| Total | 11 | 11 | 11 | 11 | 44 |

**Supplemental Table 2A -Primary Antibodies**

| Antibody | Target | Dilution | Species Raised In | Manufacturer |
|---|---|---|---|---|
| Aß 4G8 | Amyloid beta plaques | 1:1000 | Mouse | Biolegend<br>Cat #800708 |
| Iba1 | Microglia | 1:500 | Rabbit | Wako<br>Cat #019-19742 |
| AT8 | P-tau Ser202, Thr205 | 1:250 | Mouse | ThermoFisher Cat #MN1020 |
| NeuN | Neurons | 1:200 | Chicken | Millipore<br>Cat #ABN91 |
| Doublecortin | Immature Neurons | 1:200 | Rabbit | Cell Signaling Cat#56130 |

**Supplemental Table 2B -Secondary Antibodies**

| Fluorophore | Reacts with | Raised In | Manufacture Info |
|---|---|---|---|
| Alexa Fluor 568 | Mouse | Goat | ThermoFisher<br>Cat #A-11031 |
| Alexa Fluor 488 | Rabbit | Goat | ThermoFisher<br>Cat #A-11008 |
| Alexa Fluor 488 | Chicken | Goat | ThermoFisher<br>Cat #11039 |
| Alexa Fluor 568 | Rabbit | Goat | ThermoFisher<br>Cat #A-11036 |

**Supplemental Table 3A -Latency to First Entrance Three way ANOVA Females 9 months**

| Source of Variation | % of total variation | P value | P value summary | Significant? | |
|---|---|---|---|---|---|
| Trial | 8.445 | 0.0117 | * | Yes | |
| Genotype | 3.122 | 0.0973 | ns | No | |
| Treatment | 0.02445 | 0.8792 | ns | No | |
| Trial x Genotype | 4.654 | 0.1373 | ns | No | |
| Trial x Treatment | 1.147 | 0.8314 | ns | No | |
| Genotype x Treatment | 0.005789 | 0.941 | ns | No | |
| Trial x Genotype x Treatment | 4.71 | 0.1326 | ns | No | |
| Subject | 20.64 | | | | |
| ANOVA table | SS | DF | MS | F (DFn, DFd) | P value |
| Trial | 318800 | 5 | 63760 | F (5, 100) = 3.118 | P=0.0117 |
| Genotype | 117853 | 1 | 117853 | F (1, 20) = 3.025 | P=0.0973 |
| Treatment | 922.8 | 1 | 922.8 | F (1, 20) = 0.02369 | P=0.8792 |
| Trial x Genotype | 175683 | 5 | 35137 | F (5, 100) = 1.718 | P=0.1373 |
| Trial x Treatment | 43296 | 5 | 8659 | F (5, 100) = 0.4235 | P=0.8314 |
| Genotype x Treatment | 218.5 | 1 | 218.5 | F (1, 20) = 0.005610 | P=0.9410 |
| Trial x Genotype x Treatment | 177817 | 5 | 35563 | F (5, 100) = 1.739 | P=0.1326 |
| Subject | 779081 | 20 | 38954 | | |
| Residual | 2044913 | 100 | 20449 | | |

**Supplemental Table 3B - Latency to First Entrance Three way ANOVA Males 9 months**

| Source of Variation | % of total variation | P value | P value summary | Significant? | |
|---|---|---|---|---|---|
| Trial | 16.09 | 0.0004 | *** | Yes | |
| Genotype | 1.416 | 0.2592 | ns | No | |
| Treatment | 0.7727 | 0.4001 | ns | No | |
| Trial x Genotype | 2.186 | 0.6194 | ns | No | |
| Trial x Treatment | 3.182 | 0.4063 | ns | No | |
| Genotype x Treatment | 2.212 | 0.1635 | ns | No | |
| Trial x Genotype x Treatment | 2.23 | 0.609 | ns | No | |
| Subject | 15.45 | | | | |
| ANOVA table | SS | DF | MS | F (DFn, DFd) | P value |
| Trial | 615090 | 5 | 123018 | F (5, 75) = 5.208 | P=0.0004 |
| Genotype | 54143 | 1 | 54143 | F (1, 15) = 1.375 | P=0.2592 |
| Treatment | 29538 | 1 | 29538 | F (1, 15) = 0.7502 | P=0.4001 |
| Trial x Genotype | 83583 | 5 | 16717 | F (5, 75) = 0.7078 | P=0.6194 |
| Trial x Treatment | 121625 | 5 | 24325 | F (5, 75) = 1.030 | P=0.4063 |
| Genotype x Treatment | 84554 | 1 | 84554 | F (1, 15) = 2.147 | P=0.1635 |
| Trial x Genotype x Treatment | 85254 | 5 | 17051 | F (5, 75) = 0.7219 | P=0.6090 |
| Subject | 590627 | 15 | 39375 | | |
| Residual | 1771404 | 75 | 23619 | | |

**Supplemental Table 4A - Latency to First Entrance Three way ANOVA Females 11 months**

| Source of Variation | % of total v | P value | P value summa | Significant? | |
|---|---|---|---|---|---|
| Trial | 14.37 | <0.0001 | **** | Yes | |
| Genotype | 4.901 | 0.0371 | * | Yes | |
| Treatment | 5.084 | 0.0341 | * | Yes | |
| Trial x Genotype | 1.58 | 0.6349 | ns | No | |
| Trial x Treatment | 7.43 | 0.01 | ** | Yes | |
| Genotype x Treatment | 2.94 | 0.0983 | ns | No | |
| Trial x Genotype x Treatment | 4.751 | 0.0772 | ns | No | |
| Subject | 17.41 | | | | |
| ANOVA table | SS | DF | MS | F (DFn, DFd) | P value |
| Trial | 1109876 | 5 | 221975 | F (5, 90) = 6.242 | P<0.0001 |
| Genotype | 378607 | 1 | 378607 | F (1, 18) = 5.068 | P=0.0371 |
| Treatment | 392753 | 1 | 392753 | F (1, 18) = 5.258 | P=0.0341 |
| Trial x Genotype | 122075 | 5 | 24415 | F (5, 90) = 0.6865 | P=0.6349 |
| Trial x Treatment | 574036 | 5 | 114807 | F (5, 90) = 3.228 | P=0.0100 |
| Genotype x Treatment | 227143 | 1 | 227143 | F (1, 18) = 3.041 | P=0.0983 |
| Trial x Genotype x Treatment | 367003 | 5 | 73401 | F (5, 90) = 2.064 | P=0.0772 |
| Subject | 1344644 | 18 | 74702 | | |
| Residual | 3200771 | 90 | 35564 | | |

**Supplemental Table 4B - Latency to First Entrance Three way ANOVA Males 11 months**

| Source of Variation | % of total v | P value | P value summ | Significant? | |
|---|---|---|---|---|---|
| Trial | 10.95 | 0.0043 | ** | Yes | |
| Genotype | 11.63 | 0.0009 | *** | Yes | |
| Treatment | 0.05817 | 0.7814 | ns | No | |
| Trial x Genotype | 3.578 | 0.311 | ns | No | |
| Trial x Treatment | 4.061 | 0.2419 | ns | No | |
| Genotype x Treatment | 0.7632 | 0.3211 | ns | No | |
| Trial x Genotype x Treatment | 2.573 | 0.5047 | ns | No | |
| Subject | 13.2 | | | | |
| ANOVA table | SS | DF | MS | F (DFn, DFd) | P value |
| Trial | 800602 | 5 | 160120 | F (5, 90) = 3.703 | P=0.0043 |
| Genotype | 849818 | 1 | 849818 | F (1, 18) = 15.85 | P=0.0009 |
| Treatment | 4252 | 1 | 4252 | F (1, 18) = 0.07933 | P=0.7814 |
| Trial x Genotype | 261558 | 5 | 52312 | F (5, 90) = 1.210 | P=0.3110 |
| Trial x Treatment | 296886 | 5 | 59377 | F (5, 90) = 1.373 | P=0.2419 |
| Genotype x Treatment | 55791 | 1 | 55791 | F (1, 18) = 1.041 | P=0.3211 |
| Trial x Genotype x Treatment | 188050 | 5 | 37610 | F (5, 90) = 0.8698 | P=0.5047 |
| Subject | 964795 | 18 | 53600 | | |
| Residual | 3891526 | 90 | 43239 | | |

**Supplemental Table 4C - Latency to First Entrance Three way ANOVA Males vs Females Transgenic**

| Source of Variation | % of total v | P value | P value summ | Significant? | |
|---|---|---|---|---|---|
| Trial | 14.1 | 0.0002 | *** | Yes | |
| Treatment | 3.374 | 0.0494 | * | Yes | |
| Sex | 0.8658 | 0.3 | ns | No | |
| Trial x Treatment | 3.559 | 0.2285 | ns | No | |
| Trial x Sex | 1.001 | 0.8502 | ns | No | |
| Treatment x Sex | 10.35 | 0.0017 | ** | Yes | |
| Trial x Treatment x Sex | 9.98 | 0.0028 | ** | Yes | |
| Subject | 13.68 | | | | |
| ANOVA table | SS | DF | MS | F (DFn, DFd) | P value |
| Trial | 696081 | 5 | 139216 | F (5, 90) = 5.586 | P=0.0002 |
| Treatment | 166497 | 1 | 166497 | F (1, 18) = 4.438 | P=0.0494 |
| Sex | 42730 | 1 | 42730 | F (1, 18) = 1.139 | P=0.3000 |
| Trial x Treatment | 175628 | 5 | 35126 | F (5, 90) = 1.409 | P=0.2285 |
| Trial x Sex | 49390 | 5 | 9878 | F (5, 90) = 0.3963 | P=0.8502 |
| Treatment x Sex | 510915 | 1 | 510915 | F (1, 18) = 13.62 | P=0.0017 |
| Trial x Treatment x Sex | 492534 | 5 | 98507 | F (5, 90) = 3.952 | P=0.0028 |
| Subject | 675323 | 18 | 37518 | | |
| Residual | 2243046 | 90 | 24923 | | |

**Supplemental Table 4D - Latency to First Entrance Three way ANOVA Males vs Females Wildtype**

| Source of Variation | % of total v | P value | P value summ | Significant? | |
|---|---|---|---|---|---|
| Trial | 15.91 | 0.0003 | *** | Yes | |
| treatment | 0.282 | 0.6067 | ns | No | |
| Sex | 0.117 | 0.7396 | ns | No | |
| Trial x treatment | 4.983 | 0.159 | ns | No | |
| Trial x Sex | 1.317 | 0.8252 | ns | No | |
| treatment x Sex | 0.001193 | 0.9732 | ns | No | |
| Trial x treatment x Sex | 3.559 | 0.3315 | ns | No | |
| Subject | 18.49 | | | | |
| ANOVA table | SS | DF | MS | F (DFn, DFd) | P value |
| Trial | 1405479 | 5 | 281096 | F (5, 90) = 5.217 | P=0.0003 |
| treatment | 24917 | 1 | 24917 | F (1, 18) = 0.2745 | P=0.6067 |
| Sex | 10341 | 1 | 10341 | F (1, 18) = 0.1139 | P=0.7396 |
| Trial x treatment | 440329 | 5 | 88066 | F (5, 90) = 1.634 | P=0.1590 |
| Trial x Sex | 116388 | 5 | 23278 | F (5, 90) = 0.4320 | P=0.8252 |
| treatment x Sex | 105.5 | 1 | 105.5 | F (1, 18) = 0.001162 | P=0.9732 |
| Trial x treatment x Sex | 314495 | 5 | 62899 | F (5, 90) = 1.167 | P=0.3315 |
| Subject | 1634117 | 18 | 90784 | | |
| Residual | 4849252 | 90 | 53881 | | |

#### Supplemental Table 5A - Whole Hippocampus %area Abeta+

| 2way ANOVA | | | | | |
|---|---|---|---|---|---|
| Source of Variation | % of total variation | P value | P value summary | Significant? | |
| Interaction | 0.729 | 0.7112 | ns | No | |
| Sex | 10.41 | 0.1729 | ns | No | |
| Treatment | 0.823 | 0.6941 | ns | No | |
| ANOVA table | SS (Type III) | DF | MS | F (DFn, DFd) | P value |
| Interaction | 0.009018 | 1 | 0.009018 | F (1, 17) = 0.1418 | P=0.7112 |
| Sex | 0.1287 | 1 | 0.1287 | F (1, 17) = 2.024 | P=0.1729 |
| Treatment | 0.01018 | 1 | 0.01018 | F (1, 17) = 0.1601 | P=0.6941 |
| Residual | 1.081 | 17 | 0.0636 | | |
| Šídák's multiple comparisons te | Predicted (LS) Mean diff. | 95.00% CI o | Adjusted P Value | | |
| Male | | | | | |
| TGNT vs. TGTR | 0.08575 | -0.2887 to 0. | 0.8251 | | |
| Female | | | | | |
| TGNT vs. TGTR | 0.0026 | -0.3885 to 0. | 0.9998 | | |
| TGNT | | | | | |
| Male vs. Female | -0.1155 | -0.5066 to 0. | 0.7284 | | |
| TGTR | | | | | |
| Male vs. Female | -0.1987 | -0.5731 to 0. | 0.377 | | |

#### Supplemental Table 5B - Dentate Gyrus %area Abeta+

| 2way ANOVA | | | | | |
|---|---|---|---|---|---|
| Source of Variation | % of total variation | P value | P value summary | Significant? | |
| Interaction | 7.616 | 0.2158 | ns | No | |
| Sex | 7.775 | 0.2113 | ns | No | |
| Treatment | 4.436 | 0.3402 | ns | No | |
| ANOVA table | SS (Type III) | DF | MS | F (DFn, DFd) | P value |
| Interaction | 0.1095 | 1 | 0.1095 | F (1, 17) = 1.653 | P=0.2158 |
| Sex | 0.1118 | 1 | 0.1118 | F (1, 17) = 1.687 | P=0.2113 |
| Treatment | 0.06377 | 1 | 0.06377 | F (1, 17) = 0.9629 | P=0.3402 |
| Residual | 1.126 | 17 | 0.06623 | | |
| Šídák's multiple comparisons te | Predicted (LS) Mean diff. | 95.00% CI o | Adjusted P Value | | |
| Male | | | | | |
| TGNT vs. TGTR | 0.2554 | -0.1267 to 0. | 0.2249 | | |
| Female | | | | | |
| TGNT vs. TGTR | -0.0343 | -0.4334 to 0. | 0.973 | | |
| TGNT | | | | | |
| Male vs. Female | -0.0015 | -0.4006 to 0 | >0.9999 | | |
| TGTR | | | | | |
| Male vs. Female | -0.2912 | -0.6733 to 0. | 0.1517 | | |

#### Supplemental Table 5C- CA1 %area Abeta+

| 2way ANOVA | | | | | |
|---|---|---|---|---|---|
| Source of Variation | % of total variation | P value | P value summary | Significant? | |
| Interaction | 0.02243 | 0.9509 | ns | No | |
| Sex | 0.06112 | 0.919 | ns | No | |
| Treatment | 2.369 | 0.5292 | ns | No | |
| ANOVA table | SS (Type III) | DF | MS | F (DFn, DFd) | P value |
| Interaction | 0.0003725 | 1 | 0.0003725 | F (1, 17) = 0.00390 | P=0.9509 |
| Sex | 0.001015 | 1 | 0.001015 | F (1, 17) = 0.01065 | P=0.9190 |
| Treatment | 0.03935 | 1 | 0.03935 | F (1, 17) = 0.4127 | P=0.5292 |
| Residual | 1.621 | 17 | 0.09536 | | |
| Šídák's multiple comparisons te | Predicted (LS) Mean diff. | 95.00% CI o | Adjusted P Value | | |
| Male | | | | | |
| TGNT vs. TGTR | -0.0784 | -0.5369 to 0. | 0.8978 | | |
| Female | | | | | |
| TGNT vs. TGTR | -0.0953 | -0.5741 to 0. | 0.8644 | | |

| | | | |
|---|---|---|---|
| TGNT | | | |
| Male vs. Female | -0.0055 | -0.4843 to 0. | 0.9995 |
| TGTR | | | |
| Male vs. Female | -0.0224 | -0.4809 to 0. | 0.9912 |

#### Supplemental Table 5D - CA3 %area Abeta+

| | | | | | |
|---|---|---|---|---|---|
| 2way ANOVA | | | | | |
| Source of Variation | % of total variation | P value | P value summary | Significant? | |
| Interaction | 8.778 | 0.2136 | ns | No | |
| Sex | 6.419 | 0.2844 | ns | No | |
| Treatment | 1.087 | 0.6547 | ns | No | |
| ANOVA table | SS | DF | MS | F (DFn, DFd) | P value |
| Interaction | 0.155 | 1 | 0.155 | F (1, 16) = 1.678 | P=0.2136 |
| Sex | 0.1133 | 1 | 0.1133 | F (1, 16) = 1.227 | P=0.2844 |
| Treatment | 0.01919 | 1 | 0.01919 | F (1, 16) = 0.2077 | P=0.6547 |
| Residual | 1.478 | 16 | 0.09237 | | |
| Šídák's multiple comparisons te: | Mean diff. | 95.00% CI o | Adjusted P Value | | |
| Male | | | | | |
| TGNT vs. TGTR | 0.238 | -0.2361 to 0. | 0.4125 | | |
| Female | | | | | |
| TGNT vs. TGTR | -0.1141 | -0.5882 to 0. | 0.8074 | | |
| TGNT | | | | | |
| Male vs. Female | 0.0255 | -0.4486 to 0. | 0.9892 | | |
| TGTR | | | | | |
| Male vs. Female | -0.3266 | -0.8007 to 0. | 0.2055 | | |

#### Supplemental Table 5E - SB %area Abeta+

| | | | | | |
|---|---|---|---|---|---|
| 2way ANOVA | | | | | |
| Source of Variation | % of total variation | P value | P value summary | Significant? | |
| Interaction | 3.283 | 0.4114 | ns | No | |
| Sex | 16.39 | 0.0771 | ns | No | |
| Treatment | 1.921 | 0.5281 | ns | No | |
| ANOVA table | SS (Type III) | DF | MS | F (DFn, DFd) | P value |
| Interaction | 0.08678 | 1 | 0.08678 | F (1, 17) = 0.7092 | P=0.4114 |
| Sex | 0.4332 | 1 | 0.4332 | F (1, 17) = 3.540 | P=0.0771 |
| Treatment | 0.05076 | 1 | 0.05076 | F (1, 17) = 0.4148 | P=0.5281 |
| Residual | 2.08 | 17 | 0.1224 | | |
| Šídák's multiple comparisons te: | Predicted (LS) Mean diff. | 95.00% CI o | Adjusted P Value | | |
| Male | | | | | |
| TGNT vs. TGTR | -0.03033 | -0.5497 to 0. | 0.9874 | | |
| Female | | | | | |
| TGNT vs. TGTR | 0.2276 | -0.3148 to 0. | 0.5349 | | |
| TGNT | | | | | |
| Male vs. Female | -0.4171 | -0.9595 to 0. | 0.1473 | | |
| TGTR | | | | | |
| Male vs. Female | -0.1592 | -0.6785 to 0. | 0.7113 | | |

#### Supplemental Table 6A - Whole Hippocampus %area ramified

| 3way ANOVA | | | | | |
|---|---|---|---|---|---|
| Source of Variation | % of total variation | P value | P value summa | Significant? | |
| Treatment | 2.7 | 0.3351 | ns | No | |
| Sex | 0.1592 | 0.8136 | ns | No | |
| Genotype | 2.569 | 0.3469 | ns | No | |
| Treatment x Sex | 0.1067 | 0.8469 | ns | No | |
| Treatment x Genotype | 0.6482 | 0.6348 | ns | No | |
| Sex x Genotype | 1.281 | 0.5051 | ns | No | |
| Treatment x Sex x Genotype | 4.342 | 0.2237 | ns | No | |
| ANOVA table | SS (Type III) | DF | MS | F (DFn, DFd) | P value |
| Treatment | 0.4787 | 1 | 0.4787 | F (1, 31) = 0.9586 | P=0.3351 |
| Sex | 0.02823 | 1 | 0.02823 | F (1, 31) = 0.05653 | P=0.8136 |
| Genotype | 0.4555 | 1 | 0.4555 | F (1, 31) = 0.9123 | P=0.3469 |
| Treatment x Sex | 0.01892 | 1 | 0.01892 | F (1, 31) = 0.03789 | P=0.8469 |
| Treatment x Genotype | 0.1149 | 1 | 0.1149 | F (1, 31) = 0.2302 | P=0.6348 |
| Sex x Genotype | 0.227 | 1 | 0.227 | F (1, 31) = 0.4547 | P=0.5051 |
| Treatment x Sex x Genotype | 0.7698 | 1 | 0.7698 | F (1, 31) = 1.542 | P=0.2237 |
| Residual | 15.48 | 31 | 0.4993 | | |
| Šídák's multiple comparisons test | Predicted (LS) Mean diff. | 95.00% CI of diff. | Below threshol | Summary | Adjusted P |
| NT:Female WT vs. TR:Female WT | -0.3631 | -1.904 to 1.178 | No | ns | 0.9995 |
| NT:Female Tg-AD vs. TR:Female Tg-AD | -0.0053 | -1.384 to 1.373 | No | ns | >0.9999 |
| NT:Male WT vs. TR:Male WT | 0.1286 | -1.413 to 1.670 | No | ns | >0.9999 |
| NT:Male Tg-AD vs. TR:Male Tg-AD | -0.6798 | -2.000 to 0.6400 | No | ns | 0.7909 |
| NT:Female WT vs. NT:Female Tg-AD | -0.113 | -1.433 to 1.207 | No | ns | >0.9999 |
| NT:Male WT vs. NT:Male Tg-AD | 0.7868 | -0.5330 to 2.107 | No | ns | 0.6104 |
| TR:Female WT vs. TR:Female Tg-AD | 0.2448 | -1.347 to 1.837 | No | ns | >0.9999 |
| TR:Male WT vs. TR:Male Tg-AD | -0.02158 | -1.563 to 1.520 | No | ns | >0.9999 |
| NT:Female WT vs. NT:Male WT | -0.3483 | -1.607 to 0.9100 | No | ns | 0.9978 |
| NT:Female Tg-AD vs. NT:Male Tg-AD | 0.5514 | -0.8270 to 1.930 | No | ns | 0.9542 |
| TR:Female WT vs. TR:Male WT | 0.1433 | -1.636 to 1.923 | No | ns | >0.9999 |
| TR:Female Tg-AD vs. TR:Male Tg-AD | -0.1231 | -1.443 to 1.197 | No | ns | >0.9999 |

#### Supplemental Table 6B - Whole Hippocampus %area reactive

| 3way ANOVA | | | | | |
|---|---|---|---|---|---|
| Source of Variation | % of total variation | P value | P value summa | Significant? | |
| Treatment | 0.009767 | 0.9337 | ns | No | |
| Sex | 9.935 | 0.0119 | * | Yes | |
| Genotype | 20.97 | 0.0005 | *** | Yes | |
| Treatment x Sex | 2.431 | 0.1956 | ns | No | |
| Treatment x Genotype | 7.251 | 0.0294 | * | Yes | |
| Sex x Genotype | 1.736 | 0.2723 | ns | No | |
| Treatment x Sex x Genotype | 7.914 | 0.0233 | * | Yes | |
| ANOVA table | SS (Type III) | DF | MS | F (DFn, DFd) | P value |
| Treatment | 0.0001223 | 1 | 0.0001223 | F (1, 31) = 0.00702 | P=0.9337 |
| Sex | 0.1244 | 1 | 0.1244 | F (1, 31) = 7.149 | P=0.0119 |
| Genotype | 0.2626 | 1 | 0.2626 | F (1, 31) = 15.09 | P=0.0005 |
| Treatment x Sex | 0.03045 | 1 | 0.03045 | F (1, 31) = 1.749 | P=0.1956 |
| Treatment x Genotype | 0.09081 | 1 | 0.09081 | F (1, 31) = 5.217 | P=0.0294 |
| Sex x Genotype | 0.02174 | 1 | 0.02174 | F (1, 31) = 1.249 | P=0.2723 |
| Treatment x Sex x Genotype | 0.09911 | 1 | 0.09911 | F (1, 31) = 5.694 | P=0.0233 |
| Residual | 0.5395 | 31 | 0.0174 | | |
| Šídák's multiple comparisons test | Predicted (LS) Mean diff. | 95.00% CI of diff. | Below threshol | Summary | Adjusted P |
| NT:Female WT vs. TR:Female WT | -0.1431 | -0.4308 to 0.1446 | No | ns | 0.8251 |
| NT:Female Tg-AD vs. TR:Female Tg-AD | 0.2664 | 0.009046 to 0.5238 | Yes | * | 0.038 |
| NT:Male WT vs. TR:Male WT | -0.04983 | -0.3376 to 0.2379 | No | ns | >0.9999 |
| NT:Male Tg-AD vs. TR:Male Tg-AD | -0.05878 | -0.3052 to 0.1876 | No | ns | 0.9995 |
| NT:Female WT vs. NT:Female Tg-AD | -0.424 | -0.6704 to -0.1776 | Yes | *** | 0.0001 |
| NT:Male WT vs. NT:Male Tg-AD | -0.1168 | -0.3632 to 0.1296 | No | ns | 0.8652 |

| | | | | | |
|---|---|---|---|---|---|
| TR:Female WT vs. TR:Female Tg-AD | -0.01453 | -0.3117 to 0.2826 | No | ns | >0.9999 |
| TR:Male WT vs. TR:Male Tg-AD | -0.1258 | -0.4135 to 0.1620 | No | ns | 0.9171 |
| NT:Female WT vs. NT:Male WT | 0.02158 | -0.2133 to 0.2565 | No | ns | >0.9999 |
| NT:Female Tg-AD vs. NT:Male Tg-AD | 0.3288 | 0.07145 to 0.5862 | Yes | ** | 0.0052 |
| TR:Female WT vs. TR:Male WT | 0.1148 | -0.2174 to 0.4471 | No | ns | 0.9848 |
| TR:Female Tg-AD vs. TR:Male Tg-AD | 0.003617 | -0.2428 to 0.2500 | No | ns | >0.9999 |

#### Supplemental Table 6C - Whole Hippocampus %area ameoboid

| 3way ANOVA | | | | | |
|---|---|---|---|---|---|
| Source of Variation | % of total variation | P value | P value summa | Significant? | |
| Treatment | 0.1962 | 0.7091 | ns | No | |
| Sex | 12.77 | 0.0048 | ** | Yes | |
| Genotype | 16.35 | 0.0017 | ** | Yes | |
| Treatment x Sex | 1.884 | 0.2523 | ns | No | |
| Treatment x Genotype | 9.034 | 0.0158 | * | Yes | |
| Sex x Genotype | 1.548 | 0.2985 | ns | No | |
| Treatment x Sex x Genotype | 9.618 | 0.013 | * | Yes | |
| ANOVA table | SS (Type III) | DF | MS | F (DFn, DFd) | P value |
| Treatment | 0.001208 | 1 | 0.001208 | F (1, 31) = 0.1417 | P=0.7091 |
| Sex | 0.07865 | 1 | 0.07865 | F (1, 31) = 9.226 | P=0.0048 |
| Genotype | 0.1007 | 1 | 0.1007 | F (1, 31) = 11.81 | P=0.0017 |
| Treatment x Sex | 0.0116 | 1 | 0.0116 | F (1, 31) = 1.361 | P=0.2523 |
| Treatment x Genotype | 0.05564 | 1 | 0.05564 | F (1, 31) = 6.527 | P=0.0158 |
| Sex x Genotype | 0.009532 | 1 | 0.009532 | F (1, 31) = 1.118 | P=0.2985 |
| Treatment x Sex x Genotype | 0.05924 | 1 | 0.05924 | F (1, 31) = 6.948 | P=0.0130 |
| Residual | 0.2643 | 31 | 0.008525 | | |
| Šídák's multiple comparisons test | Predicted (LS) Mean diff. | 95.00% CI of diff. | Below threshol | Summary | Adjusted P |
| NT:Female WT vs. TR:Female WT | -0.1119 | -0.3133 to 0.08946 | No | ns | 0.704 |
| NT:Female Tg-AD vs. TR:Female Tg-AD | 0.2066 | 0.02648 to 0.3867 | Yes | * | 0.0154 |
| NT:Male WT vs. TR:Male WT | -0.02175 | -0.2231 to 0.1796 | No | ns | >0.9999 |
| NT:Male Tg-AD vs. TR:Male Tg-AD | -0.02673 | -0.1992 to 0.1457 | No | ns | >0.9999 |
| NT:Female WT vs. NT:Female Tg-AD | -0.2972 | -0.4696 to -0.1247 | Yes | *** | 0.0001 |
| NT:Male WT vs. NT:Male Tg-AD | -0.07052 | -0.2430 to 0.1019 | No | ns | 0.9466 |
| TR:Female WT vs. TR:Female Tg-AD | 0.02137 | -0.1866 to 0.2293 | No | ns | >0.9999 |
| TR:Male WT vs. TR:Male Tg-AD | -0.0755 | -0.2769 to 0.1259 | No | ns | 0.9714 |
| NT:Female WT vs. NT:Male WT | 0.01567 | -0.1488 to 0.1801 | No | ns | >0.9999 |
| NT:Female Tg-AD vs. NT:Male Tg-AD | 0.2423 | 0.06218 to 0.4224 | Yes | ** | 0.0029 |
| TR:Female WT vs. TR:Male WT | 0.1058 | -0.1267 to 0.3384 | No | ns | 0.8936 |
| TR:Female Tg-AD vs. TR:Male Tg-AD | 0.008967 | -0.1635 to 0.1814 | No | ns | >0.9999 |

#### Supplemental Table 6D - Dentate Gyrus %area ramified

| 3way ANOVA | | | | | |
|---|---|---|---|---|---|
| Source of Variation | % of total variation | P value | P value summa | Significant? | |
| Treatment | 2.729 | 0.3514 | ns | No | |
| Sex | 0.5644 | 0.6699 | ns | No | |
| Genotype | 1.535 | 0.4832 | ns | No | |
| Treatment x Sex | 0.2487 | 0.777 | ns | No | |
| Treatment x Genotype | 0.02003 | 0.9359 | ns | No | |
| Sex x Genotype | 0.5671 | 0.6692 | ns | No | |
| Treatment x Sex x Genotype | 2.912 | 0.336 | ns | No | |
| ANOVA table | SS (Type III) | DF | MS | F (DFn, DFd) | P value |
| Treatment | 0.4523 | 1 | 0.4523 | F (1, 30) = 0.8960 | P=0.3514 |
| Sex | 0.09353 | 1 | 0.09353 | F (1, 30) = 0.1853 | P=0.6699 |
| Genotype | 0.2544 | 1 | 0.2544 | F (1, 30) = 0.5040 | P=0.4832 |
| Treatment x Sex | 0.04121 | 1 | 0.04121 | F (1, 30) = 0.08165 | P=0.7770 |
| Treatment x Genotype | 0.00332 | 1 | 0.00332 | F (1, 30) = 0.00657 | P=0.9359 |
| Sex x Genotype | 0.09398 | 1 | 0.09398 | F (1, 30) = 0.1862 | P=0.6692 |
| Treatment x Sex x Genotype | 0.4826 | 1 | 0.4826 | F (1, 30) = 0.9561 | P=0.3360 |
| Residual | 15.14 | 30 | 0.5048 | | |

| Šídák's multiple comparisons test | Predicted (LS) Mean diff. | 95.00% CI of diff. | Below threshold | Summary | Adjusted P |
|---|---|---|---|---|---|
| NT:Female WT vs. TR:Female WT | -0.3712 | -1.925 to 1.182 | No | ns | 0.9995 |
| NT:Female Tg-AD vs. TR:Female Tg-AD | 0.0562 | -1.333 to 1.446 | No | ns | >0.9999 |
| NT:Male WT vs. TR:Male WT | -0.04133 | -1.595 to 1.512 | No | ns | >0.9999 |
| NT:Male Tg-AD vs. TR:Male Tg-AD | -0.546 | -1.936 to 0.8436 | No | ns | 0.9591 |
| NT:Female WT vs. NT:Female Tg-AD | -0.1473 | -1.478 to 1.183 | No | ns | >0.9999 |
| NT:Male WT vs. NT:Male Tg-AD | 0.5243 | -0.8061 to 1.855 | No | ns | 0.9582 |
| TR:Female WT vs. TR:Female Tg-AD | 0.28 | -1.325 to 1.885 | No | ns | >0.9999 |
| TR:Male WT vs. TR:Male Tg-AD | 0.01967 | -1.585 to 1.624 | No | ns | >0.9999 |
| NT:Female WT vs. NT:Male WT | -0.1652 | -1.434 to 1.103 | No | ns | >0.9999 |
| NT:Female Tg-AD vs. NT:Male Tg-AD | 0.5065 | -0.8831 to 1.896 | No | ns | 0.9766 |
| TR:Female WT vs. TR:Male WT | 0.1647 | -1.629 to 1.959 | No | ns | >0.9999 |
| TR:Female Tg-AD vs. TR:Male Tg-AD | -0.0957 | -1.485 to 1.294 | No | ns | >0.9999 |

#### Supplemental Table 6E - Dentate Gyrus %area reactive

| 3way ANOVA | | | | | |
|---|---|---|---|---|---|
| Source of Variation | % of total variation | P value | P value summary | Significant? | |
| Treatment | 2.032 | 0.1301 | ns | No | |
| Sex | 9.762 | 0.0019 | ** | Yes | |
| Genotype | 37.66 | <0.0001 | **** | Yes | |
| Treatment x Sex | 3.315 | 0.056 | ns | No | |
| Treatment x Genotype | 2.894 | 0.0731 | ns | No | |
| Sex x Genotype | 6.914 | 0.0074 | ** | Yes | |
| Treatment x Sex x Genotype | 5.498 | 0.0157 | * | Yes | |
| ANOVA table | SS (Type III) | DF | MS | F (DFn, DFd) | P value |
| Treatment | 0.02924 | 1 | 0.02924 | F (1, 30) = 2.422 | P=0.1301 |
| Sex | 0.1405 | 1 | 0.1405 | F (1, 30) = 11.64 | P=0.0019 |
| Genotype | 0.5418 | 1 | 0.5418 | F (1, 30) = 44.88 | P<0.0001 |
| Treatment x Sex | 0.0477 | 1 | 0.0477 | F (1, 30) = 3.952 | P=0.0560 |
| Treatment x Genotype | 0.04164 | 1 | 0.04164 | F (1, 30) = 3.449 | P=0.0731 |
| Sex x Genotype | 0.09948 | 1 | 0.09948 | F (1, 30) = 8.241 | P=0.0074 |
| Treatment x Sex x Genotype | 0.07911 | 1 | 0.07911 | F (1, 30) = 6.553 | P=0.0157 |
| Residual | 0.3622 | 30 | 0.01207 | | |
| Šídák's multiple comparisons test | Predicted (LS) Mean diff. | 95.00% CI of diff. | Below threshold | Summary | Adjusted P |
| NT:Female WT vs. TR:Female WT | -0.03217 | -0.2724 to 0.2081 | No | ns | >0.9999 |
| NT:Female Tg-AD vs. TR:Female Tg-AD | 0.2934 | 0.07850 to 0.5083 | Yes | ** | 0.0025 |
| NT:Male WT vs. TR:Male WT | 0.01 | -0.2303 to 0.2503 | No | ns | >0.9999 |
| NT:Male Tg-AD vs. TR:Male Tg-AD | -0.0418 | -0.2567 to 0.1731 | No | ns | >0.9999 |
| NT:Female WT vs. NT:Female Tg-AD | -0.5155 | -0.7212 to -0.3097 | Yes | **** | <0.0001 |
| NT:Male WT vs. NT:Male Tg-AD | -0.1152 | -0.3209 to 0.09055 | No | ns | 0.6926 |
| TR:Female WT vs. TR:Female Tg-AD | -0.1899 | -0.4380 to 0.05824 | No | ns | 0.2584 |
| TR:Male WT vs. TR:Male Tg-AD | -0.167 | -0.4151 to 0.08114 | No | ns | 0.432 |
| NT:Female WT vs. NT:Male WT | -0.001167 | -0.1973 to 0.1950 | No | ns | >0.9999 |
| NT:Female Tg-AD vs. NT:Male Tg-AD | 0.3991 | 0.1842 to 0.6140 | Yes | **** | <0.0001 |
| TR:Female WT vs. TR:Male WT | 0.041 | -0.2364 to 0.3184 | No | ns | >0.9999 |
| TR:Female Tg-AD vs. TR:Male Tg-AD | 0.0639 | -0.1510 to 0.2788 | No | ns | 0.9957 |

#### Supplemental Table 6F - Dentate Gyrus %area ameoboid

| 3way ANOVA | | | | | |
|---|---|---|---|---|---|
| Source of Variation | % of total variation | P value | P value summary | Significant? | |
| Treatment | 2.119 | 0.1296 | ns | No | |
| Sex | 12.67 | 0.0006 | *** | Yes | |
| Genotype | 26.57 | <0.0001 | **** | Yes | |
| Treatment x Sex | 3.238 | 0.0636 | ns | No | |
| Treatment x Genotype | 7.56 | 0.0062 | ** | Yes | |
| Sex x Genotype | 7.471 | 0.0065 | ** | Yes | |
| Treatment x Sex x Genotype | 5.604 | 0.0167 | * | Yes | |
| ANOVA table | SS (Type III) | DF | MS | F (DFn, DFd) | P value |
| Treatment | 0.0155 | 1 | 0.0155 | F (1, 30) = 2.429 | P=0.1296 |

| | | | | | |
|---|---|---|---|---|---|
| Sex | 0.09271 | 1 | 0.09271 | F (1, 30) = 14.52 | P=0.0006 |
| Genotype | 0.1944 | 1 | 0.1944 | F (1, 30) = 30.45 | P<0.0001 |
| Treatment x Sex | 0.02369 | 1 | 0.02369 | F (1, 30) = 3.712 | P=0.0636 |
| Treatment x Genotype | 0.05532 | 1 | 0.05532 | F (1, 30) = 8.666 | P=0.0062 |
| Sex x Genotype | 0.05467 | 1 | 0.05467 | F (1, 30) = 8.563 | P=0.0065 |
| Treatment x Sex x Genotype | 0.04101 | 1 | 0.04101 | F (1, 30) = 6.424 | P=0.0167 |
| Residual | 0.1915 | 30 | 0.006384 | | |
| Šídák's multiple comparisons test | Predicted (LS) Mean diff. | 95.00% CI of diff. | Below threshol | Summary | Adjusted P |
| NT:Female WT vs. TR:Female WT | -0.05342 | -0.2281 to 0.1213 | No | ns | 0.9945 |
| NT:Female Tg-AD vs. TR:Female Tg-AD | 0.2402 | 0.08393 to 0.3965 | Yes | *** | 0.0006 |
| NT:Male WT vs. TR:Male WT | -0.02083 | -0.1955 to 0.1539 | No | ns | >0.9999 |
| NT:Male Tg-AD vs. TR:Male Tg-AD | 0.0011 | -0.1552 to 0.1574 | No | ns | >0.9999 |
| NT:Female WT vs. NT:Female Tg-AD | -0.3731 | -0.5227 to -0.2235 | Yes | **** | <0.0001 |
| NT:Male WT vs. NT:Male Tg-AD | -0.08043 | -0.2300 to 0.06918 | No | ns | 0.7422 |
| TR:Female WT vs. TR:Female Tg-AD | -0.0795 | -0.2599 to 0.1009 | No | ns | 0.9118 |
| TR:Male WT vs. TR:Male Tg-AD | -0.0585 | -0.2389 to 0.1219 | No | ns | 0.9909 |
| NT:Female WT vs. NT:Male WT | 0.007417 | -0.1352 to 0.1501 | No | ns | >0.9999 |
| NT:Female Tg-AD vs. NT:Male Tg-AD | 0.3001 | 0.1438 to 0.4564 | Yes | **** | <0.0001 |
| TR:Female WT vs. TR:Male WT | 0.04 | -0.1617 to 0.2417 | No | ns | >0.9999 |
| TR:Female Tg-AD vs. TR:Male Tg-AD | 0.061 | -0.09527 to 0.2173 | No | ns | 0.9609 |

#### Supplemental Table 7A - Whole Hippocampus count/µm AT8

| 3way ANOVA | | | | | |
|---|---|---|---|---|---|
| Source of Variation | % of total variation | P value | P value sum | Significant? | |
| Treatment | 12.26 | 0.0249 | * | Yes | |
| Sex | 0.463 | 0.6502 | ns | No | |
| Genotype | 8.868 | 0.0538 | ns | No | |
| Treatment x Sex | 4.177 | 0.1788 | ns | No | |
| Treatment x Genotype | 2.378 | 0.3074 | ns | No | |
| Sex x Genotype | 1.216 | 0.4636 | ns | No | |
| Treatment x Sex x Genotype | 0.0153 | 0.9342 | ns | No | |
| ANOVA table | SS (Type III) | DF | MS | F (DFn, DFd) | P value |
| Treatment | 3.273E-07 | 1 | 3.273E-07 | F (1, 31) = 5.553 | P=0.0249 |
| Sex | 1.236E-08 | 1 | 1.236E-08 | F (1, 31) = 0.2097 | P=0.6502 |
| Genotype | 2.368E-07 | 1 | 2.368E-07 | F (1, 31) = 4.017 | P=0.0538 |
| Treatment x Sex | 1.115E-07 | 1 | 1.115E-07 | F (1, 31) = 1.892 | P=0.1788 |
| Treatment x Genotype | 6.349E-08 | 1 | 6.349E-08 | F (1, 31) = 1.077 | P=0.3074 |
| Sex x Genotype | 3.246E-08 | 1 | 3.246E-08 | F (1, 31) = 0.5507 | P=0.4636 |
| Treatment x Sex x Genotype | 4.086E-10 | 1 | 4.086E-10 | F (1, 31) = 0.00693 | P=0.9342 |
| Residual | 0.000001827 | 31 | 5.894E-08 | | |
| Šídák's multiple comparisons test | Predicted (LS) Mean di | 95.00% CI of diff. | Below thresh | Summary | Adjusted P |
| NT:Female WT vs. TR:Female WT | 0.0002209 | -0.0003260 to 0.0007677 | No | ns | 0.951 |
| NT:Female Tg-AD vs. TR:Female Tg-AD | 0.0003727 | -8.074e-005 to 0.0008261 | No | ns | 0.181 |
| NT:Male WT vs. TR:Male WT | -0.00001115 | -0.0005135 to 0.0004912 | No | ns | >0.9999 |
| NT:Male Tg-AD vs. TR:Male Tg-AD | 0.0001672 | -0.0002863 to 0.0006206 | No | ns | 0.9748 |
| NT:Female WT vs. NT:Female Tg-AD | -0.0001763 | -0.0006297 to 0.0002771 | No | ns | 0.9626 |
| NT:Male WT vs. NT:Male Tg-AD | -0.0003076 | -0.0007610 to 0.0001459 | No | ns | 0.4224 |
| TR:Female WT vs. TR:Female Tg-AD | -0.00002447 | -0.0005713 to 0.0005224 | No | ns | >0.9999 |
| TR:Male WT vs. TR:Male Tg-AD | -0.0001293 | -0.0006316 to 0.0003731 | No | ns | 0.9989 |
| NT:Female WT vs. NT:Male WT | 0.0001386 | -0.0003350 to 0.0006122 | No | ns | 0.9964 |
| NT:Female Tg-AD vs. NT:Male Tg-AD | 0.000007333 | -0.0004250 to 0.0004397 | No | ns | >0.9999 |
| TR:Female WT vs. TR:Male WT | -0.00009342 | -0.0006653 to 0.0004785 | No | ns | >0.9999 |
| TR:Female Tg-AD vs. TR:Male Tg-AD | -0.0001982 | -0.0006718 to 0.0002754 | No | ns | 0.9375 |

#### Supplemental Table 7B - Dentate Gyrus count/µm AT8

| 3way ANOVA | | | | | |
|---|---|---|---|---|---|
| Source of Variation | % of total variation | P value | P value sum | Significant? | |
| Treatment | 6.414 | 0.1208 | ns | No | |
| Sex | 0.4196 | 0.686 | ns | No | |
| Genotype | 1.788 | 0.4061 | ns | No | |
| Treatment x Sex | 12.15 | 0.0357 | * | Yes | |
| Treatment x Genotype | 0.5786 | 0.6352 | ns | No | |
| Sex x Genotype | 0.3807 | 0.7002 | ns | No | |
| Treatment x Sex x Genotype | 0.0448 | 0.8948 | ns | No | |
| ANOVA table | SS (Type III) | DF | MS | F (DFn, DFd) | P value |
| Treatment | 2.044E-07 | 1 | 2.044E-07 | F (1, 31) = 2.545 | P=0.1208 |
| Sex | 1.337E-08 | 1 | 1.337E-08 | F (1, 31) = 0.1665 | P=0.6860 |
| Genotype | 5.696E-08 | 1 | 5.696E-08 | F (1, 31) = 0.7094 | P=0.4061 |
| Treatment x Sex | 3.872E-07 | 1 | 3.872E-07 | F (1, 31) = 4.822 | P=0.0357 |
| Treatment x Genotype | 1.844E-08 | 1 | 1.844E-08 | F (1, 31) = 0.2296 | P=0.6352 |
| Sex x Genotype | 1.213E-08 | 1 | 1.213E-08 | F (1, 31) = 0.1511 | P=0.7002 |
| Treatment x Sex x Genotype | 1.427E-09 | 1 | 1.427E-09 | F (1, 31) = 0.01778 | P=0.8948 |
| Residual | 0.000002489 | 31 | 8.03E-08 | | |
| Šídák's multiple comparisons test | Predicted (LS) Mean di | 95.00% CI of diff. | Below thresh | Summary | Adjusted P |
| NT:Female WT vs. TR:Female WT | 0.0003198 | -0.0003185 to 0.0009581 | No | ns | 0.8181 |
| NT:Female Tg-AD vs. TR:Female Tg-AD | 0.000384 | -0.0001452 to 0.0009132 | No | ns | 0.3277 |
| NT:Male WT vs. TR:Male WT | -0.0001126 | -0.0006989 to 0.0004737 | No | ns | >0.9999 |
| NT:Male Tg-AD vs. TR:Male Tg-AD | 0.0000011 | -0.0005281 to 0.0005303 | No | ns | >0.9999 |
| NT:Female WT vs. NT:Female Tg-AD | -0.0000742 | -0.0006034 to 0.0004550 | No | ns | >0.9999 |
| NT:Male WT vs. NT:Male Tg-AD | -0.0001711 | -0.0007003 to 0.0003581 | No | ns | 0.9913 |

| | | | | | |
|---|---|---|---|---|---|
| TR:Female WT vs. TR:Female Tg-AD | -0.00001 | -0.0006483 to 0.0006283 | No | ns | >0.9999 |
| TR:Male WT vs. TR:Male Tg-AD | -0.0000574 | -0.0006437 to 0.0005289 | No | ns | >0.9999 |
| NT:Female WT vs. NT:Male WT | 0.0002144 | -0.0003384 to 0.0007672 | No | ns | 0.9632 |
| NT:Female Tg-AD vs. NT:Male Tg-AD | 0.0001175 | -0.0003871 to 0.0006221 | No | ns | 0.9996 |
| TR:Female WT vs. TR:Male WT | -0.000218 | -0.0008855 to 0.0004495 | No | ns | 0.9905 |
| TR:Female Tg-AD vs. TR:Male Tg-AD | -0.0002654 | -0.0008182 to 0.0002874 | No | ns | 0.8552 |
| 2way ANOVA Dentate Gyrus count/μm AT8 Female | | | | | |
| Source of Variation | % of total variation | P value | P value sum | Significant? | |
| Interaction | 0.2705 | 0.8058 | ns | No | |
| Treatment | 32.51 | 0.0151 | * | Yes | |
| Genotype | 0.4653 | 0.7473 | ns | No | |
| ANOVA table | SS (Type III) | DF | MS | F (DFn, DFd) | P value |
| Interaction | 4.58E-09 | 1 | 4.58E-09 | F (1, 15) = 0.06261 | P=0.8058 |
| Treatment | 5.504E-07 | 1 | 5.504E-07 | F (1, 15) = 7.524 | P=0.0151 |
| Genotype | 7.877E-09 | 1 | 7.877E-09 | F (1, 15) = 0.1077 | P=0.7473 |
| Residual | 0.000001097 | 15 | 7.315E-08 | | |
| 2way ANOVA Dentate Gyrus count/μm AT8 Male | | | | | |
| Source of Variation | % of total variation | P value | P value sum | Significant? | |
| Interaction | 1.061 | 0.6754 | ns | No | |
| Treatment | 1.02 | 0.6813 | ns | No | |
| Genotype | 4.285 | 0.404 | ns | No | |
| ANOVA table | SS (Type III) | DF | MS | F (DFn, DFd) | P value |
| Interaction | 1.583E-08 | 1 | 1.583E-08 | F (1, 16) = 0.1820 | P=0.6754 |
| Treatment | 1.522E-08 | 1 | 1.522E-08 | F (1, 16) = 0.1750 | P=0.6813 |
| Genotype | 6.393E-08 | 1 | 6.393E-08 | F (1, 16) = 0.7349 | P=0.4040 |
| Residual | 0.000001392 | 16 | 8.7E-08 | | |

#### Supplemental Table 7C - CA1 count/μm AT8

| | | | | | |
|---|---|---|---|---|---|
| 3way ANOVA | | | | | |
| Source of Variation | % of total variation | P value | P value sum | Significant? | |
| Treatment | 10.12 | 0.035 | * | Yes | |
| Sex | 0.3057 | 0.7041 | ns | No | |
| Genotype | 14.58 | 0.0126 | * | Yes | |
| Treatment x Sex | 5.321 | 0.1199 | ns | No | |
| Treatment x Genotype | 2.307 | 0.3004 | ns | No | |
| Sex x Genotype | 0.7488 | 0.5529 | ns | No | |
| Treatment x Sex x Genotype | 0.4193 | 0.6566 | ns | No | |
| ANOVA table | SS (Type III) | DF | MS | F (DFn, DFd) | P value |
| Treatment | 2.959E-07 | 1 | 2.959E-07 | F (1, 31) = 4.864 | P=0.0350 |
| Sex | 8.942E-09 | 1 | 8.942E-09 | F (1, 31) = 0.1470 | P=0.7041 |
| Genotype | 4.264E-07 | 1 | 4.264E-07 | F (1, 31) = 7.008 | P=0.0126 |
| Treatment x Sex | 1.556E-07 | 1 | 1.556E-07 | F (1, 31) = 2.558 | P=0.1199 |
| Treatment x Genotype | 6.749E-08 | 1 | 6.749E-08 | F (1, 31) = 1.109 | P=0.3004 |
| Sex x Genotype | 2.19E-08 | 1 | 2.19E-08 | F (1, 31) = 0.3599 | P=0.5529 |
| Treatment x Sex x Genotype | 1.226E-08 | 1 | 1.226E-08 | F (1, 31) = 0.2016 | P=0.6566 |
| Residual | 0.000001886 | 31 | 6.085E-08 | | |
| Šídák's multiple comparisons test | Predicted (LS) Mean di | 95.00% CI of diff. | | Adjusted P Value | |
| NT:Female WT vs. TR:Female WT | 0.0002586 | -0.0002970 to 0.0008142 | | 0.8786 | |
| NT:Female Tg-AD vs. TR:Female Tg-AD | 0.0003562 | -0.0001045 to 0.0008169 | | 0.2472 | |
| NT:Male WT vs. TR:Male WT | -0.0000724 | -0.0005828 to 0.0004380 | | >0.9999 | |
| NT:Male Tg-AD vs. TR:Male Tg-AD | 0.0001703 | -0.0002904 to 0.0006310 | | 0.9743 | |
| NT:Female WT vs. NT:Female Tg-AD | -0.0002142 | -0.0006749 to 0.0002465 | | 0.8792 | |
| NT:Male WT vs. NT:Male Tg-AD | -0.0003837 | -0.0008444 to 7.697e-005 | | 0.1682 | |
| TR:Female WT vs. TR:Female Tg-AD | -0.0001166 | -0.0006722 to 0.0004390 | | 0.9999 | |
| TR:Male WT vs. TR:Male Tg-AD | -0.000141 | -0.0006514 to 0.0003694 | | 0.9979 | |

| | | | | |
|---|---|---|---|---|
| NT:Female WT vs. NT:Male WT | 0.000183 | -0.0002982 to 0.0006642 | | 0.9682 |
| NT:Female Tg-AD vs. NT:Male Tg-AD | 0.0000135 | -0.0004258 to 0.0004528 | | >0.9999 |
| TR:Female WT vs. TR:Male WT | -0.000148 | -0.0007291 to 0.0004331 | | 0.999 |
| TR:Female Tg-AD vs. TR:Male Tg-AD | -0.0001724 | -0.0006536 to 0.0003088 | | 0.9798 |

#### Supplemental Table 7D - CA3 count/µm AT8

| 3way ANOVA | | | | | |
|---|---|---|---|---|---|
| Source of Variation | % of total variation | P value | P value sum | Significant? | |
| Treatment | 5.598 | 0.146 | ns | No | |
| Sex | 4.093 | 0.2117 | ns | No | |
| Genotype | 5.501 | 0.1494 | ns | No | |
| Treatment x Sex | 0.165 | 0.7996 | ns | No | |
| Treatment x Genotype | 1.339 | 0.4712 | ns | No | |
| Sex x Genotype | 2.412 | 0.3352 | ns | No | |
| Treatment x Sex x Genotype | 0.6944 | 0.6031 | ns | No | |
| ANOVA table | SS (Type III) | DF | MS | F (DFn, DFd) | P value |
| Treatment | 1.785E-07 | 1 | 1.785E-07 | F (1, 31) = 2.224 | P=0.1460 |
| Sex | 1.305E-07 | 1 | 1.305E-07 | F (1, 31) = 1.626 | P=0.2117 |
| Genotype | 1.754E-07 | 1 | 1.754E-07 | F (1, 31) = 2.186 | P=0.1494 |
| Treatment x Sex | 5.261E-09 | 1 | 5.261E-09 | F (1, 31) = 0.06555 | P=0.7996 |
| Treatment x Genotype | 4.271E-08 | 1 | 4.271E-08 | F (1, 31) = 0.5321 | P=0.4712 |
| Sex x Genotype | 7.691E-08 | 1 | 7.691E-08 | F (1, 31) = 0.9583 | P=0.3352 |
| Treatment x Sex x Genotype | 2.214E-08 | 1 | 2.214E-08 | F (1, 31) = 0.2759 | P=0.6031 |
| Residual | 0.000002488 | 31 | 8.026E-08 | | |
| Šídák's multiple comparisons test | Predicted (LS) Mean di | 95.00% CI of diff. | | Adjusted P Value | |
| NT:Female WT vs. TR:Female WT | 0.0001432 | -0.0004949 to 0.0007813 | | 0.9997 | |
| NT:Female Tg-AD vs. TR:Female Tg-AD | 0.0001811 | -0.0003480 to 0.0007102 | | 0.986 | |
| NT:Male WT vs. TR:Male WT | -0.0000018 | -0.0005880 to 0.0005844 | | >0.9999 | |
| NT:Male Tg-AD vs. TR:Male Tg-AD | 0.0002311 | -0.0002980 to 0.0007602 | | 0.9175 | |
| NT:Female WT vs. NT:Female Tg-AD | -0.0000653 | -0.0005944 to 0.0004638 | | >0.9999 | |
| NT:Male WT vs. NT:Male Tg-AD | -0.0003445 | -0.0008736 to 0.0001846 | | 0.4825 | |
| TR:Female WT vs. TR:Female Tg-AD | -0.0000274 | -0.0006655 to 0.0006107 | | >0.9999 | |
| TR:Male WT vs. TR:Male Tg-AD | -0.0001116 | -0.0006978 to 0.0004746 | | >0.9999 | |
| NT:Female WT vs. NT:Male WT | 0.000045 | -0.0005076 to 0.0005976 | | >0.9999 | |
| NT:Female Tg-AD vs. NT:Male Tg-AD | -0.0002342 | -0.0007387 to 0.0002703 | | 0.8805 | |
| TR:Female WT vs. TR:Male WT | -0.0001 | -0.0007674 to 0.0005674 | | >0.9999 | |
| TR:Female Tg-AD vs. TR:Male Tg-AD | -0.0001842 | -0.0007368 to 0.0003684 | | 0.9887 | |

#### Supplemental Table 7E - SB count/µm AT8

| 3way ANOVA | | | | | |
|---|---|---|---|---|---|
| Source of Variation | % of total variation | P value | P value sum | Significant? | |
| Treatment | 9.101 | 0.0379 | * | Yes | |
| Sex | 0.331 | 0.6821 | ns | No | |
| Genotype | 10.33 | 0.0277 | * | Yes | |
| Treatment x Sex | 6.543 | 0.0756 | ns | No | |
| Treatment x Genotype | 5.381 | 0.1055 | ns | No | |
| Sex x Genotype | 1.781 | 0.3448 | ns | No | |
| Treatment x Sex x Genotype | 1.192 | 0.4385 | ns | No | |
| ANOVA table | SS (Type III) | DF | MS | F (DFn, DFd) | P value |
| Treatment | 3.317E-07 | 1 | 3.317E-07 | F (1, 31) = 4.702 | P=0.0379 |
| Sex | 1.206E-08 | 1 | 1.206E-08 | F (1, 31) = 0.1710 | P=0.6821 |
| Genotype | 3.763E-07 | 1 | 3.763E-07 | F (1, 31) = 5.335 | P=0.0277 |
| Treatment x Sex | 2.385E-07 | 1 | 2.385E-07 | F (1, 31) = 3.381 | P=0.0756 |
| Treatment x Genotype | 1.961E-07 | 1 | 1.961E-07 | F (1, 31) = 2.780 | P=0.1055 |
| Sex x Genotype | 6.492E-08 | 1 | 6.492E-08 | F (1, 31) = 0.9204 | P=0.3448 |
| Treatment x Sex x Genotype | 4.345E-08 | 1 | 4.345E-08 | F (1, 31) = 0.6160 | P=0.4385 |
| Residual | 0.000002187 | 31 | 7.054E-08 | | |
| Šídák's multiple comparisons test | Predicted (LS) Mean di | 95.00% CI of diff. | | Adjusted P Value | |
| NT:Female WT vs. TR:Female WT | 0.0001353 | -0.0004630 to 0.0007335 | | 0.9997 | |

| | | | | |
|---|---|---|---|---|
| NT:Female Tg-AD vs. TR:Female Tg-AD | 0.0005619 | 6.590e-005 to 0.001058 | | 0.0173 |
| NT:Male WT vs. TR:Male WT | -0.0000481 | -0.0005976 to 0.0005014 | | >0.9999 |
| NT:Male Tg-AD vs. TR:Male Tg-AD | 0.0001055 | -0.0003906 to 0.0006015 | | 0.9998 |
| NT:Female WT vs. NT:Female Tg-AD | -0.0004977 | -0.0009938 to -1.702e-006 | | 0.0487 |
| NT:Male WT vs. NT:Male Tg-AD | -0.0001943 | -0.0006903 to 0.0003018 | | 0.9606 |
| TR:Female WT vs. TR:Female Tg-AD | -0.00007107 | -0.0006693 to 0.0005272 | | >0.9999 |
| TR:Male WT vs. TR:Male Tg-AD | -0.0000407 | -0.0005902 to 0.0005088 | | >0.9999 |
| NT:Female WT vs. NT:Male WT | 0.0000442 | -0.0004739 to 0.0005623 | | >0.9999 |
| NT:Female Tg-AD vs. NT:Male Tg-AD | 0.0003477 | -0.0001253 to 0.0008206 | | 0.3104 |
| TR:Female WT vs. TR:Male WT | -0.0001392 | -0.0007648 to 0.0004865 | | 0.9997 |
| TR:Female Tg-AD vs. TR:Male Tg-AD | -0.0001088 | -0.0006269 to 0.0004093 | | 0.9999 |

#### Supplemental Table 8 - Subgranular zone %area DCX

| 3way ANOVA | | | | | |
|---|---|---|---|---|---|
| Source of Variation | % of total variation | P value | P value summ | Significant? | |
| Treatment | 9.469 | 0.0428 | * | Yes | |
| Sex | 9.524 | 0.0423 | * | Yes | |
| Genotype | 10.56 | 0.0331 | * | Yes | |
| Treatment x Sex | 0.8401 | 0.5338 | ns | No | |
| Treatment x Genotype | 1.211 | 0.4557 | ns | No | |
| Sex x Genotype | 1.646 | 0.3853 | ns | No | |
| Treatment x Sex x Genotype | 0.6034 | 0.5977 | ns | No | |
| ANOVA table | SS (Type III) | DF | MS | F (DFn, DFd) | P value |
| Treatment | 1.877 | 1 | 1.877 | F (1, 31) = 4.461 | P=0.0428 |
| Sex | 1.888 | 1 | 1.888 | F (1, 31) = 4.488 | P=0.0423 |
| Genotype | 2.094 | 1 | 2.094 | F (1, 31) = 4.977 | P=0.0331 |
| Treatment x Sex | 0.1665 | 1 | 0.1665 | F (1, 31) = 0.3959 | P=0.5338 |
| Treatment x Genotype | 0.24 | 1 | 0.24 | F (1, 31) = 0.5706 | P=0.4557 |
| Sex x Genotype | 0.3262 | 1 | 0.3262 | F (1, 31) = 0.7756 | P=0.3853 |
| Treatment x Sex x Genotype | 0.1196 | 1 | 0.1196 | F (1, 31) = 0.2843 | P=0.5977 |
| Residual | 13.04 | 31 | 0.4206 | | |
| Šídák's multiple comparisons test | Predicted (LS) Mean diff. | 95.00% CI of diff. | Below thresh | Summary | Adjusted P |
| NT:Female WT vs. TR:Female WT | -0.0413 | -1.502 to 1.420 | No | ns | >0.9999 |
| NT:Female Tg-AD vs. TR:Female Tg-AD | -0.5888 | -1.854 to 0.6764 | No | ns | 0.8786 |
| NT:Male WT vs. TR:Male WT | -0.5352 | -1.826 to 0.7561 | No | ns | 0.9415 |
| NT:Male Tg-AD vs. TR:Male Tg-AD | -0.6296 | -1.841 to 0.5817 | No | ns | 0.7815 |
| NT:Female WT vs. NT:Female Tg-AD | 0.5606 | -0.7046 to 1.826 | No | ns | 0.9096 |
| NT:Male WT vs. NT:Male Tg-AD | 0.7082 | -0.5031 to 1.920 | No | ns | 0.6373 |
| TR:Female WT vs. TR:Female Tg-AD | 0.0131 | -1.448 to 1.474 | No | ns | >0.9999 |
| TR:Male WT vs. TR:Male Tg-AD | 0.6138 | -0.6774 to 1.905 | No | ns | 0.863 |
| NT:Female WT vs. NT:Male WT | 0.5099 | -0.7014 to 1.721 | No | ns | 0.9351 |
| NT:Female Tg-AD vs. NT:Male Tg-AD | 0.6575 | -0.6077 to 1.923 | No | ns | 0.7816 |
| TR:Female WT vs. TR:Male WT | 0.016 | -1.512 to 1.544 | No | ns | >0.9999 |
| TR:Female Tg-AD vs. TR:Male Tg-AD | 0.6167 | -0.5946 to 1.828 | No | ns | 0.8026 |

#### Supplemental Table 9A - Whole Hippocampus %area NeuN

| 3way ANOVA | | | | | |
|---|---|---|---|---|---|
| Source of Variation | % of total variation | P value | P value summ | Significant? | |
| Treatment | 0.5468 | 0.6636 | ns | No | |
| Sex | 0.05239 | 0.8928 | ns | No | |
| Genotype | 0.06539 | 0.8803 | ns | No | |
| Treatment x Sex | 0.1238 | 0.8359 | ns | No | |
| Treatment x Genotype | 5.85 | 0.1609 | ns | No | |
| Sex x Genotype | 4.431 | 0.2206 | ns | No | |
| Treatment x Sex x Genotype | 0.0227 | 0.9293 | ns | No | |
| ANOVA table | SS (Type III) | DF | MS | F (DFn, DFd) | P value |
| Treatment | 0.05991 | 1 | 0.05991 | F (1, 31) = 0.1929 | P=0.6636 |
| Sex | 0.005739 | 1 | 0.005739 | F (1, 31) = 0.01848 | P=0.8928 |
| Genotype | 0.007163 | 1 | 0.007163 | F (1, 31) = 0.02306 | P=0.8803 |
| Treatment x Sex | 0.01356 | 1 | 0.01356 | F (1, 31) = 0.04366 | P=0.8359 |
| Treatment x Genotype | 0.6409 | 1 | 0.6409 | F (1, 31) = 2.063 | P=0.1609 |
| Sex x Genotype | 0.4854 | 1 | 0.4854 | F (1, 31) = 1.563 | P=0.2206 |
| Treatment x Sex x Genotype | 0.002487 | 1 | 0.002487 | F (1, 31) = 0.008007 | P=0.9293 |
| Residual | 9.629 | 31 | 0.3106 | | |
| Šídák's multiple comparisons test | Predicted (LS) Mean | 95.00% CI of diff. | Below thresho | Summary | Adjusted P |
| NT:Female WT vs. TR:Female WT | -0.1276 | -1.383 to 1.128 | No | ns | >0.9999 |
| NT:Female Tg-AD vs. TR:Female Tg-A | 0.3642 | -0.6767 to 1.405 | No | ns | 0.9833 |
| NT:Male WT vs. TR:Male WT | -0.2365 | -1.390 to 0.9166 | No | ns | 0.9999 |
| NT:Male Tg-AD vs. TR:Male Tg-AD | 0.3206 | -0.7203 to 1.362 | No | ns | 0.9943 |
| NT:Female WT vs. NT:Female Tg-AD | -0.0454 | -1.086 to 0.9955 | No | ns | >0.9999 |
| NT:Male WT vs. NT:Male Tg-AD | -0.5345 | -1.575 to 0.5064 | No | ns | 0.7941 |
| TR:Female WT vs. TR:Female Tg-AD | 0.4464 | -0.8090 to 1.702 | No | ns | 0.981 |
| TR:Male WT vs. TR:Male Tg-AD | 0.02262 | -1.131 to 1.176 | No | ns | >0.9999 |
| NT:Female WT vs. NT:Male WT | 0.3075 | -0.7797 to 1.395 | No | ns | 0.9973 |
| NT:Female Tg-AD vs. NT:Male Tg-AD | -0.1816 | -1.174 to 0.8109 | No | ns | >0.9999 |
| TR:Female WT vs. TR:Male WT | 0.1985 | -1.114 to 1.511 | No | ns | >0.9999 |
| TR:Female Tg-AD vs. TR:Male Tg-AD | -0.2252 | -1.312 to 0.8620 | No | ns | 0.9999 |

#### Supplemental Table 9B - Dentate Gyrus %area NeuN

| 3way ANOVA | | | | | |
|---|---|---|---|---|---|
| Source of Variation | % of total variation | P value | P value summ | Significant? | |
| Treatment | 0.1279 | 0.8314 | ns | No | |
| Sex | 3.414 | 0.276 | ns | No | |
| Genotype | 0.902 | 0.5727 | ns | No | |
| Treatment x Sex | 0.1487 | 0.8185 | ns | No | |
| Treatment x Genotype | 6.081 | 0.1489 | ns | No | |
| Sex x Genotype | 0.8742 | 0.5787 | ns | No | |
| Treatment x Sex x Genotype | 1.979 | 0.4049 | ns | No | |
| ANOVA table | SS (Type III) | DF | MS | F (DFn, DFd) | P value |
| Treatment | 0.02484 | 1 | 0.02484 | F (1, 31) = 0.04609 | P=0.8314 |
| Sex | 0.6628 | 1 | 0.6628 | F (1, 31) = 1.230 | P=0.2760 |
| Genotype | 0.1751 | 1 | 0.1751 | F (1, 31) = 0.3250 | P=0.5727 |
| Treatment x Sex | 0.02887 | 1 | 0.02887 | F (1, 31) = 0.05357 | P=0.8185 |
| Treatment x Genotype | 1.181 | 1 | 1.181 | F (1, 31) = 2.191 | P=0.1489 |
| Sex x Genotype | 0.1697 | 1 | 0.1697 | F (1, 31) = 0.3150 | P=0.5787 |
| Treatment x Sex x Genotype | 0.3842 | 1 | 0.3842 | F (1, 31) = 0.7130 | P=0.4049 |
| Residual | 16.71 | 31 | 0.5389 | | |
| Šídák's multiple comparisons test | Predicted (LS) Mean | 95.00% CI of diff. | Below thresho | Summary | Adjusted P |
| NT:Female WT vs. TR:Female WT | -0.4517 | -2.105 to 1.202 | No | ns | 0.9981 |
| NT:Female Tg-AD vs. TR:Female Tg-A | 0.6662 | -0.7048 to 2.037 | No | ns | 0.8453 |
| NT:Male WT vs. TR:Male WT | -0.1569 | -1.676 to 1.362 | No | ns | >0.9999 |
| NT:Male Tg-AD vs. TR:Male Tg-AD | 0.1488 | -1.222 to 1.520 | No | ns | >0.9999 |
| NT:Female WT vs. NT:Female Tg-AD | -0.2869 | -1.658 to 1.084 | No | ns | 0.9999 |
| NT:Male WT vs. NT:Male Tg-AD | -0.1507 | -1.522 to 1.220 | No | ns | >0.9999 |

| | | | | | |
|---|---|---|---|---|---|
| TR:Female WT vs. TR:Female Tg-AD | 0.831 | -0.8226 to 2.484 | No | ns | 0.8153 |
| TR:Male WT vs. TR:Male Tg-AD | 0.155 | -1.364 to 1.674 | No | ns | >0.9999 |
| NT:Female WT vs. NT:Male WT | -0.2791 | -1.711 to 1.153 | No | ns | >0.9999 |
| NT:Female Tg-AD vs. NT:Male Tg-AD | -0.1429 | -1.450 to 1.164 | No | ns | >0.9999 |
| TR:Female WT vs. TR:Male WT | 0.01567 | -1.714 to 1.745 | No | ns | >0.9999 |
| TR:Female Tg-AD vs. TR:Male Tg-AD | -0.6603 | -2.092 to 0.7717 | No | ns | 0.885 |

#### Supplemental Table 9C - CA1 %area NeuN

| 3way ANOVA | | | | | |
|---|---|---|---|---|---|
| Source of Variation | % of total variation | P value | P value summ | Significant? | |
| Treatment | 0.7772 | 0.614 | ns | No | |
| Sex | 2.987 | 0.3256 | ns | No | |
| Genotype | 0.1558 | 0.821 | ns | No | |
| Treatment x Sex | 0.01706 | 0.9403 | ns | No | |
| Treatment x Genotype | 1.002 | 0.567 | ns | No | |
| Sex x Genotype | 2.324 | 0.3851 | ns | No | |
| Treatment x Sex x Genotype | 0.2401 | 0.7789 | ns | No | |
| ANOVA table | SS (Type III) | DF | MS | F (DFn, DFd) | P value |
| Treatment | 0.09162 | 1 | 0.09162 | F (1, 31) = 0.2596 | P=0.6140 |
| Sex | 0.3521 | 1 | 0.3521 | F (1, 31) = 0.9976 | P=0.3256 |
| Genotype | 0.01837 | 1 | 0.01837 | F (1, 31) = 0.05205 | P=0.8210 |
| Treatment x Sex | 0.002011 | 1 | 0.002011 | F (1, 31) = 0.005697 | P=0.9403 |
| Treatment x Genotype | 0.1181 | 1 | 0.1181 | F (1, 31) = 0.3348 | P=0.5670 |
| Sex x Genotype | 0.2739 | 1 | 0.2739 | F (1, 31) = 0.7762 | P=0.3851 |
| Treatment x Sex x Genotype | 0.0283 | 1 | 0.0283 | F (1, 31) = 0.08019 | P=0.7789 |
| Residual | 10.94 | 31 | 0.3529 | | |
| Šídák's multiple comparisons test | Predicted (LS) Mean | 95.00% CI of diff. | Adjusted P Value | | |
| NT:Female WT vs. TR:Female WT | -0.08323 | -1.421 to 1.255 | >0.9999 | | |
| NT:Female Tg-AD vs. TR:Female Tg-A | 0.2522 | -0.8574 to 1.362 | 0.9997 | | |
| NT:Male WT vs. TR:Male WT | 0.05635 | -1.173 to 1.286 | >0.9999 | | |
| NT:Male Tg-AD vs. TR:Male Tg-AD | 0.1713 | -0.9382 to 1.281 | >0.9999 | | |
| NT:Female WT vs. NT:Female Tg-AD | -0.04065 | -1.150 to 1.069 | >0.9999 | | |
| NT:Male WT vs. NT:Male Tg-AD | -0.2733 | -1.383 to 0.8362 | 0.9993 | | |
| TR:Female WT vs. TR:Female Tg-AD | 0.2947 | -1.043 to 1.633 | 0.9998 | | |
| TR:Male WT vs. TR:Male Tg-AD | -0.1584 | -1.388 to 1.071 | >0.9999 | | |
| NT:Female WT vs. NT:Male WT | 0.296 | -0.8629 to 1.455 | 0.999 | | |
| NT:Female Tg-AD vs. NT:Male Tg-AD | 0.06333 | -0.9946 to 1.121 | >0.9999 | | |
| TR:Female WT vs. TR:Male WT | 0.4356 | -0.9639 to 1.835 | 0.9937 | | |
| TR:Female Tg-AD vs. TR:Male Tg-AD | -0.0175 | -1.176 to 1.141 | >0.9999 | | |

#### Supplemental Table 9D - CA3 %area NeuN

| 3way ANOVA | | | | | |
|---|---|---|---|---|---|
| Source of Variation | % of total variation | P value | P value summ | Significant? | |
| Treatment | 1.76 | 0.4232 | ns | No | |
| Sex | 5.841 | 0.1493 | ns | No | |
| Genotype | 4.18 | 0.2204 | ns | No | |
| Treatment x Sex | 0.3315 | 0.7271 | ns | No | |
| Treatment x Genotype | 3.282 | 0.2762 | ns | No | |
| Sex x Genotype | 1.571 | 0.4489 | ns | No | |
| Treatment x Sex x Genotype | 1.071 | 0.5312 | ns | No | |
| ANOVA table | SS (Type III) | DF | MS | F (DFn, DFd) | P value |
| Treatment | 0.1631 | 1 | 0.1631 | F (1, 31) = 0.6586 | P=0.4232 |
| Sex | 0.5415 | 1 | 0.5415 | F (1, 31) = 2.186 | P=0.1493 |
| Genotype | 0.3875 | 1 | 0.3875 | F (1, 31) = 1.564 | P=0.2204 |
| Treatment x Sex | 0.03073 | 1 | 0.03073 | F (1, 31) = 0.1241 | P=0.7271 |
| Treatment x Genotype | 0.3043 | 1 | 0.3043 | F (1, 31) = 1.228 | P=0.2762 |
| Sex x Genotype | 0.1457 | 1 | 0.1457 | F (1, 31) = 0.5881 | P=0.4489 |
| Treatment x Sex x Genotype | 0.09933 | 1 | 0.09933 | F (1, 31) = 0.4010 | P=0.5312 |
| Residual | 7.678 | 31 | 0.2477 | | |

| Šídák's multiple comparisons test | Predicted (LS) Mean | 95.00% CI of diff. | Adjusted P Value |
|---|---|---|---|
| NT:Female WT vs. TR:Female WT | -0.4736 | -1.595 to 0.6474 | 0.9334 |
| NT:Female Tg-AD vs. TR:Female Tg-A | 0.0942 | -0.8353 to 1.024 | >0.9999 |
| NT:Male WT vs. TR:Male WT | -0.1523 | -1.182 to 0.8774 | >0.9999 |
| NT:Male Tg-AD vs. TR:Male Tg-AD | 0.002567 | -0.9270 to 0.9321 | >0.9999 |
| NT:Female WT vs. NT:Female Tg-AD | -0.3628 | -1.292 to 0.5667 | 0.9615 |
| NT:Male WT vs. NT:Male Tg-AD | -0.4064 | -1.336 to 0.5232 | 0.917 |
| TR:Female WT vs. TR:Female Tg-AD | 0.205 | -0.9160 to 1.326 | >0.9999 |
| TR:Male WT vs. TR:Male Tg-AD | -0.2515 | -1.281 to 0.7783 | 0.9993 |
| NT:Female WT vs. NT:Male WT | 0.2054 | -0.7655 to 1.176 | 0.9998 |
| NT:Female Tg-AD vs. NT:Male Tg-AD | 0.1618 | -0.7244 to 1.048 | >0.9999 |
| TR:Female WT vs. TR:Male WT | 0.5267 | -0.6457 to 1.699 | 0.9016 |
| TR:Female Tg-AD vs. TR:Male Tg-AD | 0.0702 | -0.9007 to 1.041 | >0.9999 |

#### Supplemental Table 9E - SB %area NeuN

| 3way ANOVA | | | | | |
|---|---|---|---|---|---|
| Source of Variation | % of total variation | P value | P value summ | Significant? | |
| Treatment | 4.867 | 0.1701 | ns | No | |
| Sex | 0.263 | 0.7462 | ns | No | |
| Genotype | 0.4624 | 0.6681 | ns | No | |
| Treatment x Sex | 7.747 | 0.0862 | ns | No | |
| Treatment x Genotype | 3.109 | 0.2702 | ns | No | |
| Sex x Genotype | 2.112 | 0.362 | ns | No | |
| Treatment x Sex x Genotype | 3.771 | 0.2256 | ns | No | |
| ANOVA table | SS (Type III) | DF | MS | F (DFn, DFd) | P value |
| Treatment | 1.042 | 1 | 1.042 | F (1, 31) = 1.973 | P=0.1701 |
| Sex | 0.05634 | 1 | 0.05634 | F (1, 31) = 0.1066 | P=0.7462 |
| Genotype | 0.09904 | 1 | 0.09904 | F (1, 31) = 0.1874 | P=0.6681 |
| Treatment x Sex | 1.659 | 1 | 1.659 | F (1, 31) = 3.140 | P=0.0862 |
| Treatment x Genotype | 0.666 | 1 | 0.666 | F (1, 31) = 1.260 | P=0.2702 |
| Sex x Genotype | 0.4523 | 1 | 0.4523 | F (1, 31) = 0.8559 | P=0.3620 |
| Treatment x Sex x Genotype | 0.8078 | 1 | 0.8078 | F (1, 31) = 1.529 | P=0.2256 |
| Residual | 16.38 | 31 | 0.5285 | | |
| Šídák's multiple comparisons test | Predicted (LS) Mean | 95.00% CI of diff. | Adjusted P Value | | |
| NT:Female WT vs. TR:Female WT | 0.7835 | -0.8540 to 2.421 | 0.858 | | |
| NT:Female Tg-AD vs. TR:Female Tg-A | 0.7293 | -0.6284 to 2.087 | 0.7451 | | |
| NT:Male WT vs. TR:Male WT | -0.6492 | -2.153 to 0.8549 | 0.9235 | | |
| NT:Male Tg-AD vs. TR:Male Tg-AD | 0.4742 | -0.8835 to 1.832 | 0.9835 | | |
| NT:Female WT vs. NT:Female Tg-AD | 0.1443 | -1.213 to 1.502 | >0.9999 | | |
| NT:Male WT vs. NT:Male Tg-AD | -0.8851 | -2.243 to 0.4726 | 0.4806 | | |
| TR:Female WT vs. TR:Female Tg-AD | 0.09013 | -1.547 to 1.728 | >0.9999 | | |
| TR:Male WT vs. TR:Male Tg-AD | 0.2383 | -1.266 to 1.742 | >0.9999 | | |
| NT:Female WT vs. NT:Male WT | 0.8589 | -0.5592 to 2.277 | 0.588 | | |
| NT:Female Tg-AD vs. NT:Male Tg-AD | -0.1705 | -1.465 to 1.124 | >0.9999 | | |
| TR:Female WT vs. TR:Male WT | -0.5738 | -2.286 to 1.139 | 0.9882 | | |
| TR:Female Tg-AD vs. TR:Male Tg-AD | -0.4256 | -1.844 to 0.9925 | 0.9954 | | |

**Supplemental Table 10A - Differentally Expressed Genes in Female Tg-AD rats treated with AGO**

| Gene | log2FoldChange | *P* value |
|---|---|---|
| Naa11 | 3.6418 | 0.018221 |
| Hbb-bs | 3.568193 | 0.007029 |
| LOC691215 | 3.136311 | 0.022444 |
| LOC100134871 | 3.098112 | 0.008248 |
| Gucy1b2 | 3.021704 | 0.000288 |
| Creb1.1 | 3.002541 | 0.036845 |
| Bco1 | 2.431867 | 0.024155 |
| Hba1 | 2.427623 | 0.006272 |
| Hbb | 2.122749 | 0.004627 |
| Hbb-b1 | 2.016862 | 0.030811 |
| Pde4d.2 | 1.99291 | 0.015084 |
| Cd36 | 1.934571 | 0.042578 |
| Gpihbp1 | 1.887134 | 0.034068 |
| Rhbdf2.1 | 1.689552 | 0.003329 |
| Igf1 | 1.687379 | 0.046198 |
| Ggact | 1.560284 | 0.00351 |
| Ahr.1 | 1.544474 | 0.038342 |
| Acp5.1 | 1.375113 | 0.016034 |
| H19 | 1.236545 | 0.002517 |
| Pde4c | 1.090921 | 0.030767 |
| Slc4a7 | 1.074132 | 3.46E-05 |
| Dhrs9 | 1.064596 | 0.043619 |
| Tpm1.7 | 1.055549 | 0.019825 |
| Fgfr2.1 | -1.01297 | 0.024717 |
| Mns1 | -1.02898 | 0.001521 |
| Zmynd10 | -1.03023 | 0.000963 |
| Krt71 | -1.0879 | 0.010666 |
| Ttc25 | -1.09165 | 0.004614 |
| Spag8 | -1.09688 | 0.027003 |
| Zim1 | -1.09808 | 0.034513 |
| Ddit3.1 | -1.1004 | 0.002172 |
| Drc7 | -1.13962 | 0.030827 |
| Ccdc40 | -1.14054 | 0.005877 |
| Cfap161 | -1.1538 | 0.028892 |
| Ptch2 | -1.15429 | 0.023882 |
| Tsnaxip1 | -1.16057 | 0.025128 |
| Adamtsl5 | -1.16154 | 0.035168 |
| Dnmt3a | -1.18646 | 0.005284 |
| Impg2 | -1.18745 | 0.026548 |
| Trpv2.1 | -1.19655 | 0.028527 |
| LOC682102 | -1.19836 | 0.019281 |
| Ak7 | -1.19841 | 0.000538 |

| | | |
|---|---|---|
| Rab11fip1 | -1.20455 | 0.021651 |
| Ccdc114 | -1.2401 | 0.015148 |
| Abcg5 | -1.2446 | 0.046741 |
| LOC498675 | -1.2473 | 0.001016 |
| Krt18 | -1.25909 | 0.033999 |
| Sgcg | -1.28197 | 0.019349 |
| Dnah1 | -1.29662 | 0.003511 |
| Cav3 | -1.30453 | 0.041913 |
| Gabrq | -1.31812 | 0.010705 |
| Capsl | -1.3287 | 0.010487 |
| Tmc4 | -1.36232 | 0.045608 |
| Krt8 | -1.37278 | 0.008598 |
| Mlf1 | -1.38312 | 0.003582 |
| Crabp2 | -1.44253 | 0.006289 |
| Myoc | -1.45437 | 0.022517 |
| Otc | -1.4612 | 0.049032 |
| Nudt17 | -1.46598 | 0.003417 |
| Lrrc34 | -1.47484 | 0.000812 |
| Trpv4 | -1.48535 | 0.039638 |
| Spint2 | -1.50778 | 0.011891 |
| Prrt2.1 | -1.55959 | 0.030309 |
| Kcnk16 | -1.56247 | 0.020872 |
| Glra1 | -1.64137 | 0.019226 |
| Acp5 | -1.64634 | 0.045947 |
| Adgrg2.2 | -1.67352 | 0.002381 |
| Cdh3 | -1.68942 | 0.038179 |
| Ccdc60 | -1.74091 | 0.035407 |
| P2rx3.1 | -1.74349 | 0.016935 |
| Wdr63 | -1.76697 | 0.00046 |
| Crhr2 | -1.7898 | 0.00562 |
| Slc22a14 | -1.95154 | 0.03696 |
| Scn5a.1 | -1.9689 | 0.007445 |
| Myo5c | -2.11586 | 0.001262 |
| Pih1d3 | -2.14051 | 0.025169 |
| Efhb | -2.20704 | 0.039218 |
| Sinhcaf | -2.25247 | 0.002926 |
| Aox3 | -2.2558 | 0.014411 |
| Sln | -2.26915 | 0.037361 |
| Pon3 | -2.31951 | 0.021119 |
| Calcr | -2.33344 | 0.01391 |
| Gpr15 | -2.44496 | 0.044579 |
| Tnni3 | -2.45093 | 0.032483 |
| Steap1 | -2.57184 | 0.00565 |
| Mlnr | -2.88955 | 0.020231 |

| | | |
|---|---|---|
| Trpv5 | -3.11922 | 0.025595 |
| Lypd8 | -3.21523 | 0.018821 |
| Slc47a1 | -3.25242 | 0.004339 |
| P2ry10 | -3.33881 | 0.044078 |
| Mab21l3 | -3.5105 | 0.026512 |
| RT1-T18 | -3.64226 | 0.003332 |
| Zscan10 | -4.46788 | 0.011567 |
| Csnk2b | -4.67693 | 0.03193 |

**Supplemental Table 10B - Differentally Expressed Genes in Male Tg-AD rats treated with AGO**

| Gene | log2FoldChange | *P* value |
|---|---|---|
| Nrg1.3 | 4.979004 | 0.002841638 |
| Esrp1 | 3.420768 | 0.031729875 |
| Hbb-b1 | 3.103274 | 0.025211474 |
| Dctd.1 | 3.005957 | 7.07E-05 |
| Efemp2 | 2.600966 | 0.042354273 |
| Pde4d.2 | 2.281172 | 0.005881056 |
| Baat | 2.21419 | 0.013337243 |
| Setsip | 2.160852 | 0.025275802 |
| Hba1 | 2.124446 | 0.001121112 |
| Chat | 1.959487 | 0.013441763 |
| Synpo2l | 1.879613 | 0.012390367 |
| Syce1 | 1.871352 | 0.044683749 |
| RGD1562890 | 1.609721 | 0.003176865 |
| Cartpt | 1.60888 | 0.022321277 |
| P2rx3.1 | 1.59654 | 0.017693845 |
| Cd3e | 1.556442 | 0.049766045 |
| Ceacam1.1 | 1.484991 | 0.012154195 |
| Npas4 | 1.441926 | 5.90E-11 |
| Hbb | 1.211209 | 0.039612474 |
| Cftr | 1.12859 | 0.026340721 |
| Adgrl2.2 | 1.122497 | 0.000234154 |
| Kcnip1.2 | 1.049981 | 0.018839645 |
| Maged2 | 1.014608 | 0.000794243 |
| Armc3 | -1.02592 | 0.012515487 |
| Patz1 | -1.09289 | 2.25E-05 |
| Optc | -1.21717 | 0.028018622 |
| Slc26a7 | -1.2738 | 0.041633704 |
| Sgcg | -1.27523 | 0.008492159 |
| Slc22a12 | -1.36829 | 0.011013265 |
| Odf1 | -1.44337 | 0.044462732 |
| Tmem116 | -1.53064 | 0.042588157 |
| Otx2 | -1.56981 | 0.038762209 |
| Otx2.1 | -1.56981 | 0.038762209 |

| | | |
|---|---|---|
| Vcan.3 | -1.64545 | 0.028151072 |
| Aox3 | -1.65913 | 0.033212559 |
| Wdr72 | -1.6847 | 0.0016328 |
| Mpzl2 | -1.73044 | 0.014658657 |
| Gnat2 | -1.75626 | 0.034418544 |
| Pla2g5 | -1.75728 | 0.007693307 |
| RGD1561795 | -1.95212 | 0.043640106 |
| Scube3 | -2.02558 | 0.038543587 |
| Crem.3 | -2.05769 | 0.023823492 |
| RT1-CE11 | -2.0947 | 0.028391207 |
| Phf11b | -2.15275 | 0.047178355 |
| Patz1.4 | -2.20237 | 0.000175894 |
| Steap1 | -2.24253 | 0.026080601 |
| Lbp | -2.29331 | 0.023507911 |
| Slc22a3 | -2.30148 | 0.011917916 |
| Esm1 | -2.31291 | 0.049969812 |
| Mab21l2 | -2.39362 | 0.025364437 |
| Xkr9 | -2.40397 | 0.038080764 |
| Plac8 | -2.40981 | 0.015640113 |
| Fblim1 | -2.50027 | 0.033384948 |
| Six1 | -2.66615 | 0.037698175 |
| Phox2a | -2.73158 | 0.041666987 |
| Ceacam9 | -2.75434 | 0.02428166 |
| Itgb6 | -2.75931 | 0.019589749 |
| Ces1d | -3.0493 | 0.021903299 |
| LOC100911796 | -3.08299 | 0.019558604 |
| LOC100911796.1 | -3.08299 | 0.019558604 |
| Adora3 | -3.08299 | 0.019558604 |
| Adora3.1 | -3.08299 | 0.019558604 |
| Crygs | -3.57932 | 0.042668025 |
| Tmem45al | -4.0501 | 0.035124945 |
| Phlda2 | -4.66227 | 0.010811676 |
| Pcp4 | -4.84214 | 2.78E-05 |

**Supplemental Table 11A - Positively enriched DEGs shared across all AD mutant iPSC derived neurons**

| |
|---|
| RUNX3 |
| WNT7B |
| HOXD-AS2 |
| LINC02511 |
| IGFBP5 |
| HOXB6 |
| TGFB2 |
| PDZRN4 |
| PVALB |
| NTF3 |
| MYO1E |
| COL25A1 |
| CSGALNACT1 |
| CNMD |
| CALB1 |
| PREX2 |
| CXCL12 |
| HSPB8 |
| RASGRP1 |
| GRIK1 |
| CACNG3 |
| CEMIP |
| HOXB8 |
| HOXB5 |
| SLC24A2 |
| TRPC4 |
| IFITM2 |
| KCNJ5 |
| CCKBR |
| SYNPR |
| FHL2 |
| RELN |
| CRTAC1 |
| HOXB7 |
| FAM184B |
| ANOS1 |
| GNG12 |
| IGFBP2 |
| LINC00960 |
| PLD6 |
| TMEM132B |
| TNR |
| UST |

| |
|---|
| LIX1 |
| PCDH8 |
| MOXD1 |
| HOXA1 |
| GABRA2 |
| STEAP3 |
| AC109466.1 |
| AP001803.2 |
| HOTAIRM1 |
| FAM81A |
| KCNJ2 |
| TCEAL6 |
| C8orf34 |
| YBX3 |
| SDC2 |
| HOXB3 |
| MIR1-1HG-AS1 |
| WFDC2 |
| KCNH1 |
| LUZP2 |
| ETV1 |
| EZR |
| CNTN2 |
| AC090627.1 |
| VEGFC |
| CHRM3 |
| LINC01018 |
| VAX2 |
| SLC17A6 |
| HOXB2 |
| MPP6 |
| TANC1 |
| BMPER |
| BBS9 |
| TSHZ2 |
| NANOS1 |
| SGPP2 |
| PTN |
| FSTL4 |
| SERPINE2 |
| ARRDC4 |
| TIMP1 |
| C3orf70 |
| SPTB |

| |
|---|
| ANO4 |
| TENM2 |
| RGS3 |
| TOX2 |
| LRRN1 |
| CLSTN2 |
| GABRA5 |
| PDLIM1 |
| NENF |
| MAB21L2 |
| PLEKHG3 |
| PLCB4 |
| NELL2 |
| RAPGEF4 |
| ZDHHC2 |

**Supplemental Table 11B - Negatively enriched DEGs shared across all AD mutant iPSC derived neurons**

| |
|---|
| KCNJ3 |
| EPHA4 |
| PHGDH |
| AC092376.2 |
| RPS6KA2 |
| ELFN1 |
| PHACTR3 |
| ASTN2 |
| LRRC4C |
| CPNE8 |
| AJAP1 |
| DCC |
| LANCL3 |
| NTM |
| LINC00682 |
| SSTR2 |
| CHAC1 |
| AC021504.1 |
| DHRS2 |
| CEBPB |
| CYP27C1 |
| SLIT2 |
| NXPH1 |
| NKAIN2 |
| SESN2 |
| SCGN |
| TRIM29 |

| |
|---|
| GABRG3 |
| PLXDC2 |
| RGMA |
| MET |
| CNTNAP2 |
| LHX4 |
| AC009414.2 |
| FAM46A |
| SLIT3 |
| HTR3A |
| LINC00643 |
| NKAIN4 |
| DDIT4 |
| OPN3 |
| PCK2 |
| GYPC |
| KCNS1 |
| HUNK |
| WIPF3 |
| ISLR2 |
| KCNQ3 |
| CPNE2 |
| SYT17 |
| TMEM255B |
| GFRA3 |
| LMO2 |
| AC025171.1 |
| EBF2 |
| SLC1A5 |
| AC104041.1 |
| CDK14 |
| TRIP10 |
| FAM181B |
| INPP4B |
| VEGFA |
| PHOX2B |
| PELI2 |
| HS6ST3 |
| INSM1 |
| SEMA5B |
| RASEF |
| BMP2 |
| LINC01630 |
| ADM2 |

| |
|---|
| PLCD3 |
| ZNF503-AS2 |
| PTGFR |
| HS3ST3B1 |
| TLL2 |
| PHOX2A |
| ARHGAP24 |
| LHX1-DT |
| LMO1 |
| MCUB |
| SLC6A2 |
| DMRTA1 |
| ONECUT3 |
| HS3ST4 |
| FRMPD1 |
| SOX15 |
| KLF9 |
| EPB41L2 |
| MAL2 |
| LMO3 |
| AC008875.1 |
| ZNF503 |
| HMX1 |
| SMOC1 |
| MT1F |
| LHX1 |
| UNCX |
| LRRC3B |
| TCF7L2 |
| SLC7A11 |
| SOX1 |
| CCK |
| AC010931.2 |
| AC141928.1 |
| CNTN5 |
| SPATA8 |
| KLHL1 |
| OPRK1 |
| JDP2 |
| SYT6 |
| SLC9A3R1 |
| TMCC3 |
| ARPP21 |
| PIK3AP1 |

| |
|---|
| HMOX1 |
| THBS2 |
| NR5A2 |
| TFAP2A |
| STUM |
| AC078881.1 |
| AL513318.2 |
| TMEM176A |
| MYCT1 |
| LEF1 |
| RAB37 |
| GRP |
| SATB2 |
| FRMD7 |
| SERTM1 |
| PARM1 |
| KIF25 |
| SATB2-AS1 |
| SYNPO |
| PAPPA2 |
| MYPN |
| QRFPR |
| PITX2 |
| KLHL14 |
| LINC02506 |
| TBX1 |
| B3GNT7 |

**Supplemental Table 12 - DEGs in PSEN2$^{N141}$ neurons with AGO treatment starting at D14**

| Positively enriched DEGs | Negatively enriched DEGs |
|---|---|
| RAB13 | SCN1A |
| AC092139.3 | SERPINI1 |
| WASH3P | NUP153 |
| AC015813.5 | SLC2A1 |
| AC092747.4 | APOE |
| GAD2 | LMBRD1 |
| AC026741.1 | ANKRD50 |
| AL021807.1 | TRIM35 |
| AC044849.1 | NHLRC2 |
| BAALC-AS1 | EPHA5 |
| LINC00630 | CRHBP |
| AC025754.2 | PAPD4 |
| KLF3-AS1 | MDGA2 |
| CARNMT1 | AC026790.1 |
| LINC02381 | BMP8A |
| ZFYVE26 | CCDC175 |
| KLF4 | AL583856.2 |
| ZNF586 | AL671710.1 |
| PAPPA2 | PAX6 |
| SRP14-AS1 | |
| TARBP1 | |
| AC022613.1 | |
| CALB2 | |
| TRAF3IP2-AS1 | |
| TMEM107 | |
| MAFA | |
| CLEC7A | |
| EP400P1 | |
| MCTP1 | |
| TXNIP | |

**Supplemental Table 13 - DEGs in APPV$^{717I}$ neurons with AGO treatment starting at D14**

| Positively enriched DEGs | Negatively enriched DEGs |
|---|---|
| TNC | GRIN1 |
| BBS10 | PLXNC1 |
| CCK | ADGRB2 |
| ERBB3 | APLP2 |
| ZFP42 | CACNA2D2 |
| AC018638.7 | C1QTNF4 |
| MATN3 | TNS1 |
| RBPMS | SCN3A |
| LAYN | NIPAL2 |
| IL12A | ADGRB1 |
| CLEC7A | CYP26B1 |
| RPS4XP3 | SLC22A17 |
| C11orf70 | MID1 |
| MYZAP | DPP6 |
| FAM114A1 | UNC80 |
| PSMB8 | PRSS12 |
| PRORSD1P | ANOS1 |
| ANXA3 | SLC7A8 |
| AL109811.2 | PTPRO |
| F11R | AC011008.2 |
| SCN10A | HR |
| OR7A8P | GEMIN4 |
| KCNK2 | TTLL4 |
| SCGN | RELN |
| B3GNT5 | PTPRT |
| LINC01630 | CLCN4 |
| PAPD4 | KIAA1549L |
| TRPC5 | SERPINI1 |
| STAC | NPTXR |
| VASN | MOXD1 |
| AKAIN1 | PCDHGB6 |
| C15orf65 | CDH7 |
| ZNF485 | ECEL1 |
| ZBED8 | SLC24A3 |
| TPD52L1 | EPHB6 |
| TMEM64 | GLG1 |
| PMAIP1 | PCDHAC2 |
| RPS7P1 | ATP2B4 |
| AC106869.1 | LRRN2 |
| SMCO4 | NCAN |
| NECTIN3 | SPTB |
| KYAT3 | CARTPT |

| | |
|---|---|
| CCNG1 | FKBP10 |
| TBCEL | VASH1-AS1 |
| RNF138 | ZNF774 |
| LYPLA1 | ATP1B2 |
| RPL24P8 | ADCYAP1R1 |
| WDFY1 | CCM2L |
| CD163L1 | LDLRAD4 |
| VWC2 | PCSK1 |
| ADAM12 | ANKS6 |
| LAMA4 | RASGRP1 |
| ADO | ATP13A1 |
| NUDT21 | PCDH11X |
| CDK6 | GFRA1 |
| PTPRD-AS1 | SLC24A2 |
| RPL26P19 | IGFBPL1 |
| SALL4 | RXRG |
| RPE | CALN1 |
| RASEF | SHROOM2 |
| SMIM3 | AC092111.1 |
| GMFB | KIAA1324 |
| FABP3 | FAM184B |
| SNCA | KCNJ5 |
| AC021242.3 | GRIN2D |
| AC120057.3 | GHDC |
| C6orf141 | AP003355.2 |
| MINCR | FBXL22 |
| MOSPD2 | SARNP |
| AC078845.1 | C2orf92 |
| RAP2A | SCN1A |
| C17orf49 | AL117339.4 |
| IER3-AS1 | KCNH3 |
| RAB32 | LGI1 |
| DYNC2LI1 | SORL1 |
| PCGF5 | AL645608.3 |
| HPS3 | GJD3 |
| PNRC2 | AL596220.1 |
| AC074183.2 | LINC01522 |
| KITLG | AL118558.1 |
| INSIG1 | ADAMTS18 |
| COMMD8 | AC009506.1 |
| SDCBP | MXRA5 |
| CSRP2 | AL109930.1 |
| AC099343.2 | DACT2 |
| AMD1 | ADAM8 |

| | |
|---|---|
| CALB2 | ENPP2 |
| PRUNE2 | SGPP2 |
| AL024508.2 | EPPK1 |
| PPP1CC | ASAP1-IT2 |
| SPRY2 | TBC1D8-AS1 |
| PKIB | AL359183.1 |
| GNG5 | SVOPL |
| AC080013.1 | CNTNAP4 |
| SLC9A3R1 | POU5F2 |
| MINPP1 | AC107398.3 |
| TMEM52 | TTYH1 |
| KLF4 | LINC01102 |
| UHRF2 | DNAH6 |
| ZNF569 | EDNRA |
| SYT6 | NR2F1 |
| AP002387.2 | UBE2C |
| BMI1 | BDKRB2 |
| KLHL2 | AL162390.1 |

**Supplemental Table 14 - DEGs in PSEN2$^{N141}$ neurons with AGO treatment starting at D21**

| Positively enriched DEGs | Negatively enriched DEGs |
|---|---|
| LINC02202 | GLIPR2 |
| AL031595.3 | HIST2H2BE |
| AC067930.6 | ANXA5 |
| SUCNR1 | FUBP3 |
| NA | LINC00963 |
| FEZF1 | AC027682.6 |
| AL023806.1 | NPPC |
| ENTPD1 | HOXA5 |
| AC135507.1 | GAS2L1 |
| ZDHHC11 | POLR2M |
| PAX5 | RHOU |
| RASGRF2 | PKNOX2 |
| LINC02035 | CDK6 |
| MGARP | RAB32 |
| CALB2 | GNG5 |
| TRIM66 | ILDR2 |
| MT-ND5 | LGALS1 |
| ADCY2 | ANXA11 |
| MT-ND2 | TUBB6 |
| SCGN | SYNPO |
| MT-CO3 | CALD1 |
| MTCO3P12 | LMCD1 |
| MT-ND4L | FILIP1L |
| MT-CO1 | LIMA1 |
| MT-ATP6 | B2M |
| MT-CYB | WLS |
| MT-ND1 | CROT |
| MT-CO2 | RAB34 |
| LRRC45 | BTG2 |
| LRRC55 | FSTL1 |
| MT-ND4 | CTNNAL1 |
| SHPK | NACC2 |
| CHPF2 | LITAF |
| TYW1 | FAM43A |
| PLXNA2 | HMGB2 |
| MAN2A2 | WWC3 |
| TMTC2 | TLN1 |
| SIAH3 | CCND1 |
| MDGA1 | CD99 |
|  | BCL6 |
|  | ACTN1 |
|  | CRH |

| | |
|---|---|
| | MFGE8 |
| | FBN1 |
| | COL4A1 |
| | IL1RAP |
| | PDLIM4 |
| | SH3BP4 |
| | FLNA |
| | NPPB |
| | DRAM1 |
| | MORC4 |
| | FBN2 |
| | SMAD9 |
| | NUP37 |
| | CRISPLD1 |
| | FLNC |
| | SHROOM3 |
| | IFITM2 |
| | C1orf226 |
| | LINC02381 |
| | AR |
| | PMP22 |
| | SOCS3 |
| | RRM2 |
| | CXCL12 |
| | CCL2 |
| | DLK1 |
| | SOCS1 |
| | EPHA2 |
| | MCUB |
| | SOX2 |
| | LRIG1 |
| | A2M |
| | CTGF |
| | ADAMTS9 |
| | SPARC |
| | PLAU |
| | ID1 |
| | TPM2 |
| | GNG11 |
| | BUB1B |
| | TRIM56 |
| | SNHG18 |
| | TCF7L1 |
| | TGFBI |

| | |
|---|---|
| | TUBA1C |
| | TFPI |
| | ITGA8 |
| | TOP2A |
| | ZIC1 |
| | S100B |
| | HES1 |
| | AQP1 |
| | CDC42EP1 |
| | KLHDC1 |
| | PEBP1P2 |
| | GPX8 |
| | ID3 |
| | S100A11 |
| | CFI |
| | PLAT |
| | CYR61 |
| | ADA |
| | WWTR1 |
| | HOXA4 |
| | PLS3 |
| | RGS5 |
| | SMAD3 |
| | PDLIM3 |
| | PLIN2 |
| | FAM84B |
| | LMOD1 |
| | IFITM3 |
| | BIRC5 |
| | HOXA3 |
| | FCGRT |
| | SYDE1 |
| | NEDD9 |
| | SERPINE1 |
| | GLIPR1 |
| | CDKN1A |
| | TNS3 |
| | CNN2 |
| | IGFBP7 |
| | KRT18 |
| | TNFRSF12A |
| | CDC42EP5 |
| | FGFRL1 |
| | MYL9 |

| | |
|---|---|
| | TGIF2 |
| | TAGLN |
| | ACTA2 |
| | COL1A1 |
| | COL11A1 |
| | MSX1 |
| | COL5A1 |
| | COL5A2 |
| | FAM129A |
| | CAVIN1 |
| | COL8A1 |
| | KRT8 |
| | AHNAK |
| | FN1 |

**Supplemental Table 15 - DEGs in APPV$^{717I}$ neurons with AGO treatment starting at D21**

| Positively enriched DEGs | Negatively enriched DEGs |
|---|---|
| CLEC7A | RALYL |
| PROKR1 | DRAM2 |
| AC107294.1 | FAM184B |
| FGF1 | RXRG |
| OR7C1 | TTLL3 |
| AL031432.4 | AC144831.1 |
| CMYA5 | THAP10 |
| KLB | AC100786.1 |
| CCDC178 | SMAD9 |
| GJA1 | C10orf95 |
| THRB | AP003355.2 |
| AC005224.4 | ZFP3 |
| AC040970.1 | LINC00242 |
| ALK | GOLGA6L9 |
| B3GALT1 | ZNF426-DT |
| FABP5P7 | POU4F3 |
| KCNK2 | SNCAIP |
| STOM | TOX3 |
| SCGN | GOLT1A |
| SLC9B1 | FAM185A |
| INPPL1 | AL133338.1 |
| AC012020.1 | UACA |
| KITLG | PDIA2 |
| KCNQ5 | RASL10A |
| EPHA4 | RPL35P2 |
| LRP1B | DACT2 |
| SHISA9 | AC093503.2 |
| AC099568.2 | SMTN |
| PCDHA5 | AL136295.6 |
| PAPPA2 | CTSK |
| AC007238.1 | AC000068.1 |
| GALNT18 | MICAL2 |
| PCDHA10 | HOXB-AS1 |
| UTS2 | CDKN2B |
| TENM1 | IFITM2 |
| RPH3A | SPARC |
| GPR83 | PAN3 |
| CADPS2 | OAF |
| DCC | AC118553.2 |
| TGFB2 | NR2F2 |
| GLP1R | ITGA1 |
| PHGDH | ENPP2 |

| | |
|---|---|
| SULF2 | WWTR1 |
| C6orf141 | TTYH1 |
| CALB2 | IGFBP7 |
| UNC5B | ABCA9 |
| SERPINE2 | PDLIM3 |
| GAS2L3 | PNPLA7 |
| SEMA3E | SORBS3 |
| PAM | RGS5 |
| SLC1A1 | TDRD6 |
| LPCAT2 | MDFI |
| EDIL3 | FREM1 |
| ARSK | AL356535.1 |
| LANCL3 | AC004494.1 |
| NTF3 | CRYBG2 |
| HS3ST1 | SLC24A4 |
| BTD | COL1A1 |
| TECR | FOXC1 |
| GRM7 | |
| NLGN1 | |
| PCDHA12 | |
| GRIK1 | |
| HS6ST3 | |
| PLXNA4 | |
| BTBD17 | |
| LMO2 | |
| EFNB1 | |
| RASEF | |
| SST | |
| SLC5A3 | |
| INHBB | |

A

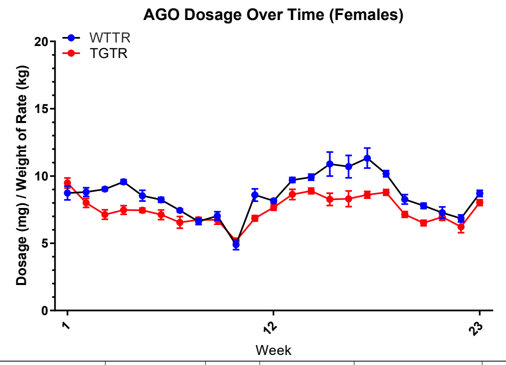

| | | | | | |
|---|---|---|---|---|---|
| Source of Variation | % of total variation | P value | P value summary | Significant? | Geisser-Greenhouse's epsilon |
| dosage over time x genotype | 8.171 | 0.0247 | * | Yes | 0.1149 |
| dosage over time | 60.28 | <0.0001 | **** | Yes | 0.1149 |
| genotype | 12.05 | 0.0041 | ** | Yes | |
| Subject | 6.100 | <0.0001 | **** | Yes | |
| ANOVA table | SS | DF | MS | F (DFn, DFd) | P value |
| dosage over time x genotype | 42.48 | 22 | 1.931 | F (2.528, 20.22) = 4.102 | P=0.0247 |
| dosage over time | 313.4 | 22 | 14.24 | F (2.528, 20.22) = 30.26 | P<0.0001 |
| genotype | 62.64 | 1 | 62.64 | F (1, 8) = 15.80 | P=0.0041 |
| Subject | 31.71 | 8 | 3.964 | F (8, 176) = 8.421 | P<0.0001 |
| Residual | 82.85 | 176 | 0.4707 | | |

B

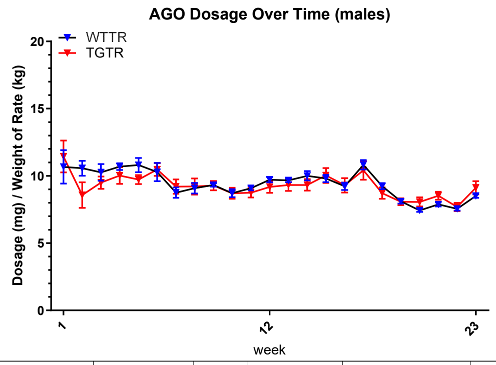

| | | | | | |
|---|---|---|---|---|---|
| Source of Variation | % of total variation | P value | P value summary | Significant? | Geisser-Greenhouse's epsilon |
| dosage over time x genotype | 4.906 | 0.2815 | ns | No | 0.1138 |
| dosage over time | 38.49 | 0.0004 | *** | Yes | 0.1138 |
| genotype | 0.2939 | 0.7773 | ns | No | |
| Subject | 27.47 | <0.0001 | **** | Yes | |
| ANOVA table | SS | DF | MS | F (DFn, DFd) | P value |
| dosage over time x genotype | 21.89 | 22 | 0.9950 | F (2.504, 20.03) = 1.363 | P=0.2815 |
| dosage over time | 171.7 | 22 | 7.807 | F (2.504, 20.03) = 10.70 | P=0.0004 |
| genotype | 1.311 | 1 | 1.311 | F (1, 8) = 0.08559 | P=0.7773 |
| Subject | 122.6 | 8 | 15.32 | F (8, 176) = 20.99 | P<0.0001 |
| Residual | 128.4 | 176 | 0.7298 | | |

C

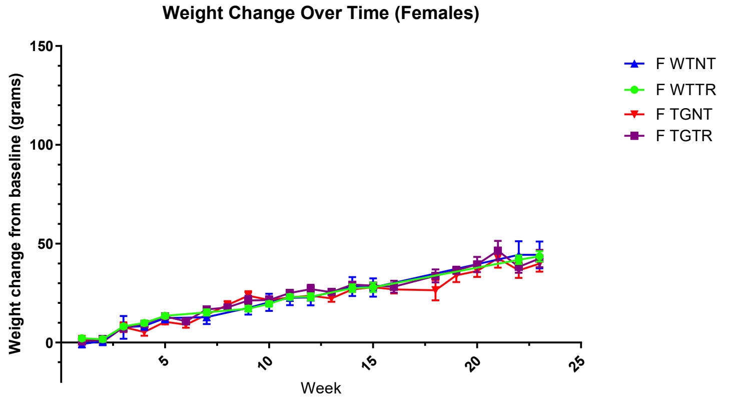

| | | | | |
|---|---|---|---|---|
| Fixed effects (type III) | P value | P value summary | Statistically significant (P < 0.05)? | F (DFn, DFd) |
| Time | <0.0001 | **** | Yes | F (3.472, 62.17) = 144.5 |
| Group | 0.9383 | ns | No | F (3, 20) = 0.1346 |

D

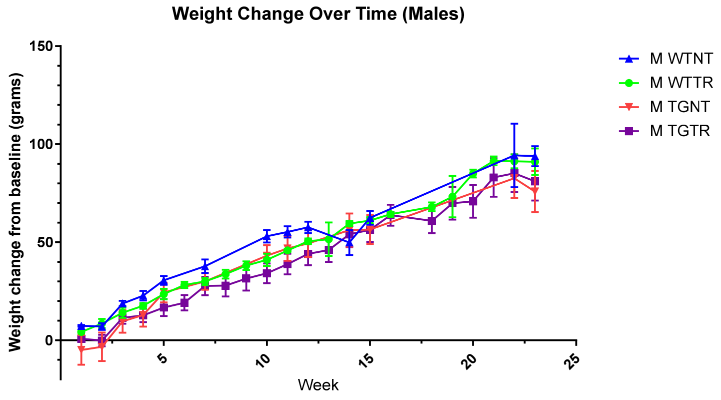

| | | | | |
|---|---|---|---|---|
| Fixed effects (type III) | P value | P value summary | Statistically significant (P < 0.05)? | F (DFn, DFd) |
| Time | <0.0001 | **** | Yes | F (4.807, 80.18) = 111.0 |
| Group | 0.2952 | ns | No | F (3, 19) = 1.326 |

### Supplemental Figure 1 – Terry et al

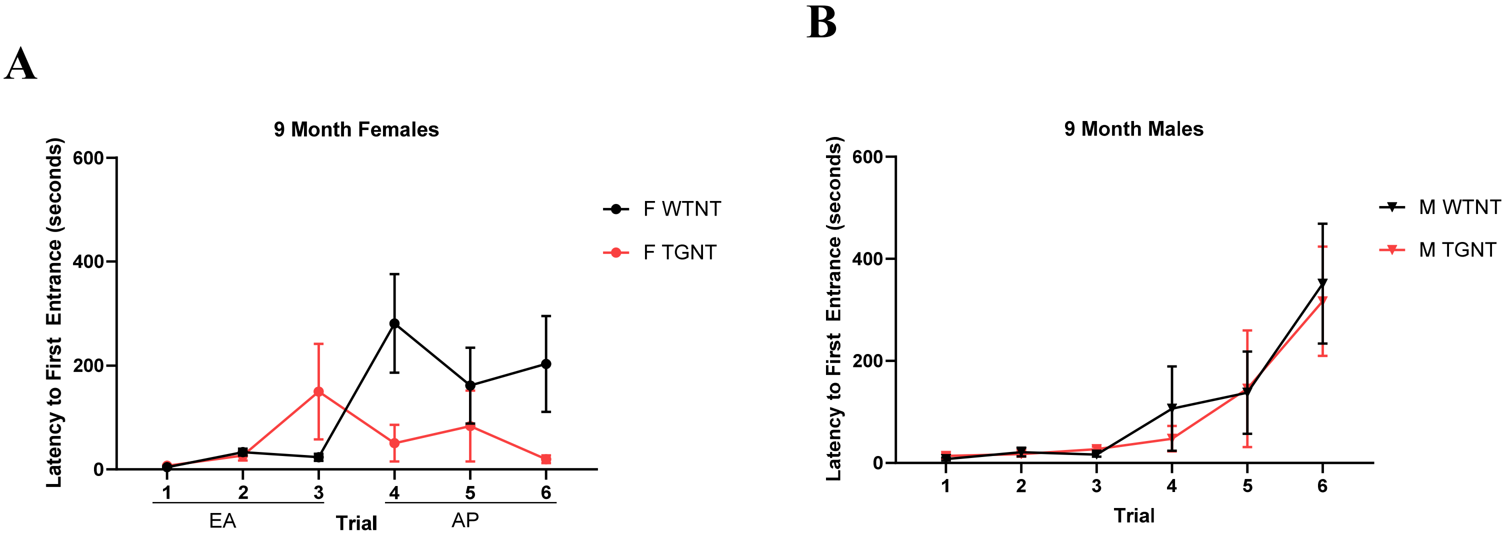

**Supplemental Figure 2 – Terry et al**
